# A standalone reverse dissimilatory sulfite reductase (rDsr) pathway enables sulfur oxidation under oxygen limitation

**DOI:** 10.1101/2024.06.14.599071

**Authors:** Xingqin Lin, Wen-Qian Zhao, Xiaoyuan Feng, Congjuan Xu, Pei-Ru Chen, Xiaojun Wang, Chunmei Deng, Hongan Long, Christian R. Voolstra, William B. Whitman, Yanlin Zhao, Peng-Fei Xia, Haiwei Luo

## Abstract

Sulfur oxidation drives nutrient and energy flow in marine sediments. Bacteria typically oxidize thiosulfate, a key sulfur intermediate, either through a complete Sox system or through an incomplete Sox system that feeds zero valent sulfur into the reverse dissimilatory sulfite reductase (rDsr) pathway. Whether rDsr alone can oxidize thiosulfate has been unclear. Here we show that ten intertidal *Ruegeria* strains carry a complete rDsr pathway but lack all *sox* genes. Under low oxygen (2% O_2_) but not under air, thiosulfate activates the pathway, enhances growth, and is associated with accelerated nitrate reduction. Using a CRISPR base editing tool for *Ruegeria*, we demonstrate that disrupting *dsrA*, which encodes a subunit of the core rDsr enzyme, significantly delays thiosulfate consumption, suggesting that rDsr is actively involved in oxidation. Previous work only documented rDsr acting downstream of an incomplete Sox system; our results establish that a standalone rDsr pathway is sufficient for thiosulfate consumption under oxygen limitation and that this activity can feed electrons into the denitrification pathway. Metagenomic surveys show rDsr genes from this group are widespread in low oxygen marine environments, including coastal sediments where they account for up to 26% of all rDsr genes. These findings revise the view that thiosulfate oxidation depends on Sox and highlight rDsr’s ecological role in deoxygenated habitats.

## Introduction

Sulfur oxidation drives key biogeochemical processes within coastal marine sediments (*1–4*). Microbial sulfate reduction mediates up to 50% of organic carbon remineralization in coastal sediments, producing substantial hydrogen sulfide (H_2_S) (*3*). Subsequently, 80–95% of this H_2_S is re-oxidized to sulfate in oxygenated surface sediments through microbial or geochemical processes involving oxygen (O_2_), iron [Fe(III)], and manganese [Mn(III, IV)] (*1, 5, 6*). Thiosulfate represents a key intermediate in this process, serving as the direct product for 68-78% of sulfide oxidation (*1, 2, 5*).

Among sulfur oxidation pathways used by sulfur-oxidizing bacteria (SOB), the Sox multi-enzyme system is the most comprehensively characterized (*7*). The complete Sox pathway, encompassing *soxXAYZBCD*, produces sulfate as its sole product (*8, 9*). However, many green sulfur bacteria (GSB) and purple sulfur bacteria (PSB) lack *soxCD* (encoding sulfur dehydrogenase), leading to an incomplete Sox pathway that accumulates zero-valent sulfur, which is subsequently oxidized through the reverse dissimilatory sulfite reductase (rDsr) pathway (*7*). Certain GSB members, such as *Chlorobium luteolum* DSM 273, lack the complete Sox pathway but possess rDsr (*10*). Yet GSB are strictly anaerobic phototrophs, and the function of rDsr in these organisms and their capacity to oxidize thiosulfate remain unexplored. More importantly, a standalone rDsr pathway (without any Sox genes) has not been reported in an aerobic bacterium.

In intertidal sediments, the Roseobacter group (Rhodobacterales, Roseobacteraceae) constitutes up to 10% of the total bacterial community (*11*). The presence of the Sox pathway in most known Roseobacter lineages from various marine habitats (*12*) and in almost all Roseobacter clusters abundant in the pelagic ocean (*13*) highlights a potential role of Roseobacters in sulfur oxidation. However, the rDsr pathway is largely absent from known Roseobacters. The only exception comes from Lenk et al. (*11*), who reported a fosmid fragment of an uncultivated Roseobacter encoding both Sox (with SoxCD) and rDsr. rDsr-exclusive Roseobacters and other aerobic bacteria have yet to be discovered.

This study reports the successful cultivation of ten new strains from intertidal marine sediments that constitute two closely related species within the Roseobacter group. These species possess the complete rDsr pathway for sulfur oxidation but lack the entire Sox pathway. Nevertheless, they efficiently oxidize thiosulfate. This lineage belongs to the genus *Ruegeria* and is evolutionarily related to the pelagic marine bacterium *Ruegeria pomeroyi* DSS-3, which encodes Sox but not rDsr (*14*). Our physiological assays, sulfur and nitrogen intermediate measurements, and transcriptomic analyses reveal that in this lineage thiosulfate oxidation may involve the standalone rDsr and is associated with denitrification under low-oxygen conditions. To establish causality, we adapted a CRISPR-guided cytosine base-editing system for *Ruegeria*, the first such tool for this genus, and generated targeted deletion mutants. Disruption of *dsrA*, encoding a subunit of the rDsr enzyme, delayed thiosulfate oxidation, suggesting that rDsr is actively involved in this process, and alternative routes might compensate for this preferred path in the mutated strain. Analysis of publicly available marine metagenomic datasets shows that rDsr of Roseobacters and marine bacterial communities is predominantly distributed in low-oxygen environments. Rhodobacterales rDsr sequences, represented by the *Ruegeria* rDsr studied here, account for up to 26% or a substantial fraction of rDsr genes found in intertidal marine sediments at a global scale.

## Results

### Cultivation and genomic characterization of an intertidal *Ruegeria* lineage encoding rDsr but not Sox

Ten bacterial strains were isolated from intertidal surface sediments (Table S1). Their 16S rRNA gene sequences shared 97.4 ±0.19% identity with that of the pelagic strain *R. pomeroyi* DSS-3. Whole-genome average nucleotide identity (ANI) between the isolates and *R. pomeroyi* DSS-3 was 81.30 ±0.15%. Phylogenomic analysis based on 2,716 single-copy orthologous genes resolved the ten strains into two distinct clades (I and II) (Fig. 1A). Members of each clade shared identical 16S rRNA gene sequences. The 16S rRNA gene identity between clades was 99.38%, whereas the maximum ANI between clades was 92.07% (Fig. 1A), below the 95% species boundary (*15*). Population structure analysis using PopCOGenT (*16*) indicated that the two clades represent separate populations with limited recent gene flow. Thus, they appear to represent two closely related species that are also related to *R. pomeroyi*.

**Figure 1.**
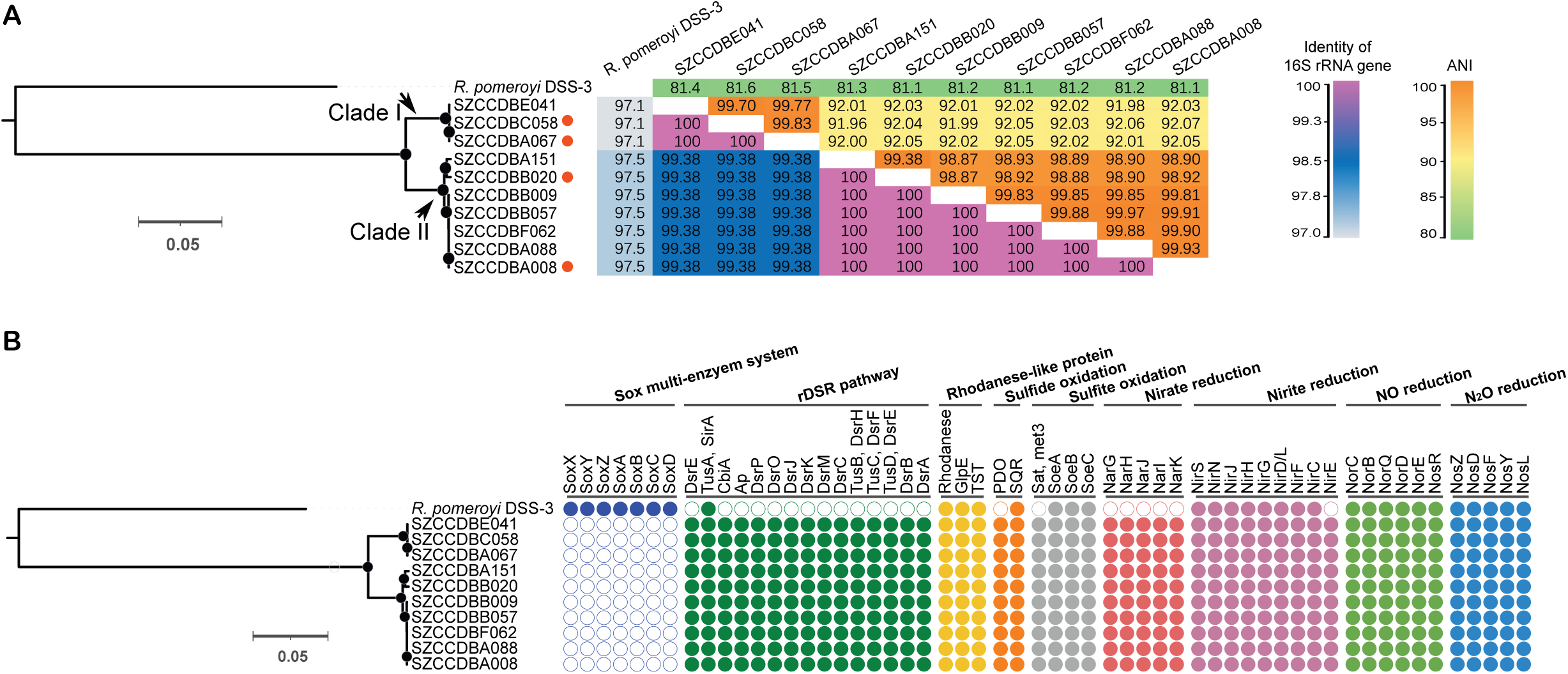
Phylogenomic tree of the intertidal *Ruegeria* lineage and the phyletic pattern of genes involved in sulfur oxidation and denitrification. **(A)** The maximum-likelihood phylogenomic tree based on the concatenation of single-copy genes shared by all strains (left) and the heatmap of the whole-genome average nucleotide identity (ANI) and the pairwise identity of 16S rRNA genes (right). Solid circles in the phylogenomic tree indicate nodes with bootstrap support of 100%. Red circles indicate strains with closed genomes. (**B**) All strains of the intertidal *Ruegeria* lineage shared identical genes involved in sulfur oxidation and denitrification. Filled and open circles indicate presence and absence of genes, respectively.

The genomes of all ten strains completely lacked the Sox pathway genes (*soxXAYZBCD*) but contained a complete reverse dissimilatory sulfite reductase (rDsr) pathway (*dsrABEFHCMKLJOP*-*dsrN*-*tusA-dsrE2*) (Fig. 1B). The absence of *sox* genes was confirmed by sequencing closed genomes of four representative strains (SZCCDBA067, SZCCDBC058, SZCCDBB020, SZCCDBA008) (Table S1). The rDsr proteins showed high sequence similarity to those of the PSB *Allochromatium vinosum* (67% for DsrA and 73% for DsrB at the amino acid level). Additional sulfur oxidation genes present included a *sqr*-*pdo* gene cluster (sulfide:quinone oxidoreductase and persulfide dioxygenase), an extra *sqr* copy distant from the cluster, several rhodanese-like sulfur-transferase genes, and the *soeABC* operon for sulfite oxidation. The adenosine-5′-phosphosulfate (APS) pathway genes *aprBA* were absent, although *sat* (ATP sulfurylase) was present.

### Thiosulfate oxidation in Sox-free yet rDsr standalone *Ruegeria* isolates

To determine whether a standalone rDsr pathway can perform thiosulfate oxidation, we cultivated the isolated strains with different sulfur sources. These sox-free isolates efficiently consumed thiosulfate, especially under microoxic conditions with 2% initial O2 in the headspace (v/v) (Fig. 2A). Thiosulfate consumption under normoxic conditions (21% oxygen) was less efficient than that under microoxic conditions (Figs. 2A and 2B). Under microoxic conditions, approximately 10 mM thiosulfate was consumed within 16 h (Fig. 2A), whereas under normoxic conditions, about 6.6 mM was consumed (Fig. 2B). Thiosulfate remained stable without inoculation at all tested conditions (Fig S1). Under these conditions, thiosulfate could serve as the sole sulfur source to sustain cell growth, and even under the sub-optimal normoxic condition, thiosulfate supported cell growth with up to 20 mM acetate, as observed in strain SZCCDBA067 and other strains of this lineage (Figs. 2A, 2B, and S2). Assuming that the sulfur content of these cells was 1% of the dry weight, a value typical of bacteria (*17*), thiosulfate consumption for this purpose would be on the order of 0.01 mM. During thiosulfate consumption, less than 1 mM of sulfate and less than 1 mM sulfite were formed under both normoxic and microoxic conditions (Figs. 2A and 2B). Tetrathionate accumulated to 3.8 mM under microoxic conditions (Fig. 2A) and to 2.3 mM under normoxic conditions (Fig. 2B). Thus, the measured extracellular sulfur species formed did not account for the total thiosulfate consumed, particularly under microoxic conditions. In addition, thiosulfate consumption also occurred in the presence of sulfate, yet with limited impacts on cell growth and slightly slower oxidation rates (Fig. 2C and 2D). These data strongly indicated that a portion of the thiosulfate was oxidized in the absence of the Sox pathway.

**Figure 2.**
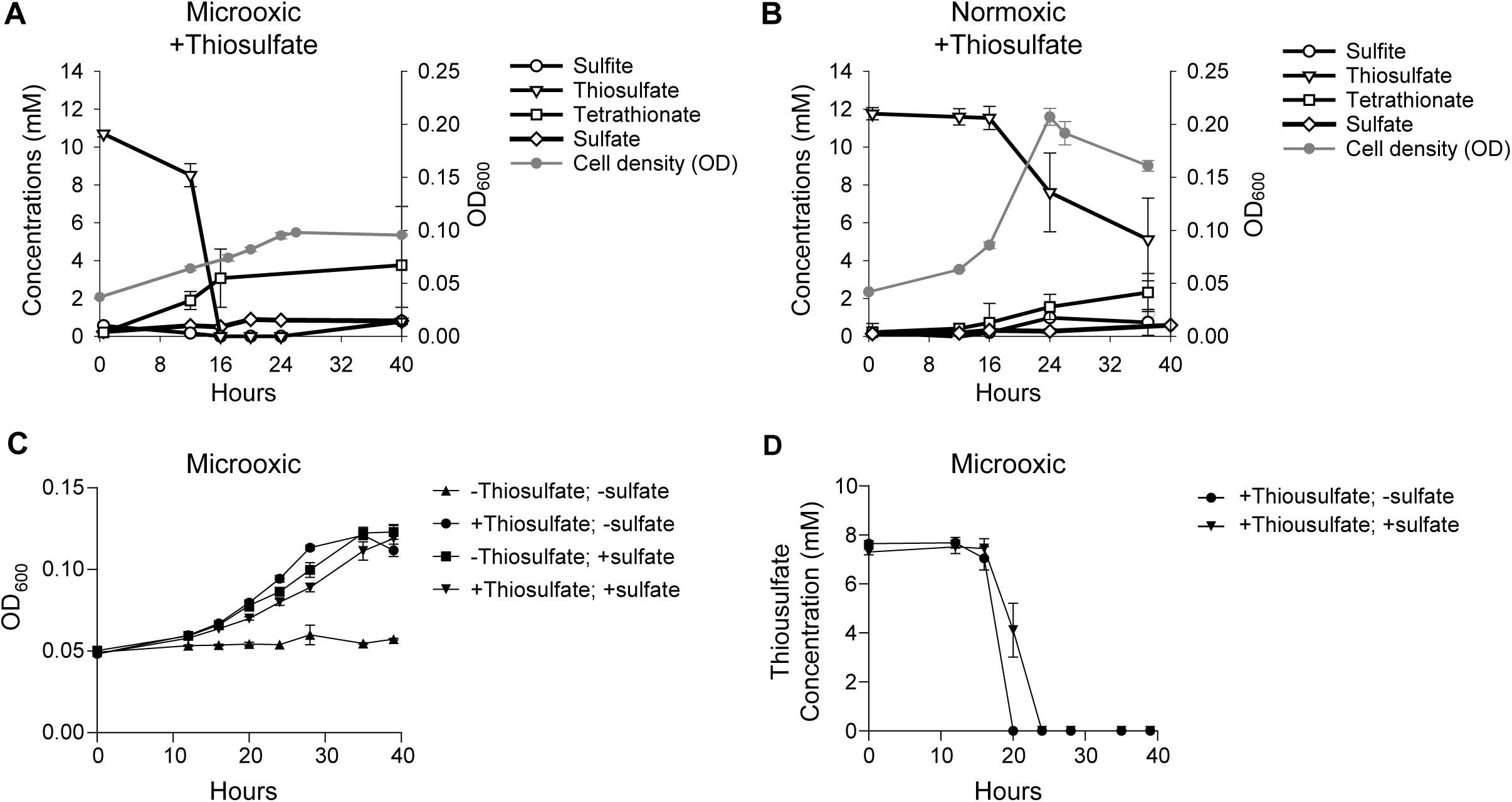
Growth and sulfur intermediate profiles of strain SZCCDBA067. **(A and B)** Time-course of bacterial growth (measured as OD_600_) and concentrations of sulfur intermediates (thiosulfate, tetrathionate, sulfite, and sulfate) under (**A**) microoxic and (**B**) normoxic conditions, with thiosulfate as the sole sulfur source. **(C)** Growth of strain SZCCDBA067 and (**D**) the concurrent thiosulfate consumption under microoxic conditions using thiosulfate, sulfate, or both as sulfur sources. Error bars indicate the standard deviation from three biological replicates.

### Transcriptional activation of the rDsr pathway in response to thiosulfate under oxygen limitation

To determine if the rDsr pathway was involved in thiosulfate consumption and further explore the mechanism underlying potential thiosulfate oxidation, we first confirmed that strain SZCCDBA067 could grow well on sulfate as the sole sulfur source under both normoxic and microoxic conditions (Figs. 2C and S3). Using sulfate as a reference, we then compared the transcriptional responses of SZCCDBA067 to thiosulfate addition under normoxic and microoxic conditions (Fig. 3). In strain SZCCDBA067, under normoxic sampling conditions used, thiosulfate addition did not detectably alter transcript abundances of sulfur oxidation genes (Fig. 3A and Table S2). Under microoxic conditions, all 16 genes in the rDsr pathway (*dsrABEFHCMKLJOP*-*dsrN*-*tusA-dsrE2*) showed significantly higher transcript levels in the presence of thiosulfate. The largest increases were observed for *dsrAB* (53-fold and 21-fold, respectively) and *dsrEFH* (45-fold, 22-fold, and 26-fold, respectively) (Fig. 3A and Table S2). Transcripts of *soeABC* and one copy of *sqr* (unlinked to the *sqr*-*pdo* cluster) were also enriched under microoxic conditions with thiosulfate (Fig. 3A). The *sqr*-*pdo* cluster did not show differential expression. Among seven rhodanese-encoding genes, only one (SZCCDBA067_01274), containing the conserved CASGXR motif (Fig. S4), was significantly upregulated in response to thiosulfate under microoxic conditions (Fig. 3A). The *dsrJ* transcript increased 6.2-fold under the same condition. Genes involved in assimilatory sulfate reduction (*cysH*, *cysI*) showed decreased transcript levels when thiosulfate was present, especially under microoxic conditions (Fig. 3A). In a sister strain (SZCCDBB020), rDsr pathway genes, *soeABC*, and the CASGXR-containing rhodanese were similarly upregulated in response to thiosulfate under microoxic but not normoxic conditions (Fig. S5A).

**Figure 3.**
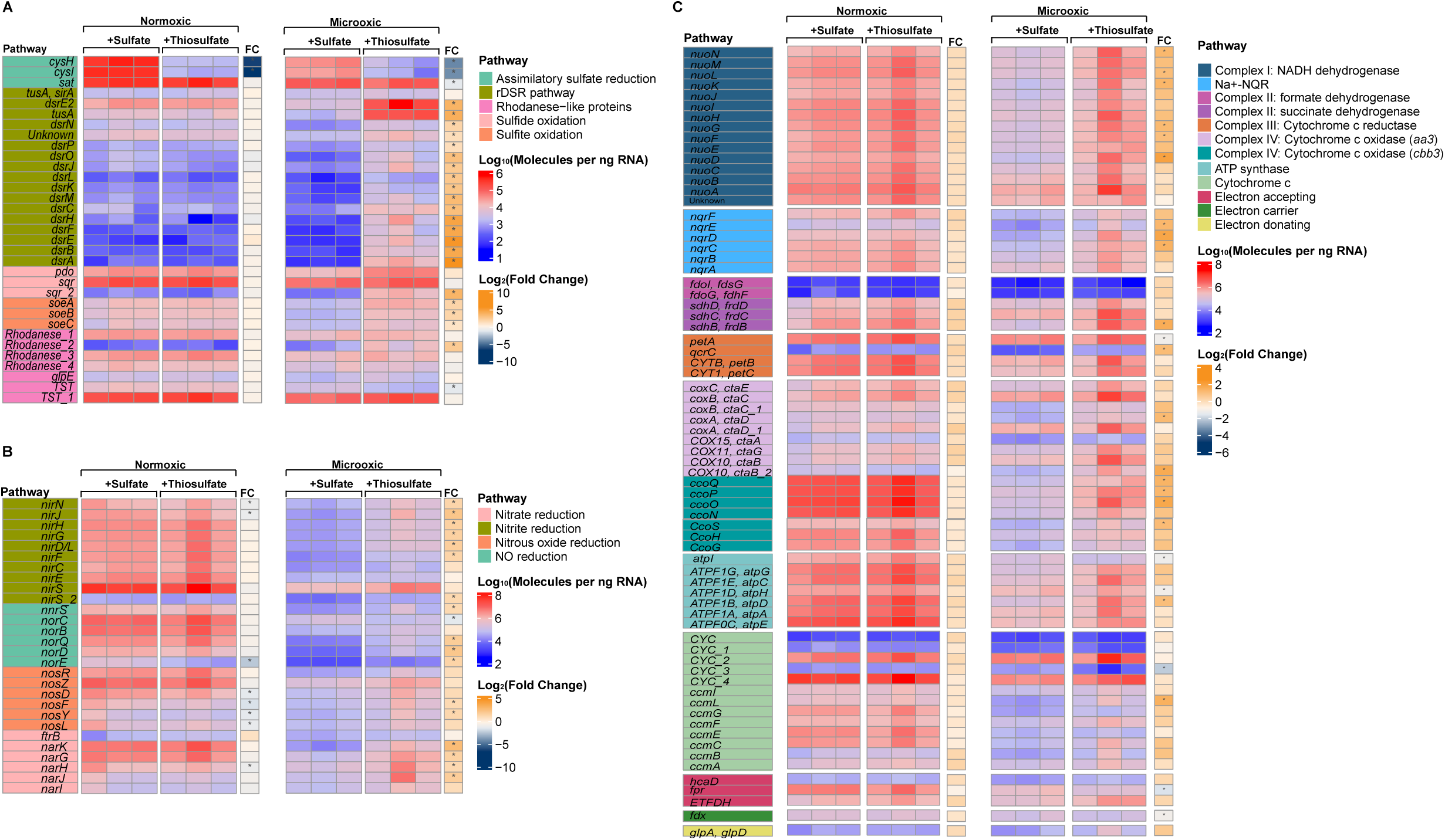
Transcript abundances of genes involved in (A) inorganic sulfur metabolism, (B) denitrification, and (C) aerobic respiration. The strain SZCCDBA067 was cultured in the presence or absence of thiosulfate, and transcript abundances were measured in triplicates by normalizing respective transcript sequencing reads to the standard curve of external spike-in RNA controls. The color bars show the transcript copies per nanogram total RNA, with red for higher and blue for lower levels. The bars labeled as FC represent the log2 fold change of transcript abundances in cultures with thiosulfate compared to those without. Stars indicate log2 fold change >1 and adjusted *p* values < 0.01. See Table S2, S3, and S4 for the abbreviations of (**A**) inorganic sulfur metabolism, (**B**) denitrification and (**C**) aerobic respiration genes.

### Establishment of a CRISPR-guided cytosine base-editing system for *Ruegeria*

A CRISPR-guided cytosine base-editing system was adapted for *Ruegeria*. The system comprises a nuclease-deactivated Cas9 (dCas9) fused to a cytidine deaminase (PmCDA1) and a uracil DNA glycosylase inhibitor (UGI) (Fig. S6). The dCas9 binds a target DNA site adjacent to an NGG protospacer-adjacent motif (PAM) without cleaving the DNA. The PmCDA1 converts cytidine to uridine within a defined editing window, and the UGI prevents removal of the uridine, thereby achieving a C-to-T change upon DNA replication or repair (Figs. 4A and S6) (*18, 19*). Expression of the editor is driven by an inducible *lacI*-P_trc_ module, and the single-guide RNA (gRNA) cassette is constitutively expressed with the target cytidine positioned within the deamination window (Fig. S6). After electroporation of the editing plasmid, induction with IPTG enables base editing, followed by colony screening for successful edits (Fig. S7). To eliminate potential effects of the editing plasmid on cell physiology, the plasmid was cured from edited strains by passage in antibiotic-free medium and screening for loss of antibiotic resistance (Fig. S8).

**Figure 4.**
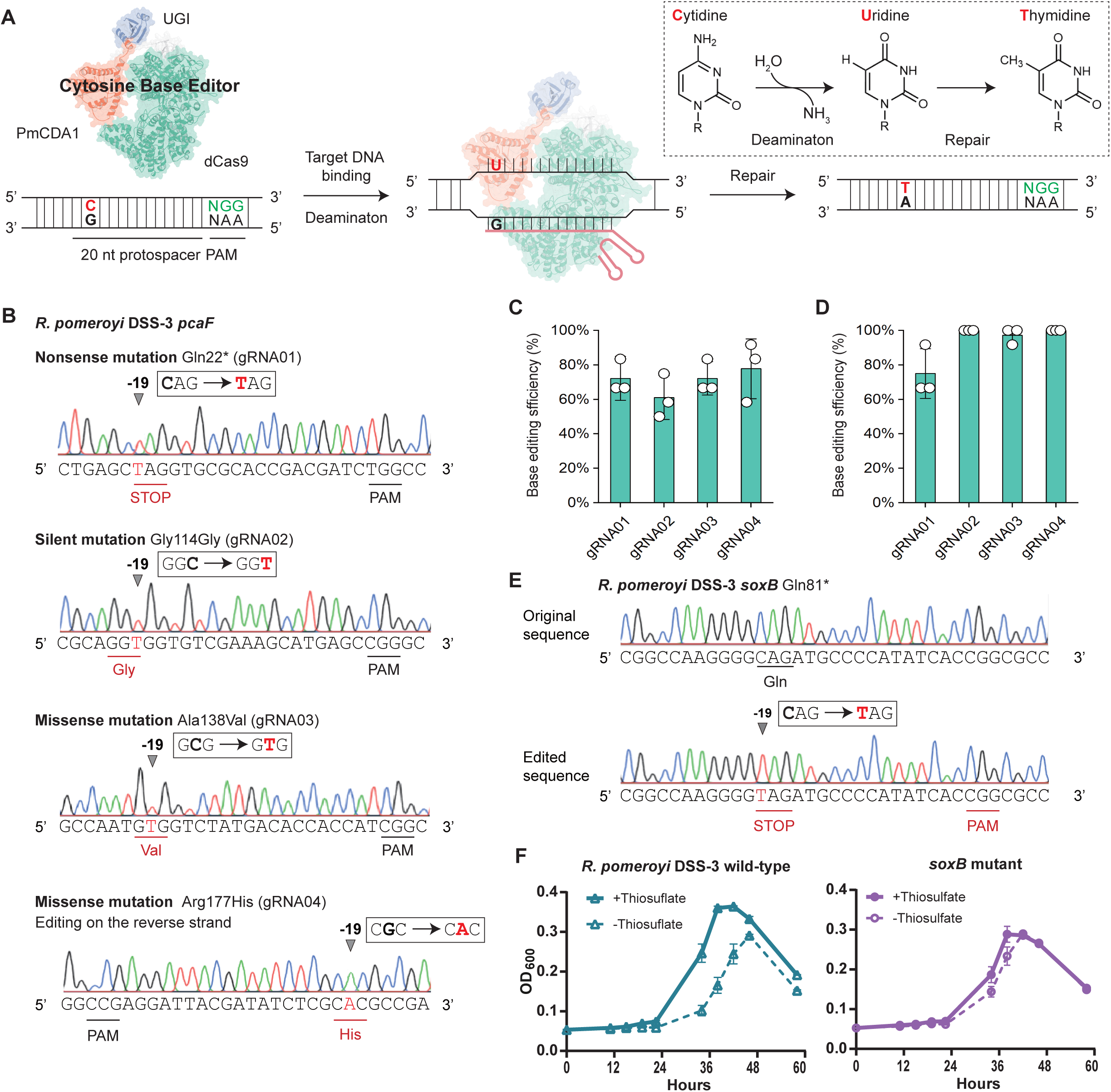
Establishment and validation of a CRISPR-guided cytosine base-editing platform in *R. pomeroyi* DSS-3. (**A**) Schematic of the cytosine base editor mechanism, showing the dCas9-PmCDA1-UGI complex binding to target DNA and catalyzing C-to-T conversion. (**B**) Representative Sanger sequencing chromatograms of four edit types generated at pcaF: nonsense (Gln22*), silent (Gly114Gly), missense (Ala138Val), and reverse-strand missense (Arg177His) mutations. (**C**) Editing efficiencies of each gRNA after 24 h IPTG induction. (**D**) Editing efficiencies after 48 h cumulative IPTG induction. (**E**) Sanger sequencing confirmation of *soxB* Gln81* nonsense mutation. (**F**) Growth profile of the wild-type and *soxB* mutant strains in media containing 10 mM acetate and 20 mM sulfate, with or without thiosulfate, under normoxic conditions. Error bars indicate the standard deviation from three biological replicates.

In the model strain *R. pomeroyi* DSS-3, the system was tested at the non-essential *pcaF* locus. Four different edit types were generated: nonsense, silent, missense, and reverse-strand missense (Fig. 4B). After 24 h induction with 0.1 mM IPTG, editing efficiencies reached 61-78% (Fig. 4C). Following an additional 24 h passage under induction, efficiencies increased to 100% for three of the four edits (Fig. 4D). The system was also used to introduce a premature stop codon into *soxB* (encoding a canonical sulfur-oxidation enzyme) in *R. pomeroyi* DSS-3, generating a Sox-deficient strain for future comparisons (Fig. 4E). While the wild-type DSS-3 showed increased growth with thiosulfate as previously reported (*14*), the Sox-deficient strain showed similar growth profiles with or without thiosulfate (Fig. 4F), indicating a desired phenotypical change of the edited strain.

### Genetic validation of rDsr function in thiosulfate oxidation

To genetically validate the role of the rDsr pathway and the candidate rhodanese in thiosulfate consumption, early stop codons were introduced into *dsrA* and the CASGXR-containing rhodanese gene (*rhd*, SZCCDBA067_01274) in the representative isolate SZCCDBA067 using the base-editing system (Fig. 5A). The wild-type strain, and the mutant strains (Δ*dsrA* and Δ*rhd*) were incubated under microoxic conditions (2% initial O_2_ in the headspace) with thiosulfate (Figs. 5B and 5C). Disruption of *dsrA* resulted in slower growth compared to the wild-type strain (*μ* = 0.043 ±0.003 h^-1^ vs 0.059 ±0.001 h^-1^, Student’s t-test, *P* < 0.001). For thiosulfate consumption, the concentration decreased over time in wild-type cultures (Fig. 5C). Inactivation of *dsrA* delayed thiosulfate consumption. These results indicated that the rDsr pathway played an important role in thiosulfate utilization.

**Figure 5.**
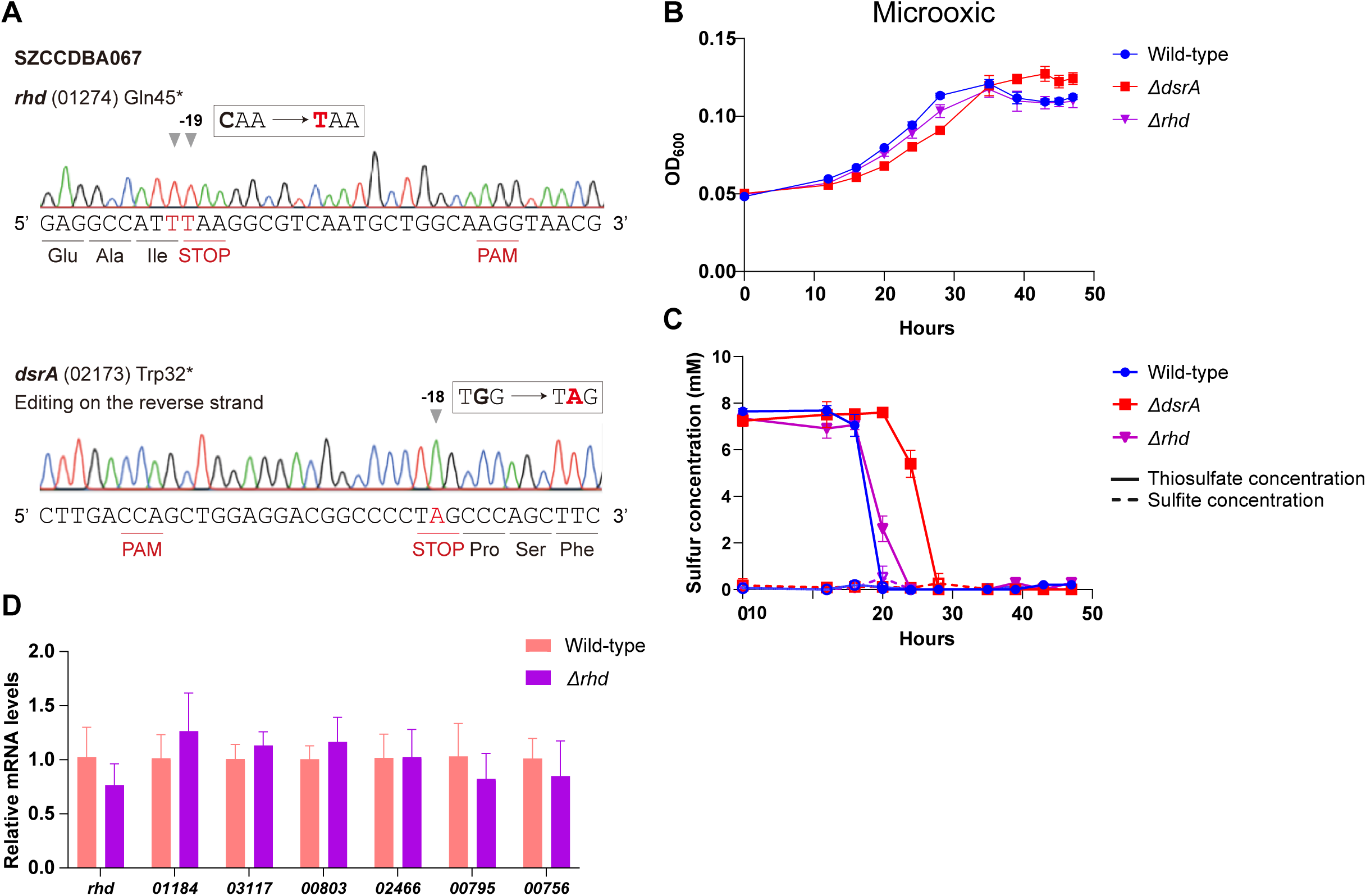
Growth, thiosulfate consumption, and gene expression analysis of Δ*dsrA* and Δ*rhd* mutants using thiosulfate as the sole sulfur source. **(A)** Sanger sequencing alignments validating the CBE-mediated C-to-T conversions, which successfully generated Gln45* and Trp32* nonsense mutations in *rhd* and *dsrA*, respectively. (**B**) Growth profile of the wild-type strain A067, Δ*dsrA*, and Δ*rhd* under microoxic conditions (2% O₂) in medium supplemented with thiosulfate as the sole sulfur source. (**C**) Time-course of thiosulfate and sulfite consumption in the concurrent cultures. (**D**) Relative mRNA levels of *rhd* and six other putative rhodanese genes (gene locus indicated on the x-axis) in A067 and the Δ*rhd* mutant, determined by RT-qPCR. Note that no significant differences were observed between the two strains for any of the genes tested (*P* > 0.05). Error bars represent the standard deviation from three biological replicates.

Disruption of *rhd* did not significantly alter the growth rate (*μ* = 0.058 ±0.001 h^-1^, *P* > 0.05, Fig. 5B). Moreover, the Δ*rhd* mutant showed consumption rates similar to A067 (Fig. 5C), whereas *rhd* is replaceable for this process under the tested conditions. Given that *rhd* is one of multiple copies of rhodanese genes in the genome, the thiosulfate oxidation in the Δ*rhd* mutant under the tested conditions is likely due to functional redundancy with other rhodanese-like proteins. We examined the expression of seven rhodanese genes and found that their mRNA levels were not significantly altered in the Δ*rhd* mutant compared to the wild type (Fig. 5D). Specifically, the *rhd* gene was downregulated to 0.77-fold of the wild-type level in the mutant, whereas genes SZCCDBA067_01184, SZCCDBA067_03117, and SZCCDBA067_00803 showed slight upregulation; however, none of these changes were statistically significant (Fig. 5D). Together, these expression data suggest either a saturated activity of the rhodanese from all copies or functional redundancy at the post-transcriptional level.

### Association of thiosulfate oxidation with denitrification under microoxic conditions

The intertidal *Ruegeria* lineage encodes a complete denitrification pathway (Fig. 1B). Under microoxic conditions, transcript levels of denitrification genes were significantly higher in cultures with thiosulfate than without (Fig. 3B and Table S3), including *narK* (a transporter for nitrate and nitrite), *narGHJ* (the respiratory nitrate reductase), *nirFDGHJN* (heme *d*_1_ biosynthesis proteins), *norDEQ* (nitric oxide reductase activation proteins), and *nosFY* (nitrous oxide reductase maturation proteins). Nitrate was also reduced more rapidly during growth in the presence of thiosulfate, a 99% decrease within 16 h, than in its absence, 61% decrease within 16 h, under microoxic conditions (Fig. S9). Nitrite accumulated in cultures without thiosulfate but was rapidly depleted when thiosulfate was present (Fig. S9). Under microoxic conditions, transcripts of genes encoding NADH dehydrogenase (Complex I and Na^+^-pumping NADH:quinone oxidoreductase) and Complex III were enriched in thiosulfate-supplemented cultures (Fig. 3C, Table S4).

Under normoxic conditions, transcripts of denitrification genes were higher than under microoxic conditions regardless of thiosulfate addition (Fig. S10). Transcripts of the *cbb₃*-type cytochrome c oxidase genes (*ccoNOPQ*) and *ctaB* (heme O synthase) were also higher under normoxic than under microoxic conditions (Fig. 3C). However, nitrate reduction rates under normoxic conditions were lower than under microoxic conditions (Fig. S9), and transcripts of assimilatory nitrate reduction genes (*nirB*, *nrtABC*, *nasST*) were less abundant under normoxic than under microoxic conditions (Fig. S10). RT-qPCR validated the elevated expression of *narG*, *nirS*, *nosZ*, *ccoN*, and *ctaB* under normoxic conditions compared to microoxic conditions, consistent with the RNA-seq data (Fig. S11). Ammonium supported faster growth and higher yields than nitrate as the sole nitrogen source (Fig. S12).

### Environmental distribution of rDsr in low-oxygen marine niches

Phylogenetic analyses confirmed that Rhodobacterales-associated rDsr sequences (DsrA, DsrB) belong to the oxidative type and form a distinct clade (Figs. S13 and S14). In *Tara* Oceans metagenomes (*20*) (Table S5), copy numbers of Rhodobacterales-associated *dsrA* (Fig. 6A) and *dsrB* (Fig. S15A) per genome were significantly higher in low-oxygen (1–5 µM O_2_) and very-low-oxygen (< 1 µM O_2_) waters than in oxic (> 60 µM O_2_) or moderately oxygenated (5–60 µM O_2_) waters (Wilcox test, *P* < 0.001). The same pattern was observed for total bacterial community-associated *dsrA* and *dsrB* (Figs. 6B & S15B). In contrast, copy numbers of *soxA* and *soxB* per genome were higher in oxygenated than low-oxygen environments (Fig. S16).

**Figure 6.**
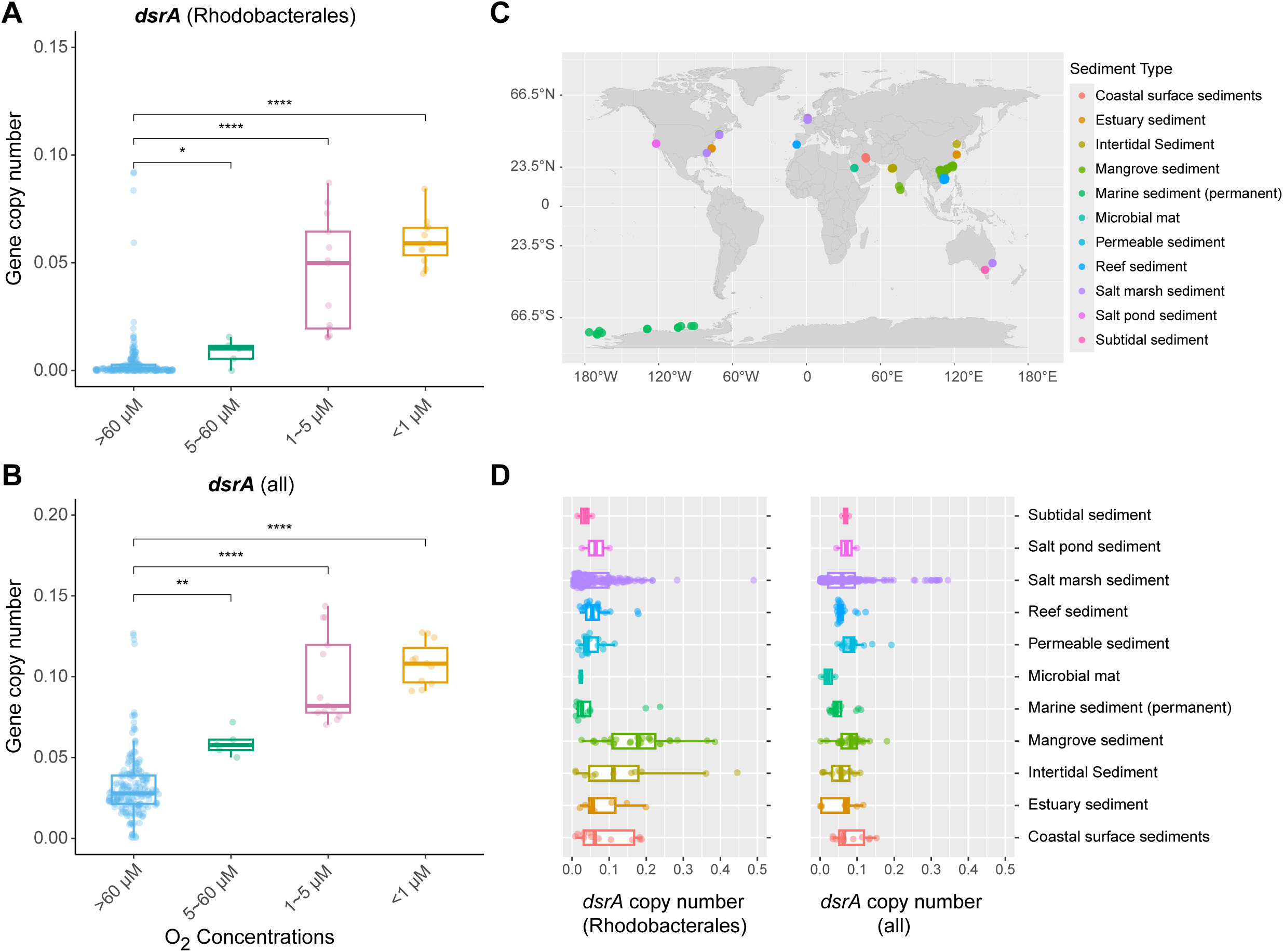
Copy numbers of *dsrA* in marine metagenomes. (**A and B)** Copy number (per cell) of *dsrA* among *Tara* Oceans metagenomes sampled from sites with different oxygen levels. The copy number was estimated based on the number of reads assigned to the oxidative bacterial-type DsrA in the reference tree, normalized by the average number of reads mapped onto 21 single-copy marker genes and by their respective gene length. **(A)** Copy numbers of Rhodobacterales-associated *dsrA* and **(B)** of *dsrA* at the bacterial community level are represented by dots. The median is represented by the center line of the box, while the upper and lower quartiles are indicated by the box limits. The whiskers show 1.5 times the interquartile range. The *Tara* Oceans samples were categorized into four groups based on the oxygen concentration at the sampling site: oxic (> 60 µM), microoxic (> 5 µM but < 60 µM), suboxic (> 1 µM but < 5 µM) and nanooxic (< 1 µM). The stars denote a *p*-value of less than 0.001 (Wilcox test). ns: not significant (*P* > 0.05). (**C** and **D)** Copy number of *dsrA* genes in coastal sediment metagenomic samples and their geographic distribution. **(C)**The distribution of different sediment types is shown on the map. **(D)** Copy numbers of Rhodobacterales-associated *dsrA* and of *dsrA* at the community level are represented by dots. Dots are color-coded based on sediment type.

In 305 coastal sediment metagenomes (salt marshes, reefs, mangroves, tidal flats) worldwide (Fig. 6C and Table S6), oxidative *dsrAB* genes were prevalent (Figs. 6D and S15D). Copy numbers of Rhodobacterales-associated *dsrA* per genome ranged from 0.0026 to 0.4898 (median 0.0538) in coastal sediments, compared to 0.0001 to 0.0920 (median 0.0022) in pelagic waters (Fig. S17A), while those of total bacterial community-associated *dsrA* ranged from 0.0004–0.3452 (median 0.0672) in coastal sediments versus 0.0081–0.1437 (median 0.0334) in pelagic waters (Fig. S17B). For *dsrB*, copy numbers of Rhodobacterales-associated sequences ranged from 0.0044 to 0.3737 (median 0.0629) in coastal sediments, compared to 0.0002 to 0.0677 (median 0.0024) in pelagic waters (Fig. S17C), and those of total bacterial community sequences ranged from 0.0004–0.2421 (median 0.0555) in coastal sediments versus 0.0054–0.1176 (median 0.0228) in pelagic waters (Fig. S17D). Rhodobacterales-associated *dsrA* genes were detected in 173 of the 305 sediment metagenomes, accounting for 0.16–20.45% of total *dsrA* (median = 3.31%, mean = 3.89%; Table S6). For *dsrB*, Rhodobacterales-associated genes were detected in 167 out of the 305 sediment metagenomes, accounting for 0.92–26.37% of total *dsrB* (median 4.30%, mean 5.28%; Table S6). Thus, Rhodobacterales-associated rDsr sequences represent up to 26% of the total rDsr pool.

## Discussion

The reverse dissimilatory sulfite reductase (rDsr) pathway is thermodynamically more efficient for sulfur oxidation than the canonical Sox system (*10, 21*). Whether bacteria can use rDsr alone to oxidize thiosulfate has remained unknown. In most known sulfur-oxidizing bacteria, rDsr acts downstream of an incomplete Sox pathway that produces zero-valent sulfur (*7*). A standalone rDsr pathway (without any *sox* genes) has never been reported in an aerobic bacterium. Here we show that a previously uncultivated lineage of intertidal *Ruegeria* possesses a complete rDsr pathway in the complete absence of *sox* genes, and that this standalone pathway participates in thiosulfate consumption under oxygen limitation. Presumably, it participates in thiosulfate oxidation, activities associated with denitrification that are widespread in deoxygenated marine environments.

All 16 genes of the rDsr pathway, including *dsrAB* (encoding the siroheme sulfite reductase) and *dsrEFH* (sulfur transfer proteins), were strongly induced by thiosulfate specifically under microoxic conditions (Fig. 3 and Table S2). Transcripts of *soeABC* (sulfite oxidation) and a single rhodanese gene, designated *rhd* (locus tag SZCCDBA067_01274), containing the conserved CASGXR motif, were also enriched under the same condition. This transcriptional response linked thiosulfate addition to rDsr pathway activation only when oxygen was limited.

Disruption of *dsrA*, encoding a core subunit of the rDsr enzyme, delayed thiosulfate consumption in the isolate SZCCDBA067 under microoxic conditions (Fig. 5). The Δ*dsrA* mutant retained partial thiosulfate oxidation activity, suggesting that alternative, as-yet-unidentified routes may compensate for rDsr function in this sox-free host.

Thiosulfate entry into the rDsr pathway in the absence of Sox requires an alternative mechanism. In the purple sulfur bacterium *Allochromatium vinosum*, which has an incomplete Sox pathway lacking *soxCD*, thiosulfate is first processed by SoxXA and SoxB to produce zero-valent sulfur that is then oxidized by rDsr (*7*). The *Ruegeria* lineage lacks all Sox genes. The upregulated *rhd* gene may transfer sulfane sulfur from thiosulfate to glutathione, forming glutathione persulfide (GSSH) and sulfite, analogous to the yeast RDL1 enzyme (*22*). GSSH could then feed into the rDsr cascade via TusA, DsrEFH, and DsrC, as demonstrated in *A. vinosum* (*23, 24*). However, direct biochemical validation of these reactions is required.

The Δ*rhd* mutant did not abolish thiosulfate oxidation; it consumed thiosulfate at rates comparable to the wild type (Fig. 5). Seven rhodanese-encoding genes are present in the genome, and transcript levels of the other six paralogs were not significantly altered in the Δ*rhd* mutant (Fig. 5). This points to functional redundancy at the post-transcriptional level or saturation of persulfide transfer activity. Compensation by alternative metabolic pathways cannot be excluded. Therefore, although *rhd* is transcriptionally responsive to thiosulfate under microoxia, it is not strictly required under the tested conditions, likely because other sulfurtransferases can functionally substitute.

Thiosulfate consumption produced tetrathionate, particularly under microoxic conditions (3.8 mM; Fig. 2A). Known thiosulfate-to-tetrathionate enzymes (TsdA, TQO, ThdT) are absent from the genomes, except a distant *thdT* homolog that was down-regulated during growth with thiosulfate. Transcripts of *dsrJ*, which encodes a triheme cytochrome c with a His/Cys coordination similar to TsdA (*25, 26*), were upregulated (Fig. 3), suggesting a possible role for the DsrMKJOP complex in tetrathionate formation. Direct testing of this hypothesis is required. The mass balance deficit for thiosulfate consumption remains unexplained; it may reflect assimilation into biomass or conversion to undetected species such as polysulfide.

The intertidal *Ruegeria* lineage carries a complete denitrification pathway, including the membrane-bound nitrate reductase NarGHJ (Fig. 1). Under microoxia, thiosulfate addition accelerated nitrate reduction and prevented nitrite accumulation (Fig. S12), coinciding with increased transcripts of denitrification genes, *dsrL* (encoding an NAD^+^ reductase), and components of the respiratory electron transport chain (NADH dehydrogenase Complex I and the bc_1_ complex Complex III) (Fig. 3). In *A. vinosum*, electrons generated by rDsrAB are transferred to NAD^+^ via the iron-sulfur flavoprotein DsrL (*27*). A similar mechanism likely operates in this *Ruegeria* lineage: electrons from thiosulfate are transferred via rDsr to NAD^+^ through DsrL, and the resulting NADH feeds into the denitrification respiratory chain. This integration of sulfur and nitrogen metabolism likely underpins the ecological success of rDsr-carrying bacteria in oxygen-limited, nitrate-rich sediments. In natural environments, other electron acceptors such as iron or manganese could potentially serve a similar role, although only nitrate was tested in pure cultures.

Sulfide, the other major reduced sulfur species in intertidal sediments (*1, 3*), was not tested in this study because rDsr cannot directly oxidize sulfide. Sulfide must first be oxidized by sulfide:quinone oxidoreductase (Sqr) to elemental sulfur or polysulfide (*28*). These products can then enter at least two downstream routes: the persulfide dioxygenase (Pdo) pathway or the rDsr pathway (*28–30*). Any observed sulfide oxidation in these isolates could therefore be attributed to Pdo, rDsr, or both. Thiosulfate, by contrast, bypasses this ambiguity because it may be directly processed by the proposed rhodanese-dependent entry mechanism. The ability of a standalone rDsr to oxidize thiosulfate had not been demonstrated previously, and testing sulfide would not have addressed this specific question.

The *Ruegeria* lineage studied here is phylogenetically close to *R. pomeroyi* DSS-3, a pelagic bacterium that uses Sox but not rDsr (*14*). Two sulfur oxidation strategies coexist in the same genus, suggesting habitat-specific environmental selection. rDsr confers a competitive advantage under oxygen limitation, whereas Sox may be favored in normoxic waters. Metagenomic analysis supports this interpretation. In *Tara* Oceans samples, Rhodobacterales-associated *dsrAB* were significantly enriched in suboxic (1–5 µM O_2_) and nanooxic (<1 µM O_2_) waters (Figs. 6 and S15), whereas *soxAB* were more abundant in oxygenated waters (Fig. S16). Total bacterial community *dsrAB* showed the same directional enrichment under low oxygen (Figs. 6 and S15). The enrichment was less pronounced for total *dsrAB* than for Rhodobacterales-associated *dsrAB*, in part because diverse bacterial lineages may contribute to the total *dsrAB* pool across all oxygen regimes, including oxic waters where Rhodobacterales contributions are minimal. This difference indicates that many bacterial lineages carry oxidative *dsrAB*, but the Rhodobacterales lineage is particularly specialized for low-oxygen niches. In 305 coastal sediment metagenomes from global sites, Rhodobacterales-associated *dsrA* and *dsrB* was detected in 57% and 55% of the samples and accounted for up to 20% and 26% of total *dsrA* and *dsrB* reads, respectively (Figs. 6 and S15; Table S6). The high contribution of Rhodobacterales in sediments, which are typically oxygen-limited, further supports their ecological role as key rDsr-driven sulfur oxidizers in deoxygenated habitats. Current surveys of sulfur-oxidizing bacteria often rely on *soxB* as a marker (*31–33*). The results presented here indicate that oxidative type *dsrAB* must be included to capture the full diversity of these communities.

Several open questions remain. The transport of thiosulfate from the periplasm into the cytoplasm, where the rDsr pathway and rhodaneses reside, is not understood. Known thiosulfate transporters (e.g., TsdA-associated or *thio* systems) are absent from the genomes. Identifying this transporter will be essential to complete the model of thiosulfate oxidation in Sox-free bacteria. Another open question is whether this lineage can oxidize thiosulfate under strict anoxia; only microoxic conditions (2% initial O_2_) were tested. The ability of this lineage to oxidize sulfide, and whether rDsr contributes to that process, remains unexplored. Biochemical validation of the proposed rhodanese-dependent thiosulfate entry mechanism and the role of DsrMKJOP in tetrathionate formation is also needed.

A *sox*-free, standalone rDsr pathway can oxidize thiosulfate under oxygen limitation. This finding revises the long-held view that thiosulfate oxidation depends on Sox enzymes. The pathway is widespread in deoxygenated marine environments, where Roseobacters, previously overlooked in this context, are major contributors to sulfur cycling.

## Methods

### Sample collection, bacterial isolation, and genome sequencing

The 10 new strains were isolated from intertidal surface sediments obtained from Deep Bay (Shenzhen, China) on January 2nd, 2019. The isolation procedure followed a previously described method (*34*). Briefly, sediments were serially diluted in sterile seawater and plated on minimum media agar containing NaCl (30 g/L), MgCl_2_·6H_2_O (4.18 g/L), MgSO_4_·7H_2_O (3.4 g/L), KCl (0.33 g/L), NH_4_Cl (0.25 g/L), CaCl_2_ (0.106 g/L), K_2_HPO_4_ (0.14 g/L), NaHCO_3_ (0.3 g/L), Fe(NH_4_)_2_(SO_4_)_2_·6H_2_O (0.01 g/L), NiCl_2_·6H_2_O (0.5 mg/L), Na_2_SeO_3_·5H_2_O (0.5 mg/L), trace minerals (10 ml/L), vitamin (10 ml/L), the Bacto agar (15 g/L), and taurine (1.25 g/L) as the sole carbon source. After incubation at 30 °C for approximately 3 days, colonies were randomly selected and purified on marine agar 2216 (Difco^TM^).

For each purified strain, biomass was collected and genomic DNA was extracted using the EZNA Bacterial DNA Kit (D3350-02, OMEGA Bio-Tek). DNA libraries were constructed with the NEBNext Ultra II FS DNA Library Prep Kit for Illumina and sequenced on an Illumina HiSeq PE150 (2×150 bp paired-end reads, average insert size 350 bp) at Guangdong Magigene Biotechnology Co. Ltd.

### Genome assembly, annotation, and phylogenomic analysis

Raw Illumina reads were trimmed with Trimmomatic v0.36 (*35*) using options ‘SLIDINGWINDOW:4:15 MAXINFO:40:0.9 MINLEN:40’, and quality was checked with FastQC v0.11.5 (http://www.bioinformatics.babraham.ac.uk/projects/fastqc). Clean reads were assembled *de novo* using SPAdes v3.10.1 (*36*) with default parameters. Contigs shorter than 1,000 bp with a sequencing depth below 5×were removed. Genome completeness, contamination, and strain heterogeneity were assessed with CheckM v1.0.7 (*37*). Protein-coding genes were predicted with Prokka v1.12 (*38*) and annotated using the RAST server v2.0 (*39*), KEGG v82.0 (*40*), and CDD v3.16 (*41*).

To confirm the absence of the *sox* genes, four representative strains (SZCCDBA067, SZCCDBC058, SZCCDBB020, SZCCDBA008) were additionally sequenced on a Nanopore sequencer (Oxford Nanopore Technologies). Raw reads were polished with NECAT v0.0.1 (*42*) and assembled with Flye v2.6 (*43*). The assembly was polished twice with racon v1.4.13 (*44*) and four times with pilon v1.23 (*45*) using Illumina paired-end reads, and circularization of both chromosomes and plasmids was confirmed with Bandage v0.8.1 (*46*), resulting in closed genomes.

For phylogenomic analysis, 2,716 single-copy orthologous gene families were identified with OrthoFinder 2.2.7 (*47*). Each family was aligned at the amino acid level using MAFFT v7.125 (*48*) and trimmed with TrimAl v1.4.rev15 (*49*). The concatenated alignment was used to construct a maximum likelihood tree with IQ-TREE 1.6.8 (*50*) and 1,000 bootstrap replicates (*51*). Pairwise 16S rRNA gene identity was calculated with BLASTN (*52*), and whole-genome average nucleotide identity (ANI) was computed with fastANI (*15*).

### Growth assay on thiosulfate and nitrate

Two representative strains from Clade I (SZCCDBA067 and SZCCDBC058) and two from Clade II (SZCCDBB020 and SZCCDBA008) were grown in minimum medium with 20 mM acetate as the sole carbon source and 20 mM nitrate as an additional electron acceptor. Thiosulfate (10 mM) was added where indicated; control cultures received 20 mM sulfate. Normoxic conditions (21% initial O_2_) were obtained by normal atmospheric exposure. Microoxic conditions (2% initial O_2_) were established by flushing vials with pure argon, sealing, sterilizing, and then injecting 0.8 mL of pure O_2_ gas per vial. Cultures were incubated at 28 °C with shaking at 200 rpm. Growth was monitored by OD_600_.

For strain SZCCDBA067, samples for measurement of sulfur species (thiosulfate, sulfate, sulfite, tetrathionate) and nitrogen species (nitrate, nitrite) were collected at multiple time points. Supernatants were obtained by centrifugation (10,000 g, 2 min, 4 °C) and stored at −80 °C. Cell-free controls were run in parallel to check for abiotic transformations.

An additional test was performed in strain SZCCDBA067 to compare growth under microoxic conditions using thiosulfate, sulfate, or both as sulfur sources, and thiosulfate concentrations were measured at multiple time points.

### Nitrate assimilation assay

To test whether nitrate could serve as the sole nitrogen source for assimilation, the same four strains were grown aerobically in minimum medium with 20 mM acetate and either 20 mM nitrate or 20 mM ammonium chloride as the sole nitrogen source. Growth was monitored by OD_600_.

### Measurement of sulfur and nitrogen intermediates

Concentrations of thiosulfate, sulfite, and tetrathionate in the medium were measured using the iodometric method (*53*). Sulfate concentrations were determined by barium sulfate turbidimetry (*54*). Nitrite concentrations were measured colorimetrically via the acidic Griess reaction (*55*).Nitrate was reduced to nitrite by vanadium(III) and then measured with the Griess reaction, and its concentration was calculated by subtracting the nitrite concentration from the total NOx (*56*).

### RNA extraction, RNA sequencing, and differential expression analysis

At the late exponential growth phase under both normoxic and microoxic conditions, 10 mL of culture were centrifuged (10,000 g, 5 min, 4 °C). Pellets were resuspended in minimum media and frozen at −80 °C. Total RNA was extracted using the RNAprep Pure Kit DP430 (Tiangen). Ribosomal RNA (rRNA) was depleted with the Ribo-Zero Plus rRNA Depletion Kit (Illumina).

For strain SZCCDBA067, RNA concentrations were normalized and ERCC (External RNA Controls Consortium) spike-in mix (Catalog no. 4456740, Ambion) was added at a 5% ratio relative to the total RNA. Libraries were prepared with the NEBNext Ultra Directional RNA Library Prep Kit and sequenced on a NovaSeq 6000 platform (PE150) at NovoGene. Reads were trimmed with fastp (*57*), mapped to the SZCCDBA067 genome and ERCC sequences using Burrows–Wheeler Aligner (BWA), and read counts per gene were obtained with FADU (*58*). To account for unwanted technical variation, we applied the RUVSeq method (*59*): factors of unwanted variation were estimated using the ERCC spike-in controls, and these factors were included in the design matrix of a DESeq2 model (*60*). Differentially expressed genes were then identified using DESeq2, with a Wald test for significance. Transcripts showing a fold change of at least two and an adjusted P-value ≤0.01 were considered differentially expressed. Absolute transcript abundances (copies per ng total RNA) were calculated from a standard curve generated for each sample based on the known concentrations of ERCC spike-ins and their normalised read counts.

For strain SZCCDBB020, a relative quantification strategy was used (no spike-in). Reads were mapped to the SZCCDBB020 genome, and differential expression was determined with DESeq2. Normalization was based on total number of mapped reads per sample, and the same fold-change and adjusted P-value thresholds were applied.

### RT-qPCR validation

The same RNA samples used for RNA-seq were reverse-transcribed with the iScript cDNA Synthesis Kit (Bio-Rad). Quantitative PCR was performed with the iTaq Universal SYBR Green Supermix (Bio-Rad) and gene-specific primers (Table S7) on a CFX Opus 96 instrument. Relative transcript abundances were normalized to *rpoB*, a commonly used reference gene due to its stable expression under varying conditions (*61, 62*).

### CRISPR-guided cytosine base-editing system

To inactivate specific genes (*rhd* and *dsrA*) in our *Ruegeria* strains, we adapted a cytosine base-editing system previously developed for another marine bacterium (*19*). Unlike traditional CRISPR-Cas9, which creates double-strand DNA breaks, base editing converts a single cytosine (C) to thymine (T) without cutting the DNA backbone (*18*). This is achieved by three components delivered on a single plasmid (pBE): a nuclease-deactivated Cas9 (dCas9) from *Streptococcus pyogenes* that binds but does not cut DNA; a cytidine deaminase (PmCDA1) from sea lamprey (*Petromyzon marinus*) that changes C to uracil (U) on exposed single-stranded DNA; and a uracil DNA glycosylase inhibitor (UGI) that prevents removal of the U, making the change permanent.

The dCas9-PmCDA1-UGI fusion is expressed under an IPTG (isopropyl β-D-1-thiogalactopyranoside)-inducible *lacI*-P_trc_ promoter. A separate, constitutively expressed single-guide RNA (gRNA) contains a 20-nucleotide spacer complementary to the target genomic sequence (Fig. S7). For binding to occur, the target DNA must contain a short protospacer-adjacent motif (PAM) immediately downstream of the spacer-complementary region. We used the dCas9 from *S. pyogenes*, which recognizes the PAM sequence NGG (where N is any nucleotide). Upon binding, the dCas9-gRNA complex locally unwinds the DNA, creating a small single-stranded bubble. The PmCDA1 deaminase then acts on a cytidine located approximately 16 to 20 nucleotides upstream of the PAM (the active editing window). After DNA replication or repair, the original C-G pair is permanently converted to a T-A pair. By designing the gRNA spacer to target a CAG (glutamine) codon, we convert it to a TAG stop codon, thereby generating a premature nonsense mutation that inactivates the gene.

For each target locus, a 20-bp spacer was designed such that the target cytidine lay within the deamination window (positions −20 to −16 relative to the PAM-proximal nucleotide, designated −1). The spacer was introduced into a pTemplate plasmid by inverse PCR, and the complete gRNA cassette was then assembled into pBE using the In-Fusion Snap Assembly Master Mix (Takara Bio). All strains, plasmids, and primers are listed in Tables S8-S10.

Electrocompetent cells of *R. pomeroyi* DSS-3 and SZCCDBA067 were prepared by growing strains in MB2216 liquid medium at 30 °C with shaking (180 rpm) to OD_600_ ≈ 0.3, harvesting by centrifugation, washing three times with ice-cold 10% (v/v) glycerol, and resuspending in the same solution. Aliquots (80 µL) were stored at −80 °C. For electroporation, 500 ng of plasmid DNA were mixed with 80 µL of competent cells, transferred to a 0.1-cm cuvette (Bio-Rad), and pulsed at 1.8 kV using a MicroPulser electroporator (Bio-Rad). Immediately after pulsing, 1 mL of fresh MB2216 was added for recovery. Cells were incubated at 30 °C for 6 h, then transferred to MB2216 containing gentamicin (20 µg mL^-1^) and grown overnight.

Base editing was induced by adding IPTG to a final concentration of 0.1 mM for 24 h. To maximize editing efficiency, a second induction passage was performed: 200 µL of the induced culture were transferred to fresh MB2216 with IPTG and gentamicin and incubated for another 24 h. Cultures were then plated onto MB2216 agar with gentamicin. Individual colonies were screened by colony PCR, and the target locus was amplified and sequenced (Sanger) to confirm the desired C-to-T mutation. Detailed workflow can be found in Fig. S6.

After confirmation, the editing plasmid was cured by passaging the edited strains twice in antibiotic-free MB2216 liquid medium. Cells were then streaked onto non-selective MB2216 agar. Single colonies were tested for plasmid loss by colony PCR (absence of plasmid-specific bands) and by loss of gentamicin resistance (no growth on MB2216 with 20 µg mL^-1^ gentamicin). The final plasmid-free, edited mutant strains were stored at −80 °C in 15% glycerol. Detailed workflow can be found in Fig. S8.

Using this system, we introduced early stop codons into *dsrA* and the CASGXR-containing rhodanese gene (*rhd*, locus tag SZCCDBA067_01274) in SZCCDBA067 with gRNA spacers listed in Table S11. For each target, the editing efficiency after two induction passages was 100% as verified by Sanger sequencing. The resulting mutants (Δ*dsrA* and Δ*rhd*) were cured of the editing plasmid before phenotypic characterisation.

### Thiosulfate consumption assay for mutant strains

Wild-type SZCCDBA067, the Δ*dsrA* mutant, Δ*rhd* mutant, and a no-cell control were incubated under microoxic conditions (2% initial O_2_) in minimum medium containing 10 mM thiosulfate, 20 mM acetate and 20 mM nitrate. Thiosulfate concentration, sulfite concentration and cell density (OD_600_) were monitored over time as described above. All assays were performed in triplicate.

### Metagenomic analysis of rDsr abundance

To estimate the relative abundance and distribution of rDsr genes in marine environments, we constructed reference phylogenetic trees for DsrA and DsrB. First, we updated a previously reported DsrAB database containing both reductive and oxidative types of bacterial or archaeal origin (*63*). Oxidative bacterial-type DsrAB sequences from Muller et al. (*63*) were used as seeds to search against NCBI’s non-redundant databases, and novel candidates were added based on a bit-score threshold. The resulting DsrA and DsrB sequences were merged with those from the newly discovered *Ruegeria* lineage and the existing database. Sequences were aligned using MAFFT v7.125 (*48*) and trimmed with TrimAl v1.4.rev15 (*49*). Maximum-likelihood phylogenetic trees for DsrA and DsrB were constructed separately using raxmlHPC v8.2.12 with the setting “-m PROTGAMMALGX” (*64*), visualized using iTol (*65*), and used as the reference trees for downstream phylogenetic placement analysis.

We collected the *Tara* Oceans metagenomic sequencing data (Table S5) and 305 coastal sediment metagenomes (Table S6) from global sites (salt marshes, coral reefs, mangroves, tidal flats). Raw reads were quality-trimmed with Trimmomatic v0.36 and mapped to reference sequences of DsrA, DsrB, SoxA and SoxB using the blastx function in DIAMOND v2.0.4 (*66*) with the setting ‘-e 1e-5 --id 30 --query-cover 50’. The mapped reads were then phylogenetically placed onto the respective reference tree using pplacer v1.1 (*67*). For DsrA and DsrB, the relative contribution of the Rhodobacterales order was calculated as the number of reads assigned to the Rhodobacterales (threshold of >=10 reads) divided by the total number of reads belonging to the oxidative bacterial type in each reference tree. For SoxA and SoxB, which lack distinct oxidative and reductive types, the relative contribution was calculated similarly but using all mapped reads without subtype classification.

To determine the copy number of *dsrA* and *dsrB* per genome, we extended the analysis to 21 phylogenetic marker genes that typically occur once per genome (*68–70*). These marker genes included *polA* (DNA polymerase I, K02335), *rpoC* (beta subunit of DNA-directed RNA polymerase, K03046), the beta subunit of the proton-translocating ATPase (K02112), *tpi* (triosephosphate isomerase, K01803), *secY* (preprotein translocase subunit, K03076), *ruvB* (holliday junction DNA helicase, K03551), *radA* (DNA repair protein, K04485), and 14 genes encoding ribosomal proteins (K02863, K02871, K02878, K02886, K02906, K02931, K02933, K02948, K02952, K02965, K02967, K02988, K02992, K02994). The copy number of each target gene was calculated based on the number of reads assigned to the oxidative bacterial-type DsrA or DsrB in the reference tree, normalized by the average number of reads mapped onto 21 single-copy marker genes and by their respective gene lengths.

For *Tara* Oceans samples, we categorized sampling sites into four oxygen regimes according to Berg et al (*71*): oxic (O_2_ > 60 µM), microoxic (5 µM < O_2_ < 60 µM), suboxic (1 µM < O_2_ < 5 µM) and nanooxic (O_2_ < 1 µM). Enrichment patterns of *dsrA*, *dsrB*, *soxA* and *soxB* were assessed using the Wilcoxon rank-sum test as implemented in the ggpubr package. For coastal sediment metagenomes, copy numbers of Rhodobacterales-associated and community-level *dsrA* and *dsrB* were calculated similarly, and their global distribution was mapped.

### Statistical analysis

All growth and chemical measurements were performed with three biological replicates. Data are presented as means ±standard deviation. Comparisons between conditions were made using Student’s t-test or Wilcoxon test as indicated, with *P* <0.05 considered significant. For RNA-seq, differential expression was assessed with DESeq2 using an adjusted P-value threshold of 0.01 and fold change ≥ 2.

## Acknowledgements

This work is supported by the National Natural Science Foundation of China (42506111, 22578250), the Hong Kong Research Grants Council (RGC) General Research Fund (GRF) (14107625), and the Natural Science Foundation of Fujian Province (2025J01612).

## Data availability

The genomic sequences of the 10 newly sequenced *Ruegeria* strains, along with the raw reads of transcriptome for SZCCDBA067 and SZCCDBB020, have been uploaded to NCBI (Project ID: PRJNA1124084) with the reviewer link https://dataview.ncbi.nlm.nih.gov/object/PRJNA1124084?reviewer=mcdhhka8ij2bkd7kd971 muph5i.

