## Supplementary Figures and Tables for "A standalone reverse dissimilatory sulfite reductase (rDsr) pathway enables sulfur oxidation under oxygen limitation"

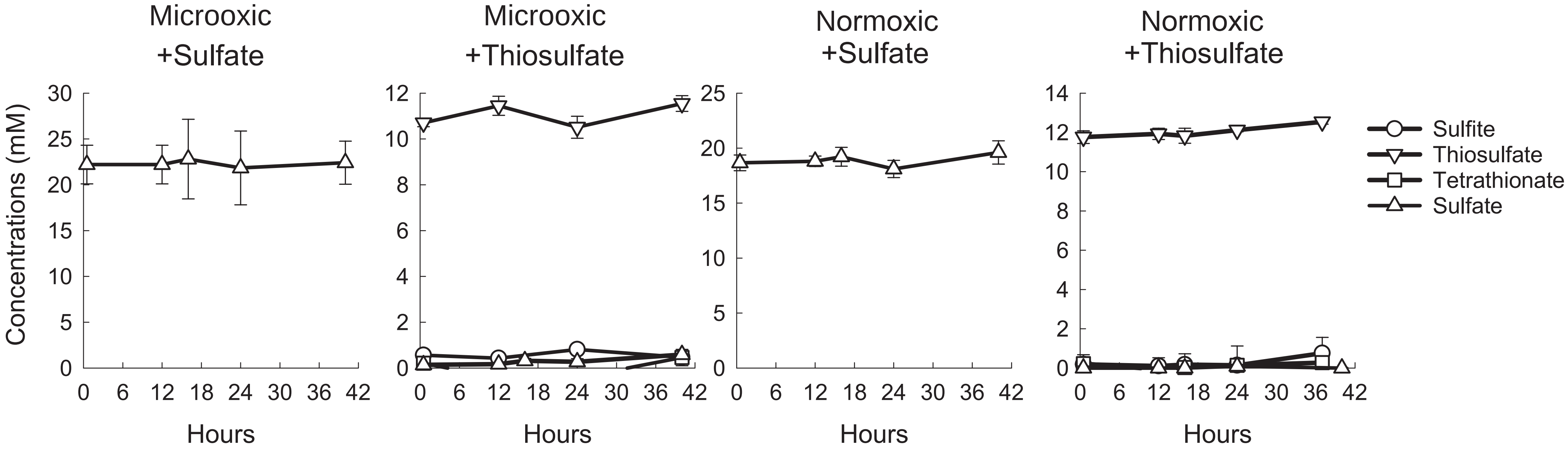


**Supplementary Figure 1. Concentrations of sulfur intermediates in blank samples without bacterial cells.** Error bars indicate the standard deviation from three biological replicates.

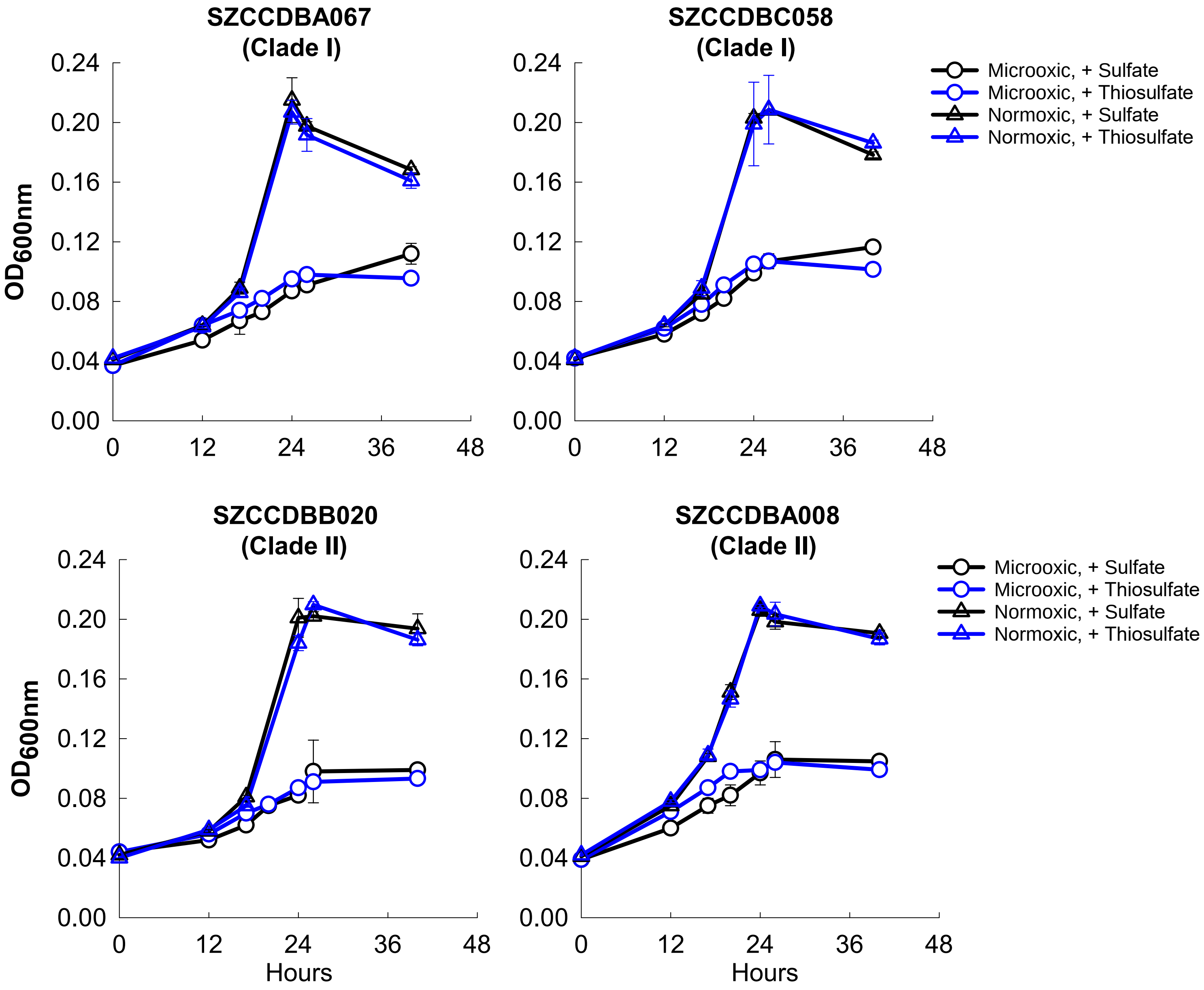


**Supplementary Figure 2. Growth of two representative strains from each clade under normoxic (21% oxygen) or microoxic (2% initial headspace oxygen) conditions, with thiosulfate or sulfate as the sole sulfur source.** Acetate (20 mM) was supplied as the sole carbon source. Error bars indicate the standard deviation from three biological replicates.

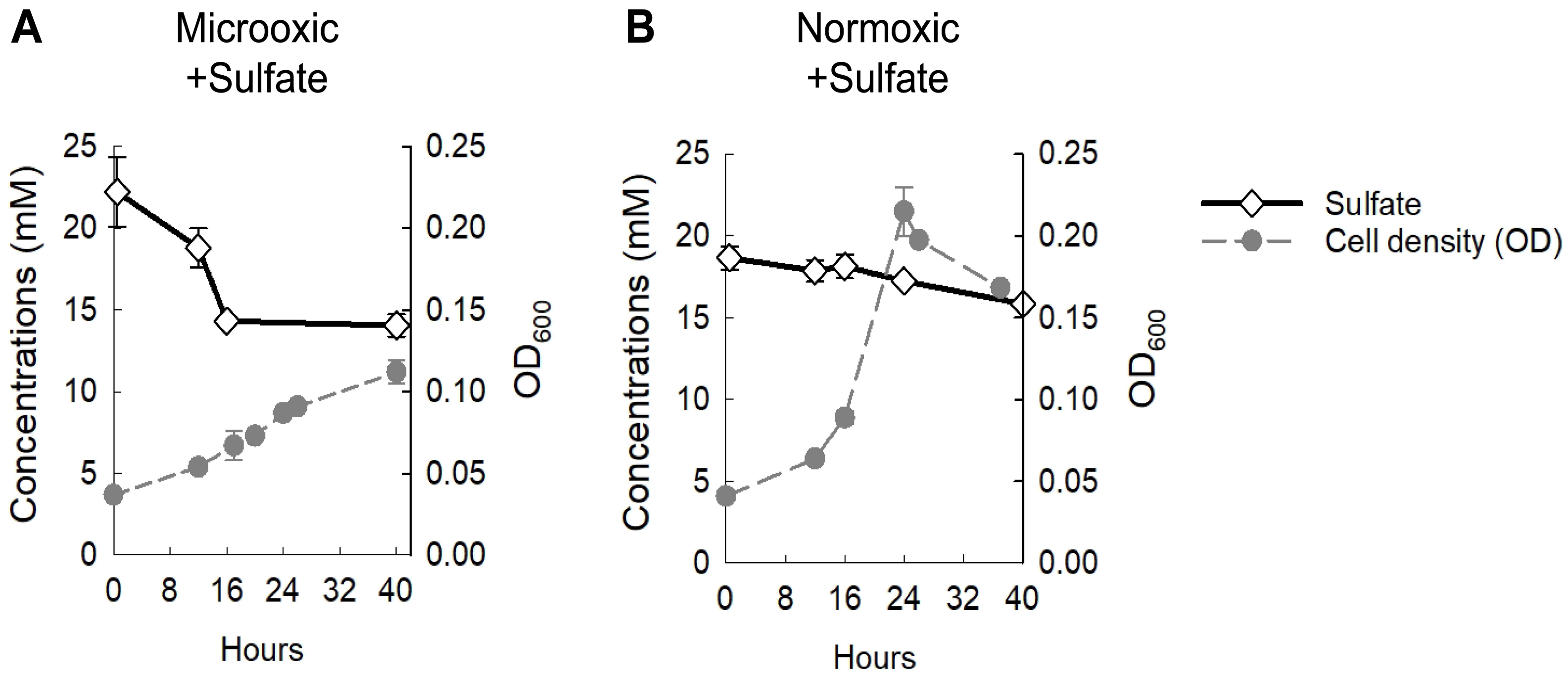


**Supplementary Figure 3. Growth of strain SZCCDBA067 with sulfate as the sole sulfur source under microoxic (A) and normoxic (B) conditions.** Sulfate concentration is shown on the left y-axis, and cell de nsity (estimated by measuring optical density at 600 nm, $OD_{600}$) is shown on the right y-axis. Error bars represent the standard deviation from three biological replicates.

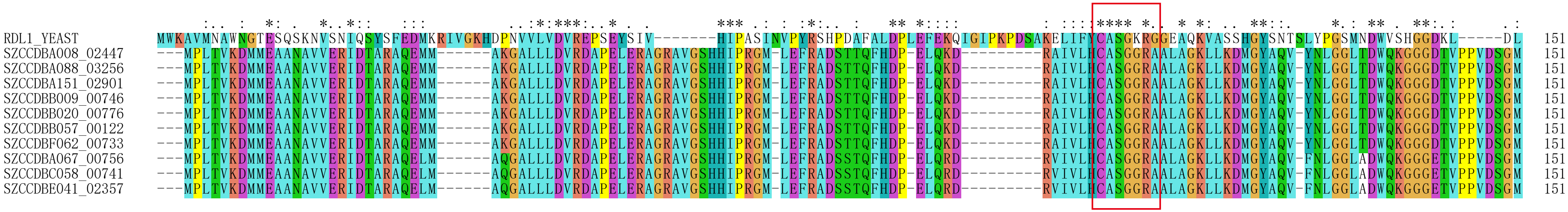


**Supplementary Figure 4. Sequence alignment of rhodaneses encoded by *Saccharomyces cerevisiae* (*RDL1*) and strains of the intertidal *Ruegeria* lineage.** The red box indicates the active-site loop of the catalytic domain in *RDL1* (CASGXR). X refers to the occurrence of varying amino acids in the multiple sequence alignment.

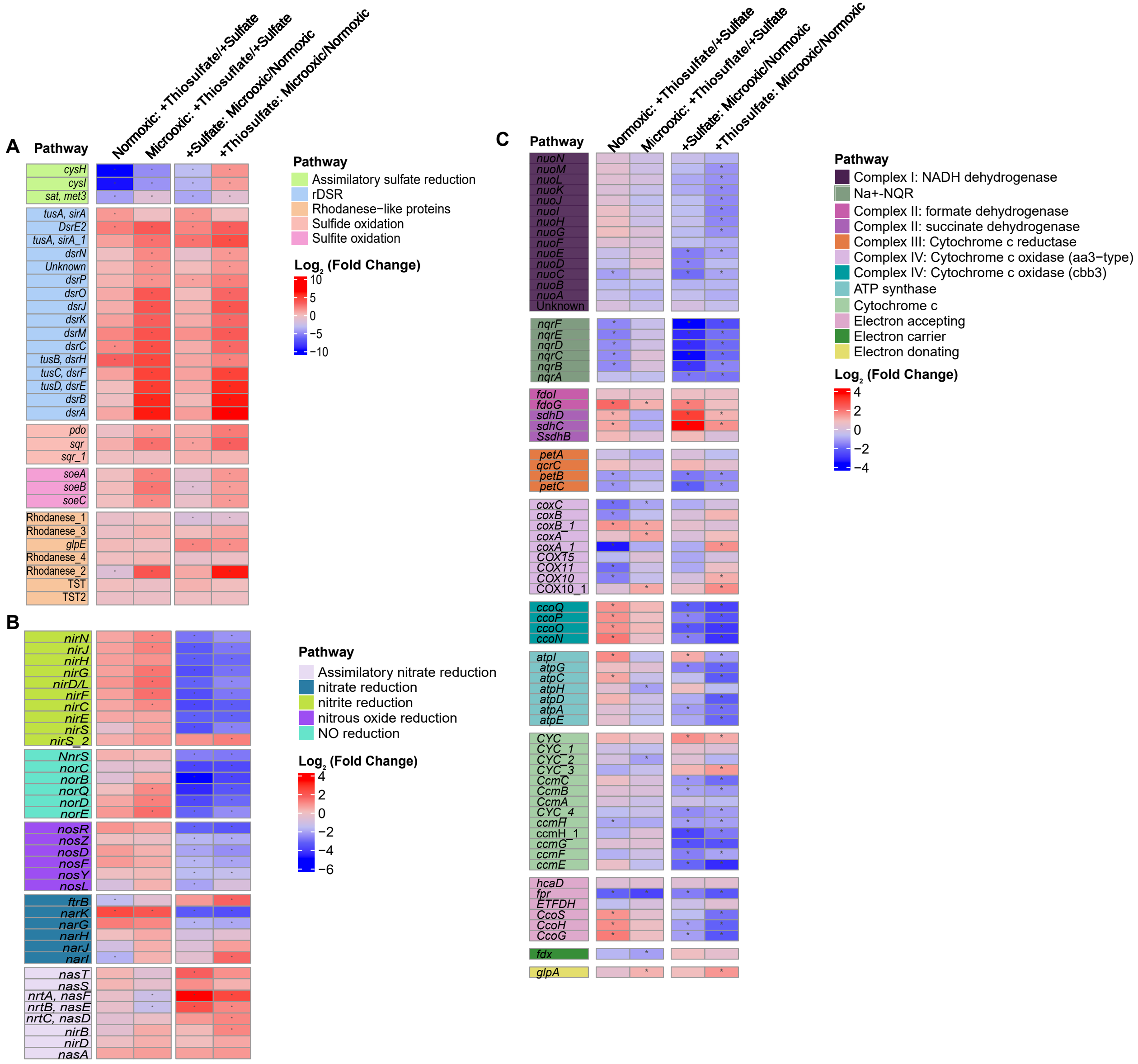


**Supplementary Figure 5. Relative transcript abundances of genes involved in inorganic sulfur oxidation (A), denitrification and assimilatory nitrate reduction (B), and electron transport chain (C).** Transcript abundances were investigated in the strain SZCCDBB020 under four conditions: i) normoxic with thiosulfate; ii) microoxic with thiosulfate; iii) normoxic with sulfate; iv) Microoxic with sulfate. Therefore, four comparisons in the transcript abundances were made and indicated above the panels. The $\log_2$ fold changes of relative transcript abundances are shown in color bars. Stars indicate $\log_2$ fold change >1 and adjusted $p$ values < 0.01.

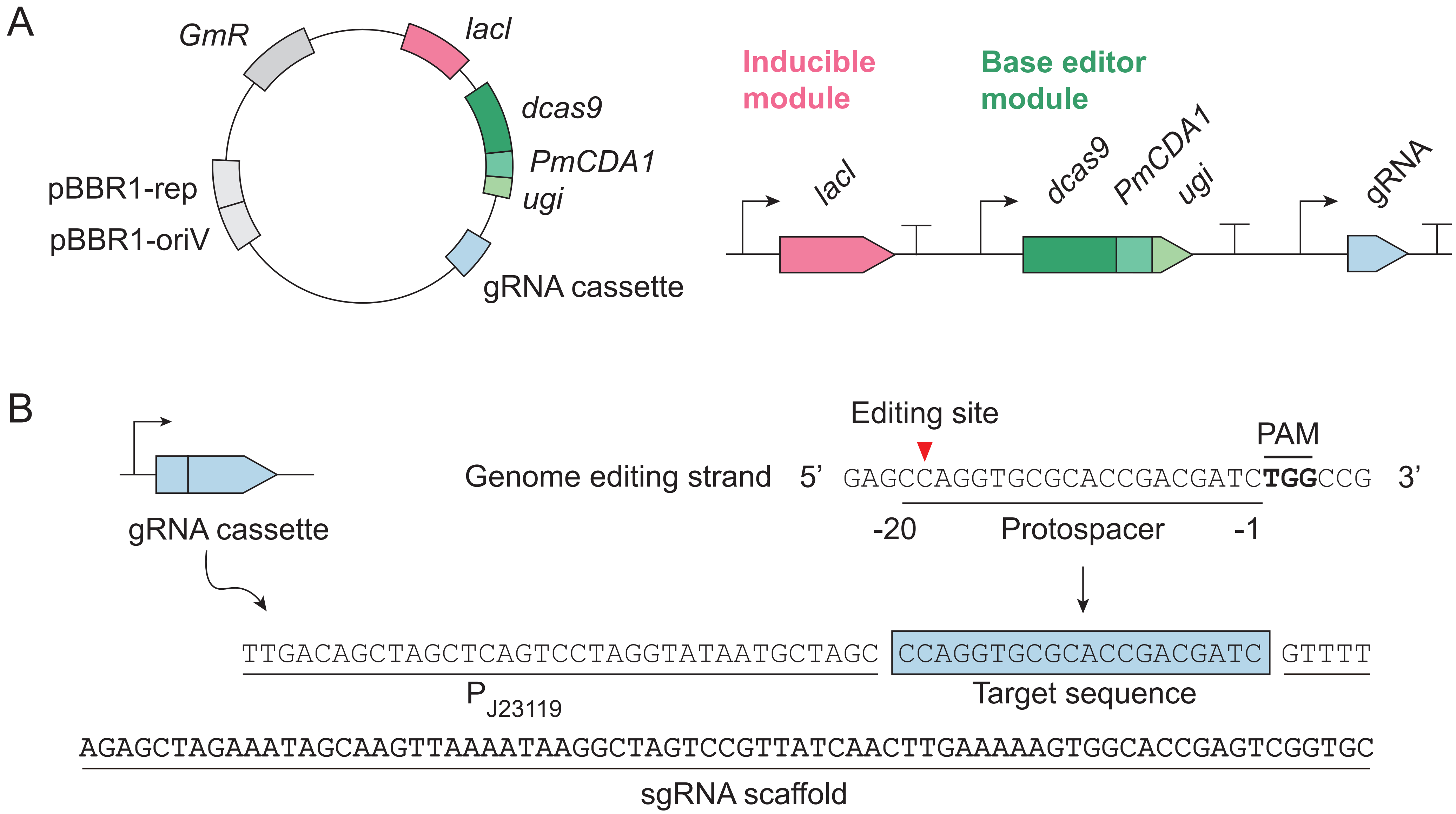


**Supplementary Figure 6. Design of the base editing plasmid and gRNA expression cassette.** (A) Design of the base editing plasmid pBE, which contains a *lacI*-$P_{trc}$ inducible module driving expression of *dCas9*, *PmCDA1*, and *ugi*, together with a constitutively expressed gRNA cassette. (B) Architecture of the gRNA cassette and target-site layout. The gRNA cassette is driven by $P_{J23119}$ and contains a 20-nt spacer (blue) fused to the sgRNA scaffold. Protospacer positions are numbered relative to the PAM-proximal nucleotide (1), and the edited cytidine is indicated in red.

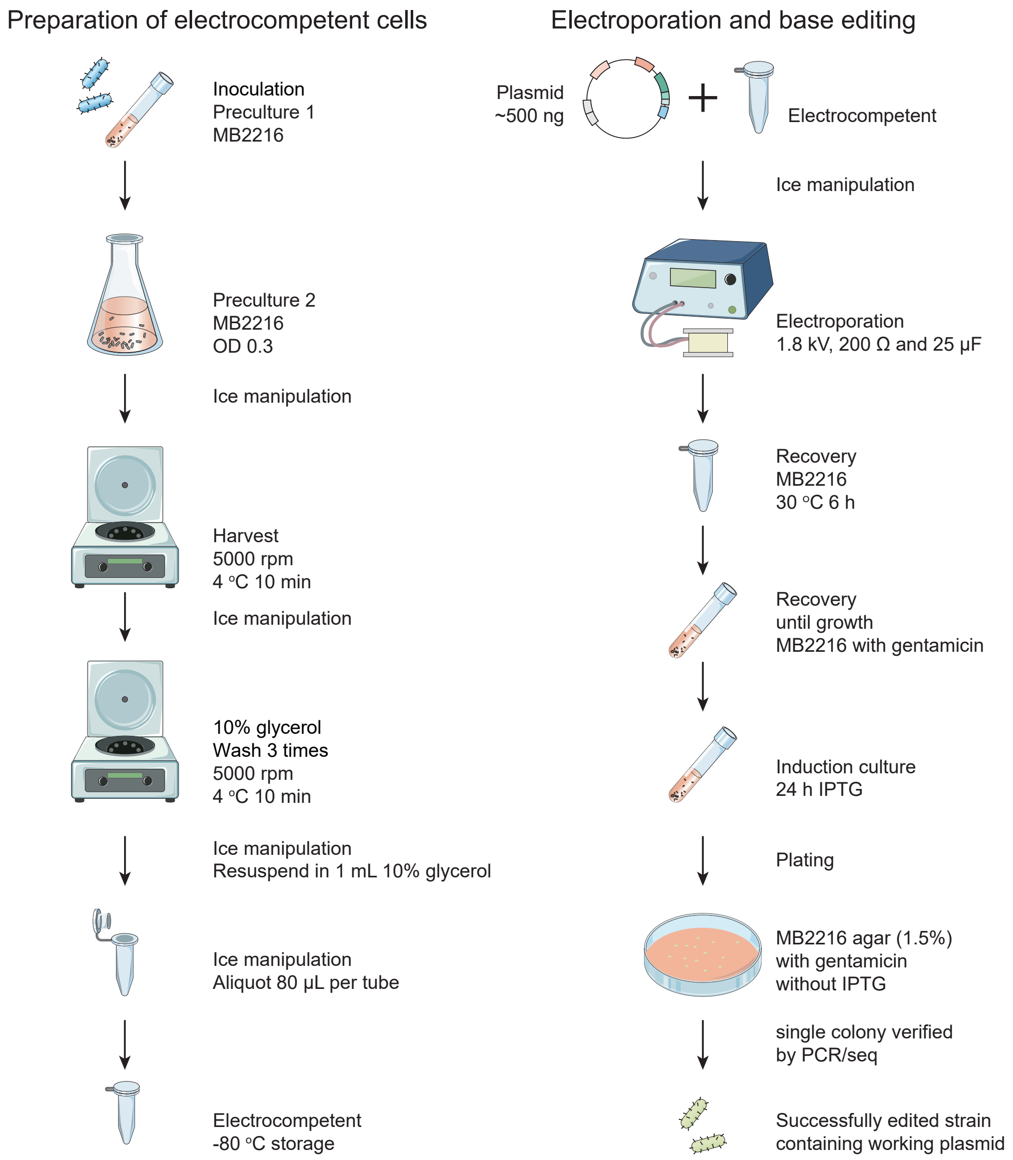


**Supplementary Figure 7. Schematic workflow of the base editing process.** The procedure is divided into two main phases. The left panel outlines the preparation of electrocompetent cells, including cell cultivation to early exponential phase, cold harvesting, and repeated washing with 10% glycerol. The right panel illustrates plasmid electroporation, followed by cell recovery, IPTG-mediated induction of the base editor, and final screening of mutant colonies by sequencing.

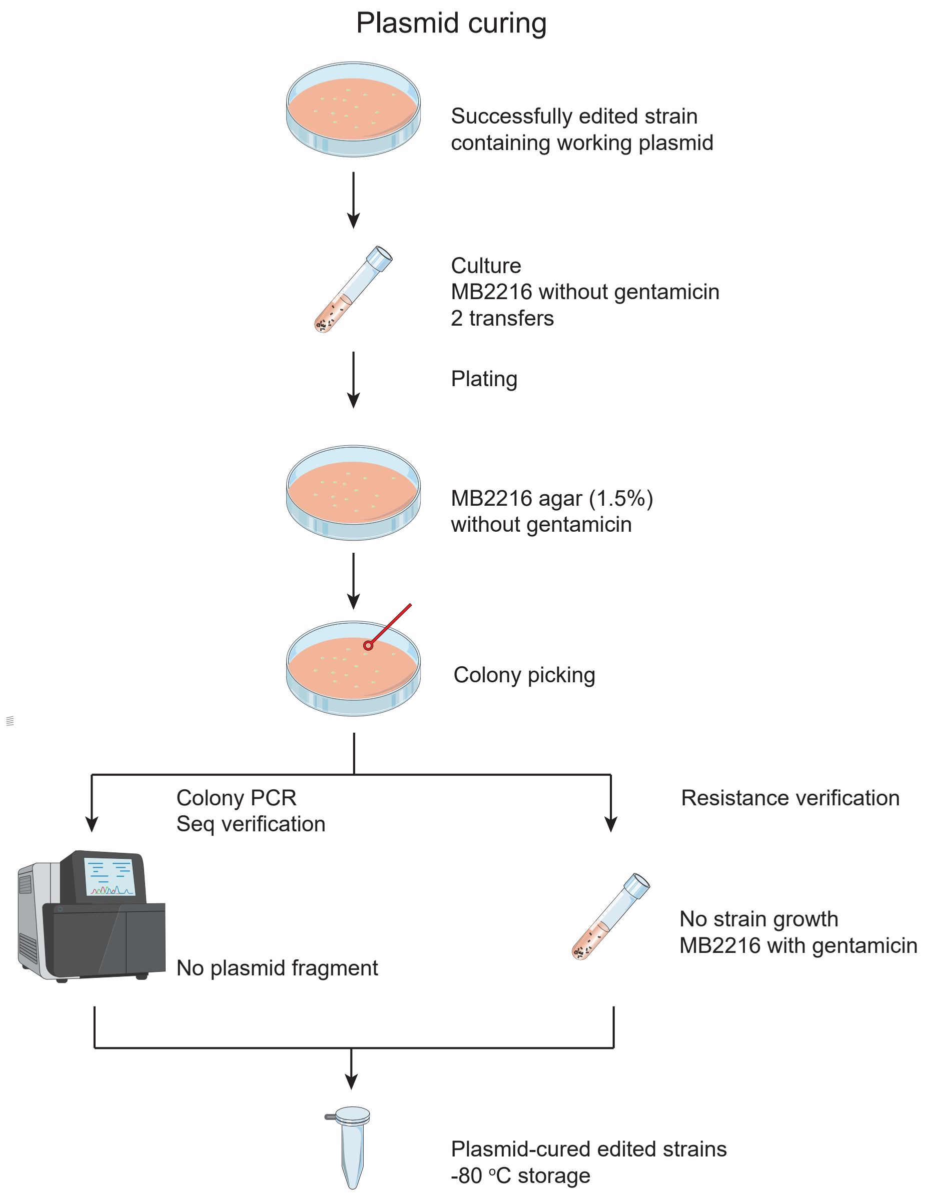


**Supplementary Figure 8. Schematic workflow of the plasmid curing process.** Successfully edited strains harboring the working plasmid are cultured and passaged for two successive generations in antibiotic-free MB2216 medium. After isolation on antibiotic-free agar plates, single colonies are subjected to a dual-verification process. Successful plasmid clearance is confirmed by both the absence of specific fragments of plasmid via colony PCR and the loss of gentamicin resistance (no cell growth in gentamicin-supplemented medium). The validated, plasmid-cured strains are subsequently cryopreserved at -80 °C.

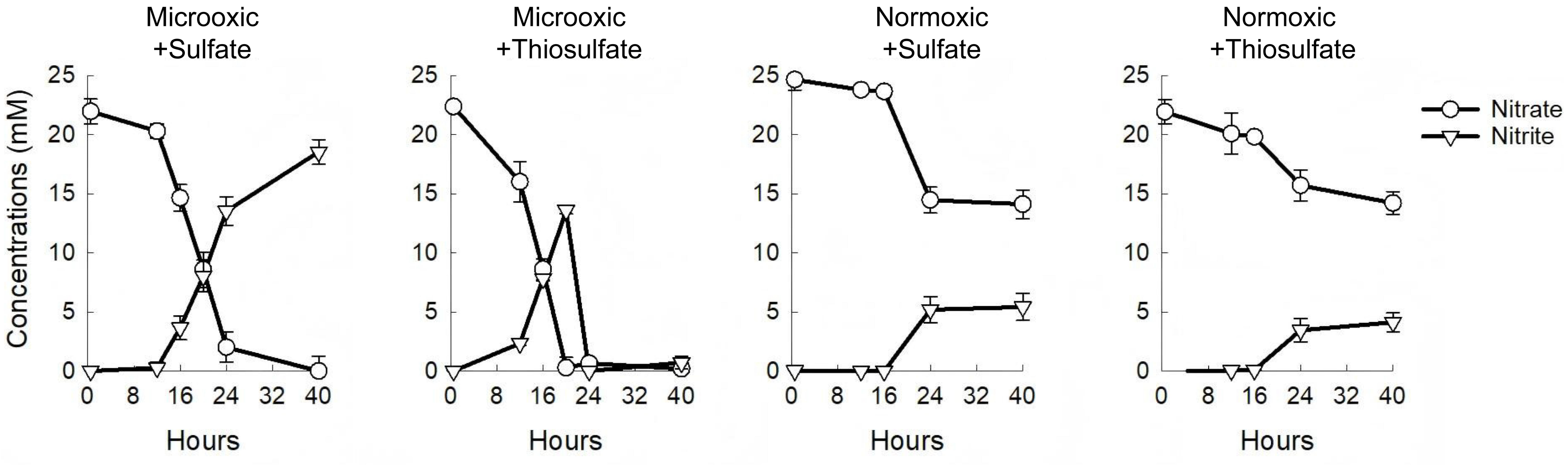


**Supplementary Figure 9. Profiles of nitrate and nitrite during normoxic and microoxic growth of strain SZCCDBA067 using thiosulfate (10 mM) or sulfate (20 mM) as the sole sulfur source.** Error bars represent the standard deviation from three biological replicates.

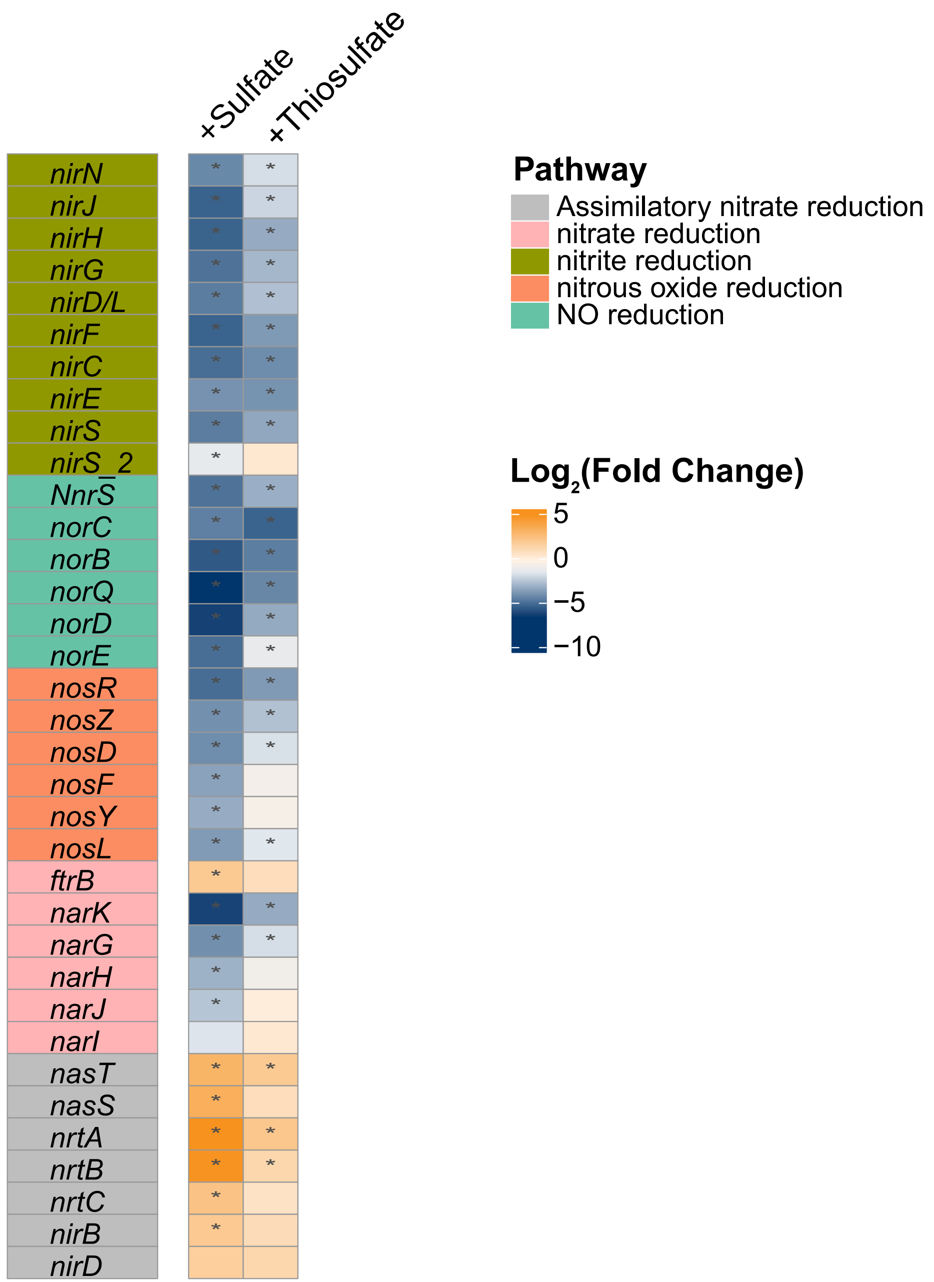


**Supplementary Figure 10. Relative transcript abundances of genes involved in denitrification and assimilatory nitrate reduction.** Transcript abundances of the strain SZCCDBA067 under microoxic conditions were compared to those under normoxic conditions, and $\log_2$ fold changes were shown in color bars. Stars indicate $\log_2$ fold change >1 and adjusted *P* values < 0.01. See Table S3 for more information related to the gene abbreviations.

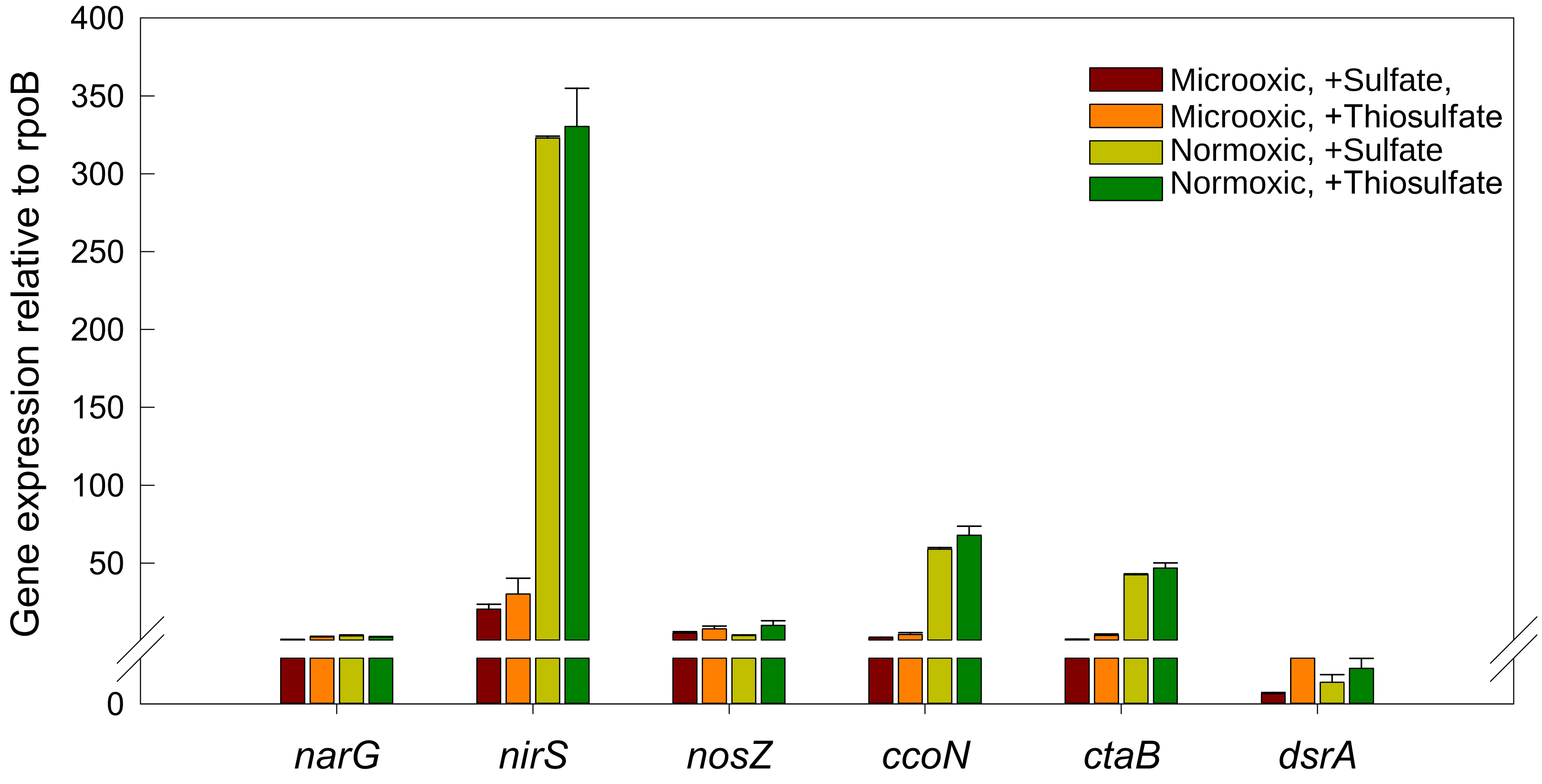


**Supplementary Figure 11. Expression of selected genes involved in denitrification, aerobic respiration and sulfur oxidation by reverse-transcription quantitative PCR (RT-qPCR).** RNA samples for RNA-seq were aliquoted for RT-qPCR. The expression of rpoB was used as the internal control. NarG, Respiratory nitrate reductase alpha chain; NirS, Nitrite reductase; NosZ, Nitrous oxide reductase; CcoN, Cytochrome c oxidase (*cbb3*-type) subunit CcoN; CtaB, Heme O synthase; DsrA, Dissimilatory sulfite reductase, alpha subunit.

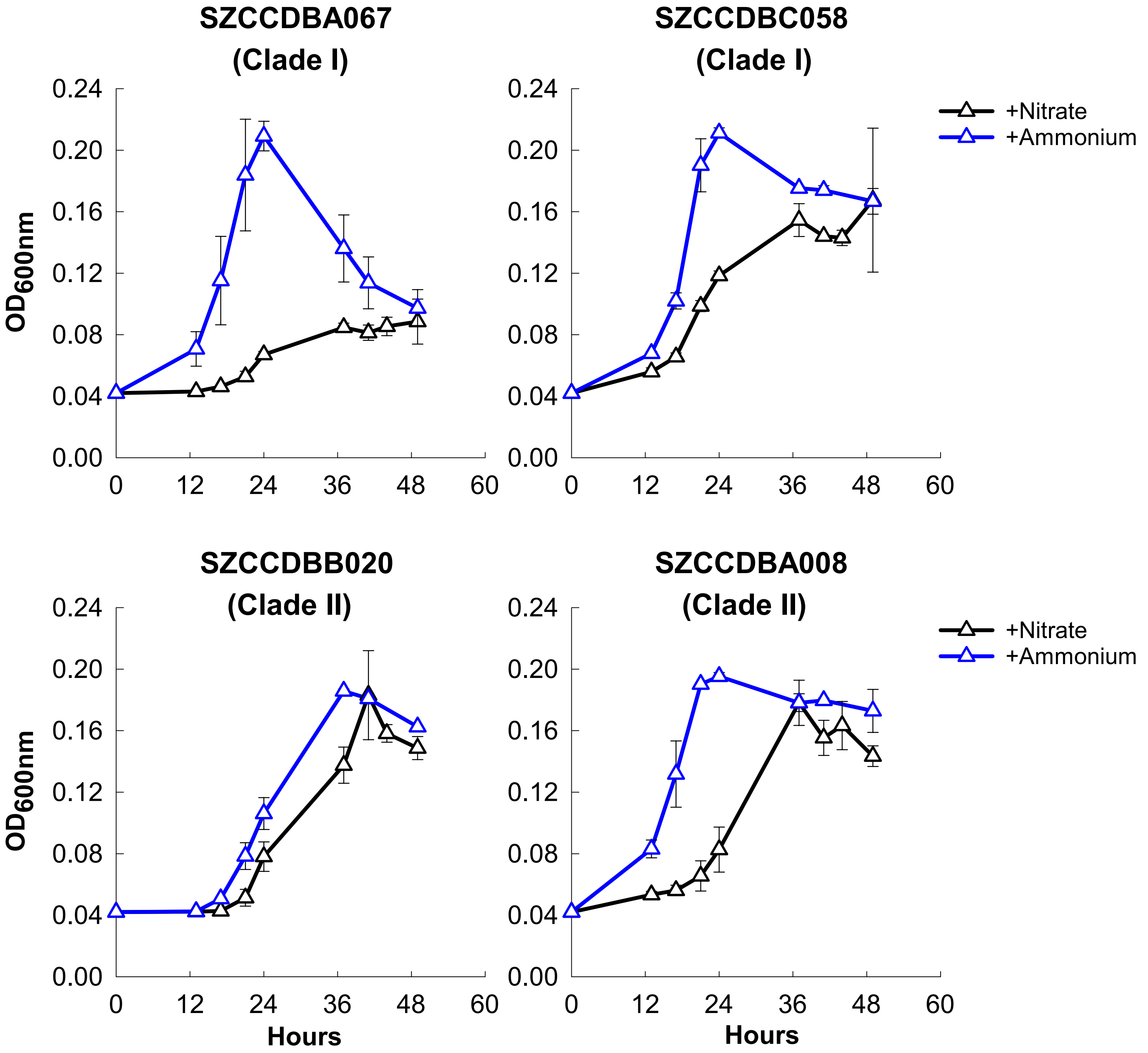


**Supplementary Figure 12. Growth of representative strains under normoxic conditions with either nitrate or ammonium as the sole nitrogen source.** Error bars indicate the standard deviation from three biological replicates.

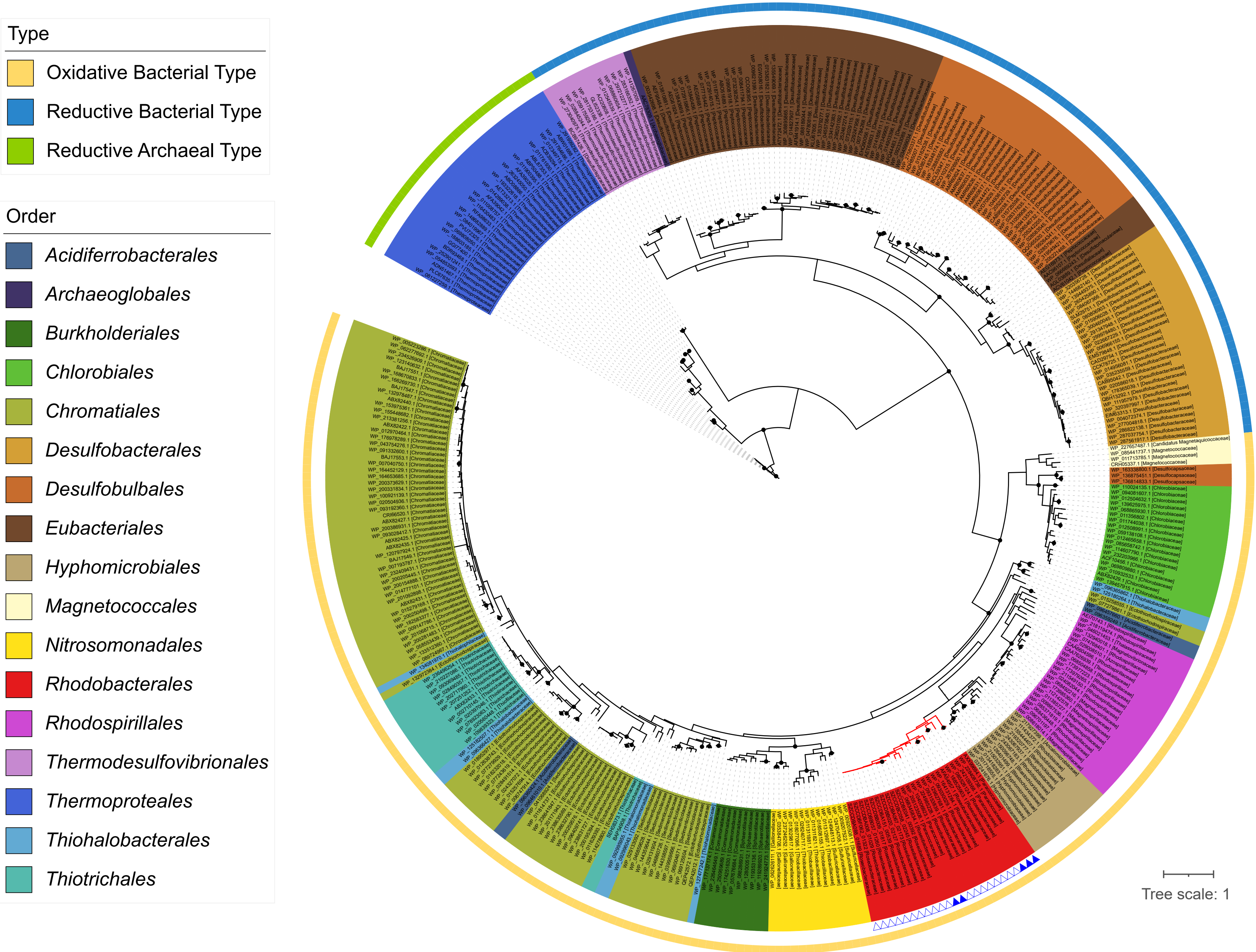


**Supplementary Figure 13. DsrA reference tree used for phylogenetic placement analysis.** The maximum likelihood phylogeny was constructed using raxmlHPC. Nodes with bootstrap support of 100% are denoted by solid circles. Taxonomic family is indicated in brackets next to the Accession ID on the leaf label. Color ranges represent taxonomic order. The type of each DsrA sequence (reductive archaeal-, reductive bacterial-, or oxidative bacterial-type) is shown in the outer ring. Branches of the Rhodobacterales order are highlighted in red, with families Roseobacteraceae and Paracoccaceae distinguished by open and filled blue triangles, respectively.

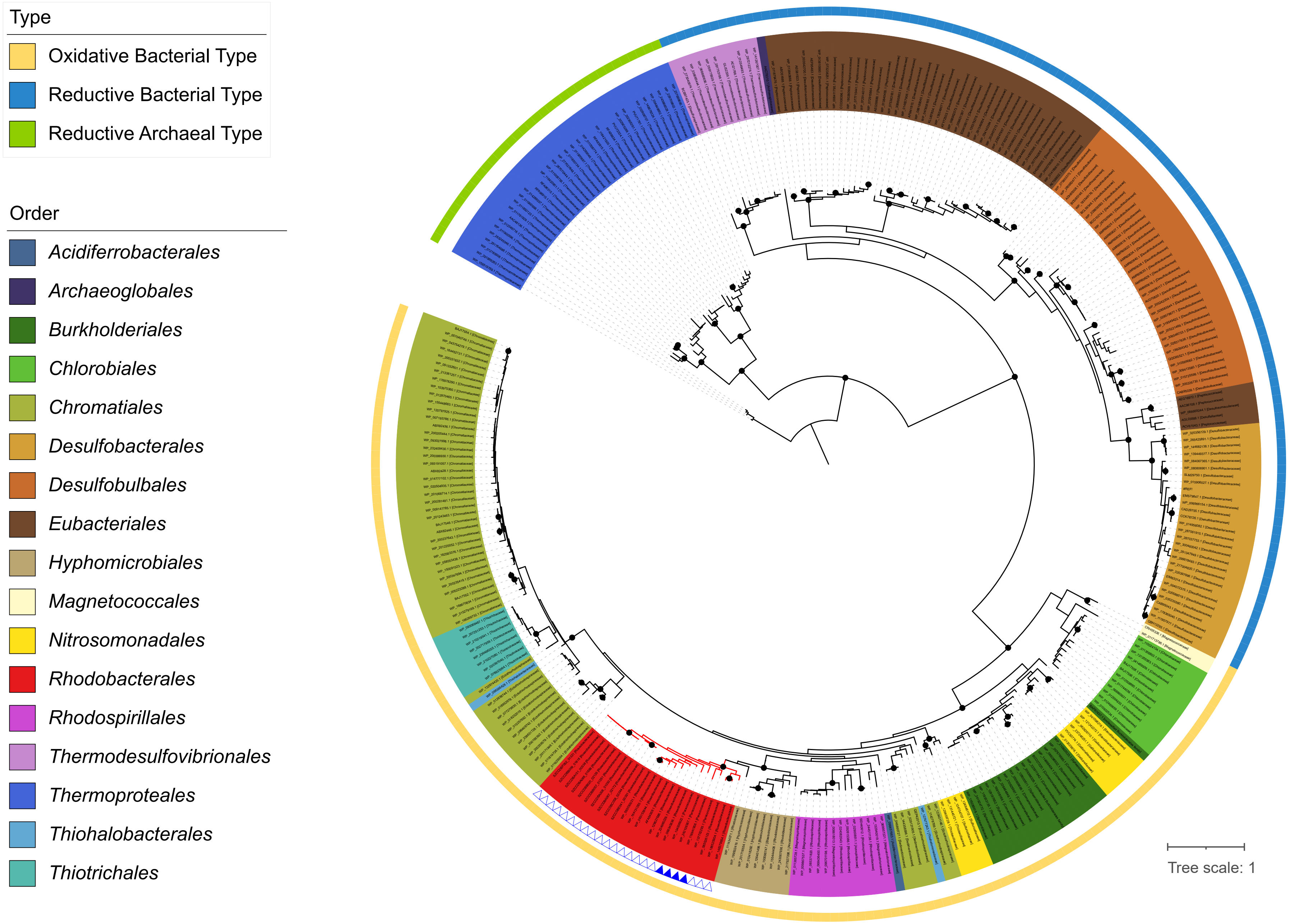


**Supplementary Figure 14. DsrB reference tree used for phylogenetic placement analysis.** The maximum likelihood phylogeny was constructed using raxmlHPC. Nodes with bootstrap support of 100% are denoted by solid circles. Taxonomic family is indicated in brackets next to the Accession ID on the leaf label. Color ranges represent taxonomic order. The type of each DsrB sequence (reductive archaeal-, reductive bacterial-, or oxidative bacterial-type) is shown in the outer ring. Branches of the Rhodobacterales order are highlighted in red, with families Roseobacteraceae and Paracoccaceae distinguished by open and filled blue triangles, respectively.

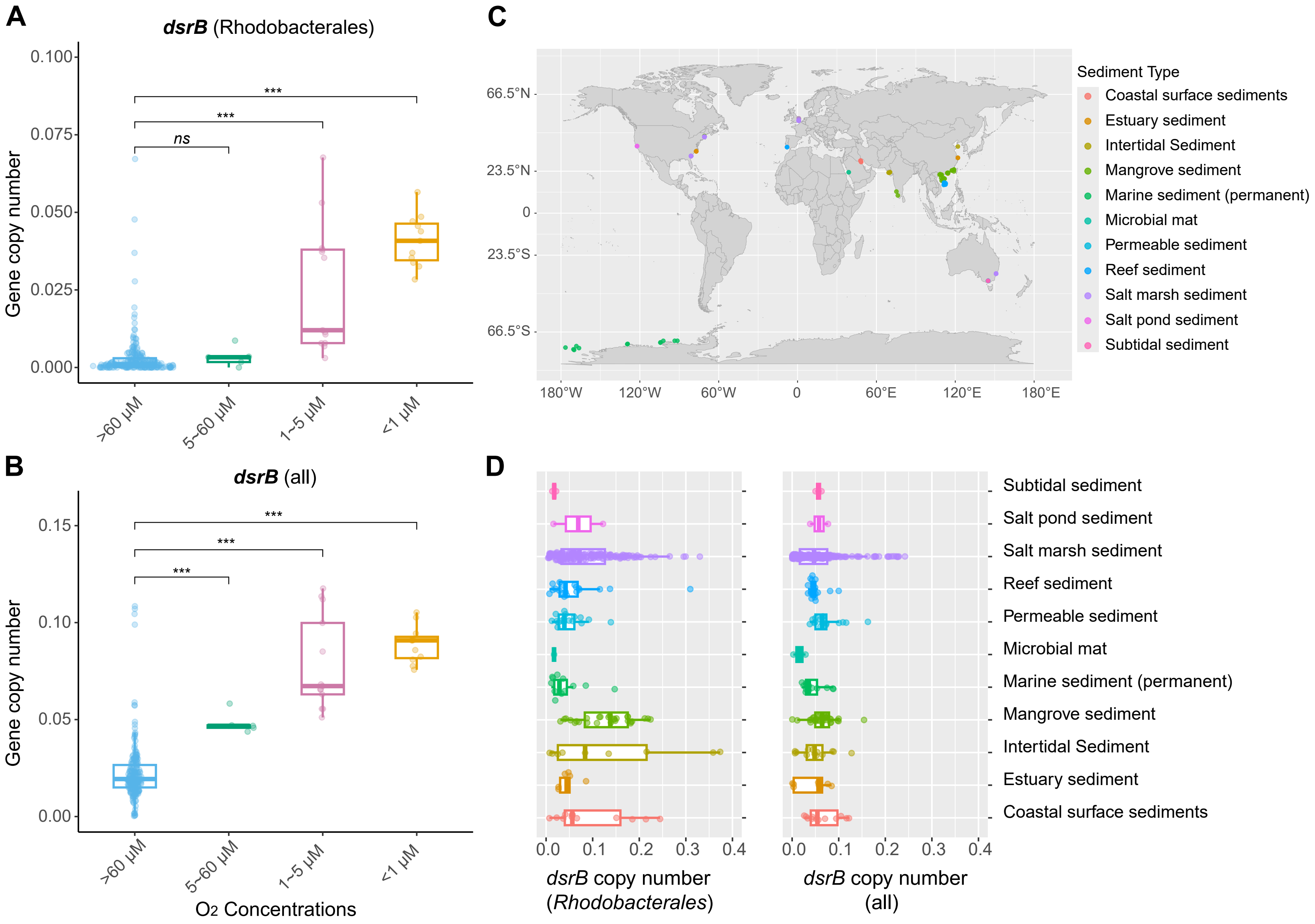


**Supplementary Figure 15. Copy numbers of *dsrB* in marine metagenomes.** (**A and B**) Copy number (per cell) of *dsrB* among *Tara* Oceans metagenomes sampled from sites with different oxygen levels. The copy number was estimated based on the number of reads assigned to the oxidative bacterial-type DsrB in the reference tree, normalized by the average number of reads mapped onto 21 single-copy marker genes and by their respective gene length. **(A)** Copy numbers of Rhodobacterales-associated *dsrB* and **(B)** of *dsrB* at the bacterial community level are represented by dots. The median is represented by the center line of the box, while the upper and lower quartiles are indicated by the box limits. The whiskers show 1.5 times the interquartile range. The *Tara* Oceans samples were categorized into four groups based on the oxygen concentration at the sampling site: oxic (> 60 µM), microoxic (> 5 µM but < 60 µM), suboxic (> 1 µM but < 5 µM) and nanooxic (< 1 µM). The stars denote a *p*-value of less than 0.001 (Wilcox test). (**C and D**) Copy number of *dsrB* genes in coastal sediment metagenomic samples and their geographic distribution. **(C)** Geographic distribution of different sediment types. **(D)** Copy numbers of Rhodobacterales-associated *dsrB* and of *dsrB* at the community level as indicated by the solid dots. Dots are color-coded based on sediment type.

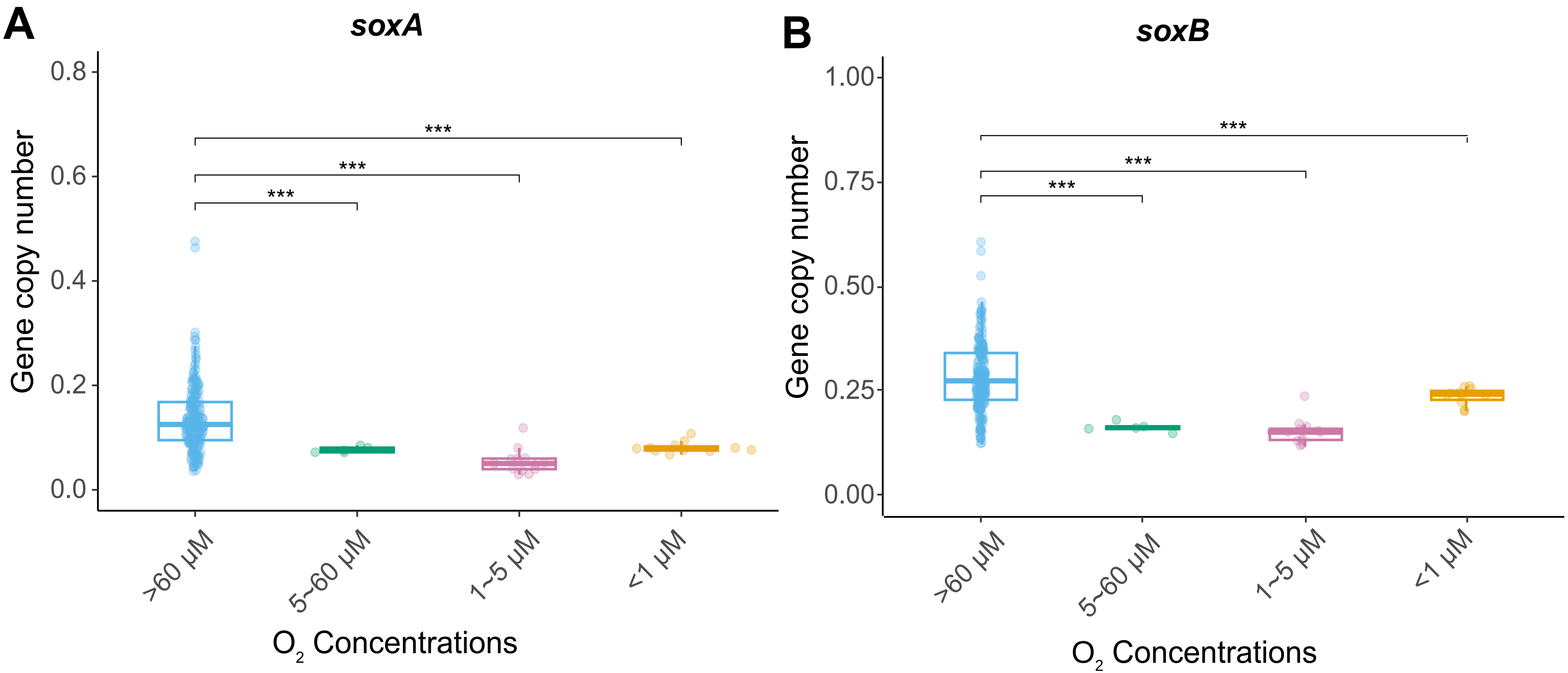


**Supplementary Figure 16. Copy numbers of (A) *soxA* and (B) *soxB* among *Tara* Oceans metagenomes from sites with different oxygen levels** .Copy numbers of genes at the bacterial community level are represented by dots. The median is represented by the center line of the box.The upper and lower quartiles are denoted by the box limits, while whiskers show 1.5 times the interquartile range. Stars indicate statistical significance ($P < 0.05$, Wilcox test).

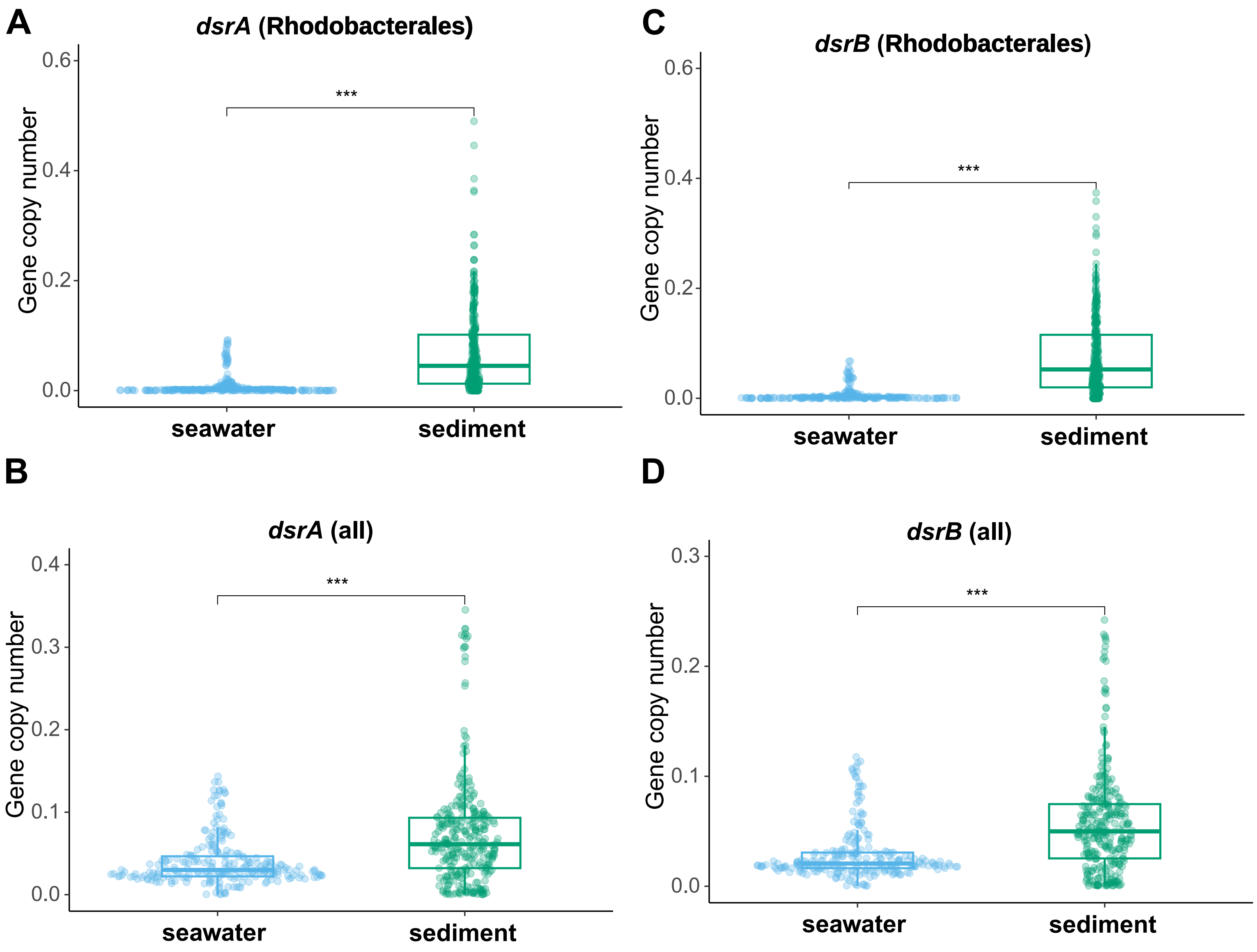


**Supplementary Figure 17. (A, B) Comparison between *dsrA* and (C, D) *dsrB* copy numbers in seawater and coastal sediments.(A, C)** Copy numbers of genes assigned to the Rhodobacterales order and **(B, D)** of genes at the community level are represented by dots. The median is represented by the center line of the box. The upper and lower quartiles are denoted by the box limits, while whiskers represent 1.5 times the interquartile range. Stars indicate statistical significance ($P < 0.001$, Wilcox test).

**Table S1. Summary of genome assembly statistics of the strains isolated from the surface sediment of tidal flats.**

| Strain | GC (%) | Completeness | Contamination | Number of contigs | Closed genome | Number of plasmids[a] | Total size (bp) | N50 (bp) | Number of CDS |
|---|---|---|---|---|---|---|---|---|---|
| SZCCDBA008 | 62.3 | 99.57 | 1.25 | 1 | Yes | 0 | 3,876,361 | 3,876,361 | 3,689 |
| SZCCDBA067 | 62.5 | 99.57 | 1.17 | 1 | Yes | 0 | 3,923,379 | 3,923,379 | 3,747 |
| SZCCDBA088 | 62.4 | 99.57 | 1.41 | 18 | No | - | 3,865,049 | 423,953 | 3,700 |
| SZCCDBA151 | 62.4 | 99.57 | 1.25 | 26 | No | - | 4,034,501 | 552,978 | 3,880 |
| SZCCDBB009 | 62.4 | 99.57 | 1.57 | 15 | No | - | 3,841,971 | 525,579 | 3,683 |
| SZCCDBB020 | 62.5 | 99.57 | 1.41 | 2 | Yes | 1 | 3,855,144 | 3,813,992 | 3,693 |
| SZCCDBB057 | 62.5 | 99.57 | 0.93 | 16 | No | - | 3,791,328 | 541,733 | 3,630 |
| SZCCDBC058 | 62.6 | 99.57 | 1.17 | 2 | Yes | 1 | 4,127,254 | 3,979,580 | 3,979 |
| SZCCDBE041 | 62.3 | 99.57 | 1.17 | 35 | No | - | 4,074,834 | 389,130 | 3,891 |
| SZCCDBF062 | 62.4 | 99.57 | 1.25 | 19 | No | - | 3,838,335 | 375,792 | 3,667 |

[a] We cannot predict the number of plasmids in strains with only draft genome sequences available

**Table S2. Transcript levels of genes in the inorganic sulfur metabolism. Fold changes of enriched transcripts (log2 ratio >1 and adjust p value < 0.01) were highlighted in red, and depleted (log2 ratio < -1 and adjust p value < 0.01) in green.**

| Gene Locus | Gene name | Gene annotation (rast) | Pathway | Transcriptional responses in SZCCDBA067 | | | | | | | | Transcriptional responses in SZCCDBB020 | | | | | | | |
|---|---|---|---|---|---|---|---|---|---|---|---|---|---|---|---|---|---|---|---|
| | | | | log2 ratio (+thiosulfate/+sulfate) | | | | log2 ratio (microaerobic/aerobic) | | | | log2 ratio (+thiosulfate/+sulfate) | | | | log2 ratio (microaerobic/aerobic) | | | |
| | | | | Aerobic | adjust *p* | Microaerobic | adjust *p* | +sulfate | adjust *p* | +Thiosulfate | adjust *p* | Aerobic | adjust *p* | Microaerobic | adjust *p* | +sulfate | adjust *p* | +Thiosulfate | adjust *p* |
| SZCCDBA067_02258 | *pdo* | MBL-fold metallo-hydrolase superfamily | Sulfide oxidation | -0.22 | 1.09E-07 | 0.79 | 1.09E-07 | -0.05 | 1.09E-07 | 0.97 | 1.09E-07 | -0.40 | 6.89E-02 | 1.35 | 1.57E-15 | 0.66 | 9.38E-04 | 2.42 | 4.32E-46 |
| SZCCDBA067_01474 | *sqr* | Sulfide:quinone oxidoreductase, Type I | Sulfide oxidation | -0.42 | 5.08E-84 | 3.71 | 5.08E-84 | 0.50 | 5.08E-84 | 4.63 | 5.08E-84 | 0.62 | 1.57E-02 | 2.78 | 6.75E-78 | 1.12 | 1.29E-06 | 3.27 | 1.65E-91 |
| SZCCDBA067_02259 | *sqr* | Protein tyrosine phosphatase | Sulfide oxidation | -0.07 | 1.12E-02 | -0.50 | 1.12E-02 | 0.00 | 1.12E-02 | -0.43 | 1.12E-02 | -0.16 | 3.60E-01 | -0.30 | 3.06E-02 | 0.51 | 7.82E-04 | 0.37 | 6.15E-03 |
| SZCCDBA067_01326 | *tusA, sirA* | tRNA 5-methylaminomethyl-2-thiouridine synthesis sulfur carrier protein TusA | rDSR | 0.21 | 8.13E-01 | 0.04 | 8.13E-01 | 0.58 | 8.13E-01 | 0.41 | 8.13E-01 | 1.41 | 1.25E-03 | -0.22 | 5.33E-01 | 1.54 | 2.22E-04 | -0.09 | 7.99E-01 |
| SZCCDBA067_02158 | *dsrE2* | Peroxiredoxin family protein | rDSR | -0.21 | 2.70E-77 | 3.75 | 2.70E-77 | -1.30 | 2.70E-77 | 2.66 | 2.70E-77 | 2.31 | 2.82E-11 | 3.63 | 2.44E-87 | 2.08 | 1.05E-09 | 3.41 | 1.00E-75 |
| SZCCDBA067_02159 | *tusA, sirA* | tRNA 2-thiouridine synthesizing protein A | rDSR | -0.08 | 4.07E-41 | 4.03 | 4.07E-41 | 0.36 | 4.07E-41 | 4.47 | 4.07E-41 | 1.13 | 5.98E-02 | 2.72 | 1.77E-37 | 2.70 | 1.03E-07 | 4.29 | 8.15E-51 |
| SZCCDBA067_02160 | *dsrN* | Protein similar to cobyrinic acid a,c-diamide synthetase clustered with dissimilatory sulfite reductase | rDSR | -0.02 | 1.46E-11 | 1.86 | 1.46E-11 | -0.93 | 1.46E-11 | 0.94 | 1.46E-11 | -0.20 | 3.82E-01 | 1.94 | 2.07E-30 | -0.75 | 2.32E-04 | 1.40 | 7.06E-17 |
| SZCCDBA067_02161 | *Unknown* | Anthranilate phosphoribosyltransferase-like protein | rDSR | -0.34 | 1.90E-08 | 1.36 | 1.90E-08 | -0.93 | 1.90E-08 | 0.77 | 1.90E-08 | 0.27 | 2.22E-01 | 1.61 | 5.42E-26 | 0.13 | 5.50E-01 | 1.48 | 6.52E-22 |
| SZCCDBA067_02162 | *dsrP* | Sulfite reduction-associated complex DsrMKJOP protein DsrP (= HmeB) | rDSR | -0.01 | 4.63E-10 | 1.34 | 4.63E-10 | 0.03 | 4.63E-10 | 1.38 | 4.63E-10 | 0.43 | 1.09E-01 | 1.59 | 2.28E-19 | 1.09 | 3.54E-06 | 2.25 | 1.38E-34 |
| SZCCDBA067_02163 | *dsrO* | Sulfite reduction-associated complex DsrMKJOP iron-sulfur protein DsrO (=HmeA) | rDSR | -0.82 | 2.42E-14 | 3.08 | 2.42E-14 | -1.53 | 2.42E-14 | 2.37 | 2.42E-14 | 0.79 | 8.62E-02 | 3.77 | 4.22E-37 | 0.12 | 8.15E-01 | 3.10 | 1.27E-27 |
| SZCCDBA067_02165 | *dsrJ* | Protein similar to glutamate synthase [NADPH] small chain, clustered with sulfite reductase | rDSR | -0.01 | 1.12E-14 | 2.64 | 1.12E-14 | -0.22 | 1.12E-14 | 2.43 | 1.12E-14 | 0.65 | 2.39E-01 | 3.91 | 3.03E-38 | 0.63 | 2.20E-01 | 3.89 | 3.06E-35 |
| SZCCDBA067_02166 | *dsrK* | Sulfite reduction-associated complex DsrMKJOP protein DsrK (=HmeD) | rDSR | -0.06 | 4.13E-15 | 2.78 | 4.13E-15 | -0.75 | 4.13E-15 | 2.09 | 4.13E-15 | 1.12 | 2.73E-02 | 3.43 | 2.26E-28 | 0.72 | 1.52E-01 | 3.03 | 4.63E-23 |
| SZCCDBA067_02167 | *dsrM* | Sulfite reduction-associated complex DsrMKJOP protein DsrM (= HmeC) | rDSR | -0.26 | 1.24E-13 | 3.35 | 1.24E-13 | -1.63 | 1.24E-13 | 1.98 | 1.24E-13 | 2.05 | 1.72E-02 | 3.97 | 4.42E-22 | 1.70 | 4.38E-02 | 3.61 | 5.18E-19 |
| SZCCDBA067_02168 | *dsrC* | tRNA 2-thiouridine synthesis protein TusE @ Sulfur redox associated protein DsrC | rDSR | -0.40 | 6.56E-10 | 3.13 | 6.56E-10 | -0.91 | 6.56E-10 | 2.62 | 6.56E-10 | 1.90 | 2.80E-08 | 4.22 | 8.62E-89 | 0.69 | 6.17E-02 | 3.01 | 6.38E-56 |
| SZCCDBA067_02169 | *tusB, dsrH* | tRNA 5-methylaminomethyl-2-thiouridine synthase subunit TusB | rDSR | -0.85 | 6.79E-11 | 4.70 | 6.79E-11 | -2.08 | 6.79E-11 | 3.47 | 6.79E-11 | 3.33 | 2.41E-10 | 4.57 | 1.78E-53 | 0.94 | 1.13E-01 | 2.18 | 3.54E-21 |
| SZCCDBA067_02170 | *tusC, dsrF* | tRNA 5-methylaminomethyl-2-thiouridine synthase subunit TusC | rDSR | 0.21 | 1.47E-14 | 4.45 | 1.47E-14 | -0.88 | 1.47E-14 | 3.36 | 1.47E-14 | 1.07 | 1.07E-01 | 4.81 | 6.01E-54 | 1.07 | 8.26E-02 | 4.81 | 3.50E-48 |
| SZCCDBA067_02171 | *tusD, dsrE* | tRNA 5-methylaminomethyl-2-thiouridine synthase subunit TusD | rDSR | 0.64 | 2.00E-20 | 5.48 | 2.00E-20 | -1.42 | 2.00E-20 | 3.43 | 2.00E-20 | 0.03 | 9.72E-01 | 5.08 | 2.55E-64 | 0.79 | 1.77E-01 | 5.83 | 8.54E-54 |
| SZCCDBA067_02172 | *dsrB* | Dissimilatory sulfite reductase, beta subunit (EC 1.8.99.5) | rDSR | -0.40 | 1.98E-49 | 4.38 | 1.98E-49 | -0.61 | 1.98E-49 | 4.16 | 1.98E-49 | -0.20 | 6.90E-01 | 5.87 | 1.10E-136 | 0.21 | 6.37E-01 | 6.28 | 1.20E-113 |
| SZCCDBA067_02173 | *dsrA* | Dissimilatory sulfite reductase, alpha subunit (EC 1.8.99.5) | rDSR | -0.10 | 3.73E-78 | 5.73 | 3.73E-78 | -0.52 | 3.73E-78 | 5.31 | 3.73E-78 | 0.89 | 1.23E-01 | 6.21 | 1.12E-128 | 1.22 | 1.81E-02 | 6.54 | 2.59E-112 |
| SZCCDBA067_01242 | *cysH* | Phosphoadenylyl-sulfate reductase [thioredoxin] (EC 1.8.4.8) | Assimilatory sulfate reduction | -6.54 | 7.26E-191 | -4.28 | 7.26E-191 | -1.84 | 7.26E-191 | 0.42 | 7.26E-191 | -9.77 | 3.74E-96 | -4.93 | 6.12E-34 | -3.25 | 2.46E-13 | 1.59 | 4.81E-04 |
| SZCCDBA067_01243 | *cysI* | Sulfite reductase [NADPH] hemoprotein beta-component (EC 1.8.1.2) | Assimilatory sulfate reduction | -6.88 | 1.77E-129 | -4.10 | 1.77E-129 | -2.75 | 1.77E-129 | 0.03 | 1.77E-129 | -9.55 | 1.86E-94 | -4.71 | 6.24E-30 | -3.62 | 1.73E-15 | 1.22 | 6.06E-03 |
| SZCCDBA067_02374 | *sat, met3* | Sulfate adenylyltransferase (EC 2.7.7.4) / Adenylylsulfate kinase (EC 2.7.1.25) | Assimilatory sulfate reduction | -0.51 | 3.11E-19 | -1.12 | 3.11E-19 | -0.74 | 3.11E-19 | -1.35 | 3.11E-19 | -4.27 | 2.09E-302 | -1.38 | 1.70E-37 | -4.06 | 4.00E-276 | -1.17 | 2.63E-27 |
| SZCCDBA067_02936 | *soeA* | Sulfite dehydrogenase (quinone), molybdopterin-containing subunit SoeA (EC 1.8.5.6) | Sulfite oxidation | 0.17 | 2.19E-33 | 1.87 | 2.19E-33 | -0.49 | 2.19E-33 | 1.20 | 2.19E-33 | -0.12 | 4.06E-01 | 2.22 | 9.09E-103 | -0.89 | 2.41E-13 | 1.45 | 9.76E-46 |
| SZCCDBA067_02937 | *soeB* | Sulfite dehydrogenase (quinone), iron-sulfur subunit SoeB (EC 1.8.5.6) | Sulfite oxidation | 0.15 | 9.47E-20 | 1.83 | 9.47E-20 | -1.12 | 9.47E-20 | 0.57 | 9.47E-20 | 0.12 | 6.41E-01 | 2.61 | 1.86E-44 | -1.26 | 1.59E-08 | 1.23 | 2.35E-12 |
| SZCCDBA067_02938 | *soeC* | Sulfite dehydrogenase (quinone), membrane-anchor subunit SoeC (EC 1.8.5.6) | Sulfite oxidation | 0.21 | 9.60E-06 | 1.02 | 9.60E-06 | 0.40 | 9.60E-06 | 1.21 | 9.60E-06 | -0.02 | 9.57E-01 | 1.89 | 1.04E-29 | -0.55 | 8.76E-03 | 1.36 | 1.67E-16 |
| SZCCDBA067_00080 | *Rhodanese_1* | Rhodanese-like domain protein | | -0.30 | 4.36E-01 | -0.13 | 4.36E-01 | -0.42 | 4.36E-01 | -0.25 | 4.36E-01 | -0.64 | 6.49E-07 | -0.18 | 1.71E-01 | -1.74 | 8.41E-43 | -1.28 | 3.56E-28 |
| SZCCDBA067_00756 | *Rhodanese_3* | Rhodanese-like domain protein | | 0.06 | 8.50E-01 | -0.25 | 8.50E-01 | 0.26 | 8.50E-01 | -0.05 | 8.50E-01 | -0.43 | 4.97E-02 | 0.44 | 1.05E-02 | 0.05 | 8.32E-01 | 0.92 | 7.33E-08 |
| SZCCDBA067_00803 | *glpE* | Rhodanese-related sulfurtransferase | | 0.01 | 6.18E-01 | 0.12 | 6.18E-01 | 0.46 | 6.18E-01 | 0.56 | 6.18E-01 | 0.17 | 6.68E-01 | -0.17 | 5.41E-01 | 2.11 | 4.96E-12 | 1.77 | 1.52E-11 |
| SZCCDBA067_01184 | *Rhodanese_4* | Rhodanese-related sulfurtransferase | | -0.30 | 1.94E-01 | -0.54 | 1.94E-01 | 0.12 | 1.94E-01 | -0.12 | 1.94E-01 | -0.08 | 9.16E-01 | 0.23 | 7.01E-01 | -0.67 | 2.74E-01 | -0.36 | 5.00E-01 |
| SZCCDBA067_01274 | *Rhodanese_2* | Rhodanese-like domain protein | | 0.10 | 3.04E-17 | 2.45 | 3.04E-17 | 2.58 | 3.04E-17 | 4.93 | 3.04E-17 | -1.48 | 2.77E-03 | 3.90 | 5.29E-75 | 0.85 | 2.04E-02 | 6.24 | 3.61E-64 |
| SZCCDBA067_02201 | *tdhT* | Quino(hemo)protein alcohol dehydrogenase, PQQ-dependent (EC 1.1.2.8) | | 0.32 | 3.92E-08 | -1.41 | 3.92E-08 | -0.65 | 3.92E-08 | -2.39 | 3.92E-08 | 0.38 | 2.85E-01 | 0.50 | 7.92E-02 | -0.19 | 5.79E-01 | -0.07 | 8.22E-01 |
| SZCCDBA067_02466 | *TST* | Thiosulfate sulfurtransferase, rhodanese (EC 2.8.1.1) | | -0.29 | 1.96E-14 | -1.14 | 1.96E-14 | -0.86 | 1.96E-14 | -1.70 | 1.96E-14 | -0.01 | 9.81E-01 | -0.46 | 9.72E-02 | 0.47 | 1.51E-01 | 0.02 | 9.61E-01 |
| SZCCDBA067_03117 | *TST2* | 3-mercaptopyruvate sulfurtransferase (EC 2.8.1.2) | | -0.08 | 5.22E-01 | -0.21 | 5.22E-01 | -0.12 | 5.22E-01 | -0.25 | 5.22E-01 | -0.44 | 2.28E-03 | -0.44 | 4.91E-04 | -0.23 | 1.18E-01 | -0.23 | 7.22E-02 |

**Table S3. Transcript levels of genes in denitrification and assimilatory nitrate reduction. Fold changes of enriched transcripts (log2 ratio >1 and adjust p value < 0.01) were highlighted in red, and depleted (log2 ratio < -1 and adjust p value < 0.01) in green**

| Gene Locus | Gene name | Gene annotation (rast) | Pathway | Transcriptional responses in SZCCDBA067 | | | | | | | | Transcriptional responses in SZCCDBB020 | | | | | | | |
|---|---|---|---|---|---|---|---|---|---|---|---|---|---|---|---|---|---|---|---|
| | | | | log2 ratio (+thiosulfate/-thiosulfate) | | | | log2 ratio (microaerobic/aerobic) | | | | log2 ratio (+thiosulfate/-thiosulfate) | | | | log2 ratio (microaerobic/aerobic) | | | |
| | | | | Aerobic | adjust *p* | Microaerobic | adjust *p* | -Thiosulfa | adjust *p* | +Thiosulfate | adjust *p* | Aerobic | adjust *p* | Microaerobic | adjust *p* | -Thiosulfate | adjust *p* | +Thiosulfate | adjust *p* |
| SZCCDBA067_00029 | *nirN* | Nitrite reductase associated c-type cytochorome NirN | nitrite reduction | -1.16 | 2.44E-10 | **1.29** | 2.44E-10 | -4.31 | 2.44E-10 | -1.86 | 2.44E-10 | 0.57 | 5.25E-03 | **1.11** | 2.40E-10 | -2.86 | 2.98E-52 | -2.31 | 5.13E-43 |
| SZCCDBA067_00030 | *nirJ* | Heme d1 biosynthesis protein NirJ | nitrite reduction | -1.20 | 4.72E-22 | **2.11** | 4.72E-22 | -5.39 | 4.72E-22 | -2.08 | 4.72E-22 | 0.66 | 6.69E-05 | **1.21** | 7.37E-17 | -3.39 | 4.46E-105 | -2.83 | 6.46E-93 |
| SZCCDBA067_00031 | *nirH* | Heme d1 biosynthesis protein NirH | nitrite reduction | -0.55 | 2.86E-44 | **1.51** | 2.86E-44 | -5.36 | 2.86E-44 | -3.30 | 2.86E-44 | 0.70 | 1.72E-04 | 0.92 | 4.91E-08 | -3.30 | 5.56E-77 | -3.08 | 3.00E-85 |
| SZCCDBA067_00032 | *nirG* | Heme d1 biosynthesis protein NirG | nitrite reduction | -0.45 | 6.54E-43 | **1.51** | 6.54E-43 | -4.92 | 6.54E-43 | -2.97 | 6.54E-43 | 0.67 | 7.01E-05 | **1.43** | 2.66E-20 | -3.66 | 1.48E-109 | -2.91 | 4.58E-91 |
| SZCCDBA067_00033 | *nirD/L* | Heme d1 biosynthesis protein NirD / Heme d1 biosynthesis protein NirL | nitrite reduction | -0.31 | 1.89E-21 | **1.64** | 1.89E-21 | -4.64 | 1.89E-21 | -2.69 | 1.89E-21 | 0.88 | 1.92E-04 | **1.55** | 9.70E-14 | -3.20 | 2.45E-46 | -2.53 | 1.00E-36 |
| SZCCDBA067_00034 | *nirF* | Heme d1 biosynthesis protein NirF | nitrite reduction | -0.24 | 8.48E-31 | **1.32** | 8.48E-31 | -5.37 | 8.48E-31 | -3.81 | 8.48E-31 | 0.67 | 2.63E-04 | **1.52** | 1.82E-21 | -3.67 | 2.30E-100 | -2.82 | 4.69E-75 |
| SZCCDBA067_00035 | *nirC* | Cytochrome c55X precursor NirC | nitrite reduction | -0.28 | 7.29E-41 | 0.57 | 7.29E-41 | -5.06 | 7.29E-41 | -4.21 | 7.29E-41 | 0.56 | 2.06E-04 | **1.06** | 2.81E-14 | -3.84 | 1.67E-153 | -3.34 | 9.30E-152 |
| SZCCDBA067_00036 | *nirE* | Uroporphyrinogen-III methyltransferase (EC 2.1.1.107) | nitrite reduction | -0.25 | 6.74E-27 | -0.24 | 6.74E-27 | -4.02 | 6.74E-27 | -4.01 | 6.74E-27 | 0.53 | 8.37E-05 | 0.56 | 5.67E-06 | -3.24 | 1.62E-141 | -3.21 | 2.49E-175 |
| SZCCDBA067_00039 | *nirS* | Nitrite reductase (EC 1.7.2.1) | nitrite reduction | -0.87 | 2.60E-21 | 0.32 | 2.60E-21 | -4.62 | 2.60E-21 | -3.43 | 2.60E-21 | -0.40 | 1.14E-02 | 0.60 | 8.60E-06 | -3.62 | 1.88E-143 | -2.62 | 2.22E-94 |
| SZCCDBA067_02095 | *nirS_2* | Nitrite reductase (EC 1.7.2.1) | nitrite reduction | -0.48 | 5.78E-03 | **1.15** | 5.78E-03 | -1.34 | 5.78E-03 | 0.30 | 5.78E-03 | 0.41 | 6.96E-02 | 0.75 | 2.31E-05 | 0.92 | 6.82E-06 | **1.25** | 4.19E-13 |
| SZCCDBA067_00040 | *NnrS* | NnrS protein involved in response to NO | NO reduction | -0.64 | 1.34E-17 | **1.10** | 1.34E-17 | -4.93 | 1.34E-17 | -3.19 | 1.34E-17 | 0.35 | 2.31E-02 | 0.25 | 8.91E-02 | -2.69 | 4.13E-79 | -2.79 | 1.12E-107 |
| SZCCDBA067_00041 | *norC* | Nitric-oxide reductase subunit C (EC 1.7.99.7) | NO reduction | -0.66 | 2.63E-36 | -1.47 | 2.63E-36 | -4.56 | 2.63E-36 | -5.37 | 2.63E-36 | -0.51 | 1.68E-03 | -0.40 | 7.61E-03 | -3.90 | 5.03E-145 | -3.79 | 1.52E-169 |
| SZCCDBA067_00042 | *norB* | Nitric-oxide reductase subunit B (EC 1.7.99.7) | NO reduction | -0.78 | 2.68E-45 | 0.26 | 2.68E-45 | -5.65 | 2.68E-45 | -4.61 | 2.68E-45 | -0.61 | 6.45E-05 | 0.40 | 5.26E-03 | -4.91 | 1.10E-252 | -3.90 | 2.60E-199 |
| SZCCDBA067_00043 | *norQ* | Nitric oxide reductase activation protein NorQ | NO reduction | -0.62 | 9.70E-110 | **1.74** | 9.70E-110 | -6.70 | 9.70E-110 | -4.35 | 9.70E-110 | 0.06 | 8.00E-01 | **1.06** | 3.18E-09 | -4.48 | 1.55E-125 | -3.49 | 3.34E-96 |
| SZCCDBA067_00044 | *norD* | Nitric oxide reductase activation protein NorD | NO reduction | -0.98 | 3.66E-70 | **2.04** | 3.66E-70 | -6.36 | 3.66E-70 | -3.34 | 3.66E-70 | 0.54 | 4.12E-02 | **1.24** | 2.10E-08 | -3.78 | 5.19E-58 | -3.08 | 2.75E-48 |
| SZCCDBA067_00045 | *norE* | Nitric oxide reductase activation protein NorE | NO reduction | -2.47 | 2.38E-16 | **1.36** | 2.38E-16 | -5.07 | 2.38E-16 | -1.24 | 2.38E-16 | 0.81 | 9.38E-08 | **1.55** | 1.88E-24 | -3.17 | 8.81E-88 | -2.44 | 1.03E-75 |
| SZCCDBA067_00052 | *nosR* | Nitrous oxide reductase maturation protein NosR | nitrous oxide reduction | -0.42 | 1.09E-55 | 0.87 | 1.09E-55 | -5.09 | 1.09E-55 | -3.80 | 1.09E-55 | 0.89 | 2.76E-08 | 0.62 | 2.38E-05 | -3.19 | 8.73E-95 | -3.46 | 2.03E-139 |
| SZCCDBA067_00053 | *nosZ* | Nitrous-oxide reductase (EC 1.7.99.6) | nitrous oxide reduction | -0.85 | 1.66E-17 | 0.56 | 1.66E-17 | -4.09 | 1.66E-17 | -2.67 | 1.66E-17 | 0.03 | 9.17E-01 | -0.14 | 5.21E-01 | -1.71 | 3.12E-17 | -1.88 | 3.08E-25 |
| SZCCDBA067_00054 | *nosD* | Nitrous oxide reductase maturation protein NosD | nitrous oxide reduction | -1.42 | 2.25E-15 | 0.98 | 2.25E-15 | -4.18 | 2.25E-15 | -1.77 | 2.25E-15 | 0.74 | 2.35E-03 | 0.35 | 1.20E-01 | -2.02 | 4.71E-19 | -2.41 | 8.31E-33 |
| SZCCDBA067_00055 | *nosF* | Nitrous oxide reductase maturation protein NosF (ATPase) | nitrous oxide reduction | -1.73 | 2.01E-09 | **1.08** | 2.01E-09 | -3.56 | 2.01E-09 | -0.76 | 2.01E-09 | 0.79 | 1.25E-04 | 0.47 | 1.38E-02 | -1.70 | 3.81E-18 | -2.02 | 3.21E-31 |
| SZCCDBA067_00056 | *nosY* | Nitrous oxide reductase maturation transmembrane protein NosY | nitrous oxide reduction | -1.42 | 2.09E-05 | **1.20** | 2.09E-05 | -3.28 | 2.09E-05 | -0.66 | 2.09E-05 | -0.01 | 9.77E-01 | 0.20 | 4.81E-01 | -1.61 | 3.91E-09 | -1.40 | 1.30E-08 |
| SZCCDBA067_00057 | *nosL* | Nitrous oxide reductase maturation protein, outer-membrane lipoprotein NosL | nitrous oxide reduction | -1.26 | 2.00E-03 | 0.92 | 2.00E-03 | -3.78 | 2.00E-03 | -1.59 | 2.00E-03 | -0.88 | 9.51E-04 | 0.33 | 1.96E-01 | -2.04 | 5.24E-16 | -0.83 | 3.26E-04 |
| SZCCDBA067_01137 | *ftrB* | Transcriptional regulator NarR | nitrate reduction | 0.64 | 1.24E-04 | -0.39 | 1.24E-04 | **1.88** | 1.24E-04 | 0.85 | 1.24E-04 | -1.46 | 9.27E-17 | -0.51 | 7.17E-04 | 0.79 | 2.27E-06 | **1.73** | 7.49E-30 |
| SZCCDBA067_01138 | *NRT, narK, nrtP, nasA* | Nitrate/nitrite transporter NarK/U 1 / Nitrate/nitrite transporter NarK/U | nitrate reduction | -0.38 | 7.87E-40 | **2.62** | 7.87E-40 | -6.30 | 7.87E-40 | -3.30 | 7.87E-40 | **2.37** | 2.35E-04 | **2.08** | 3.01E-04 | -3.39 | 4.26E-08 | -3.67 | 2.87E-11 |
| SZCCDBA067_01139 | *narG, narZ, nxrA* | Respiratory nitrate reductase alpha chain (EC 1.7.99.4) | nitrate reduction | -0.80 | 2.75E-12 | **1.46** | 2.75E-12 | -4.09 | 2.75E-12 | -1.84 | 2.75E-12 | **1.31** | 4.08E-02 | **1.11** | 4.84E-02 | -1.75 | 3.74E-03 | -1.95 | 2.67E-04 |
| SZCCDBA067_01140 | *narH, narY, nxrB* | Respiratory nitrate reductase beta chain (EC 1.7.99.4) | nitrate reduction | -1.14 | 1.55E-04 | **1.11** | 1.55E-04 | -3.10 | 1.55E-04 | -0.86 | 1.55E-04 | 0.07 | 9.26E-01 | 0.49 | 3.84E-01 | -0.90 | 1.28E-01 | -0.48 | 3.71E-01 |
| SZCCDBA067_01141 | *narJ, narW* | Respiratory nitrate reductase delta chain (EC 1.7.99.4) | nitrate reduction | -0.85 | 2.47E-03 | **1.70** | 2.47E-03 | -2.56 | 2.47E-03 | -0.01 | 2.47E-03 | -1.00 | 5.92E-02 | 0.38 | 4.51E-01 | -0.68 | 1.92E-01 | 0.70 | 1.26E-01 |
| SZCCDBA067_01142 | *narI, narV* | Respiratory nitrate reductase gamma chain (EC 1.7.99.4) | nitrate reduction | -1.02 | 2.50E-02 | 0.90 | 2.50E-02 | -1.65 | 2.50E-02 | 0.26 | 2.50E-02 | -1.78 | 5.55E-09 | 0.33 | 2.69E-01 | -0.45 | 1.60E-01 | **1.66** | 7.99E-10 |
| SZCCDBA067_01395 | *nasT* | two-component system, response regulator / RNA-binding antiterminator | Assimilatory nitrate reduction | 0.12 | 1.39E-14 | -1.02 | 1.39E-14 | **3.03** | 1.39E-14 | **1.90** | 1.39E-14 | 0.26 | 3.89E-01 | -0.57 | 1.09E-02 | **1.80** | 1.14E-12 | 0.96 | 1.78E-05 |
| SZCCDBA067_01396 | *nasS* | two-component system, oxyanion-binding sensor | Assimilatory nitrate reduction | 0.25 | 3.37E-14 | -2.22 | 3.37E-14 | **3.31** | 3.37E-14 | 0.85 | 3.37E-14 | 0.27 | 1.80E-01 | -0.37 | 1.26E-02 | 0.99 | 8.58E-09 | 0.35 | 2.40E-02 |
| SZCCDBA067_01397 | *nrtA, nasF, cynA* | Nitrate ABC transporter, substrate-binding protein | Assimilatory nitrate reduction | 0.20 | 8.45E-72 | -2.65 | 8.45E-72 | **4.89** | 8.45E-72 | **2.04** | 8.45E-72 | -0.23 | 5.23E-01 | -1.22 | 4.42E-09 | **3.44** | 5.46E-39 | **2.44** | 1.01E-25 |
| SZCCDBA067_01398 | *nrtB, nasE, cynB* | Nitrate ABC transporter, permease protein | Assimilatory nitrate reduction | 0.09 | 1.48E-25 | -3.54 | 1.48E-25 | **4.83** | 1.48E-25 | **1.20** | 1.48E-25 | -0.29 | 5.47E-01 | -1.35 | 7.75E-06 | **2.18** | 2.03E-09 | **1.12** | 8.10E-04 |
| SZCCDBA067_01399 | *nrtC, nasD* | Nitrate ABC transporter, ATP-binding protein | Assimilatory nitrate reduction | -0.06 | 4.29E-05 | -1.71 | 4.29E-05 | **2.24** | 4.29E-05 | 0.59 | 4.29E-05 | -0.51 | 1.34E-01 | -0.17 | 5.43E-01 | 0.80 | 7.01E-03 | **1.14** | 1.32E-05 |
| SZCCDBA067_01400 | *nirB* | Nitrite reductase [NAD(P)H] large subunit (EC 1.7.1.4) | Assimilatory nitrate reduction | 0.23 | 5.81E-04 | -0.79 | 5.81E-04 | **1.94** | 5.81E-04 | 0.92 | 5.81E-04 | -0.47 | 1.38E-01 | 0.47 | 6.82E-02 | 0.20 | 5.19E-01 | **1.14** | 4.19E-06 |
| SZCCDBA067_01401 | *nirD* | Nitrite reductase [NAD(P)H] small subunit (EC 1.7.1.4) | Assimilatory nitrate reduction | 0.02 | 2.18E-01 | -0.42 | 2.18E-01 | 1.63 | 2.18E-01 | 1.19 | 2.18E-01 | -0.05 | 9.41E-01 | 0.39 | 3.84E-01 | 0.09 | 8.86E-01 | 0.53 | 2.23E-01 |
| SZCCDBA067_01402 | *nasA* | assimilatory nitrate reductase catalytic subunit | Assimilatory nitrate reduction | 0.00 | 3.53E-02 | 0.30 | 3.53E-02 | 0.38 | 3.53E-02 | 0.69 | 3.53E-02 | 0.56 | 5.59E-02 | 0.73 | 1.73E-03 | 0.67 | 1.49E-02 | 0.83 | 2.54E-04 |

Table S4. Transcript levels of genes in aerobic respiration. Fold changes of enriched transcripts (log2 ratio >1 and adjust p value < 0.01) were highlighted in red, and depleted (log2 ratio < -1 and adjust p value < 0.01) in green.

| Gene Locus | Gene name | Gene annotation (rast) | Pathway | Transcriptional responses in SZCCDBA067 | | | | | | | | Transcriptional responses in SZCCDBB020 | | | | | | | |
|---|---|---|---|---|---|---|---|---|---|---|---|---|---|---|---|---|---|---|---|
| | | | | log2 ratio (+thiosulfate/-thiosulfate) | | | | log2 ratio (microaerobic/aerobic) | | | | log2 ratio (+thiosulfate/-thiosulfate) | | | | log2 ratio (microaerobic/aerobic) | | | |
| | | | | Aerobic | adjust *p* | Microaerobic | adjust *p* | -Thiosulfate | adjust *p* | +Thiosulfate | adjust *p* | Aerobic | adjust *p* | Microaerobic | adjust *p* | -Thiosulfate | adjust *p* | +Thiosulfate | adjust *p* |
| SZCCDBA067_02005 | *ccmC* | Cytochrome c-type biogenesis protein CcmC, putative heme lyase for CcmE | ABC transporter (Heme) | -0.57 | 7.91E-14 | 0.77 | 7.91E-14 | -3.93 | 7.91E-14 | -2.59 | 7.91E-14 | 0.62 | 1.82E-06 | -0.02 | 8.66E-01 | -1.28 | 1.61E-24 | -1.92 | 1.07E-67 |
| SZCCDBA067_02006 | *ccmB* | ABC transporter involved in cytochrome c biogenesis, CcmB subunit | ABC transporter (Heme) | 0.16 | 1.62E-01 | 0.85 | 1.62E-01 | -0.66 | 1.62E-01 | 0.03 | 1.62E-01 | 0.19 | 1.85E-01 | 0.04 | 7.67E-01 | -1.07 | 8.79E-17 | -1.22 | 2.86E-27 |
| SZCCDBA067_02007 | *ccmA* | ABC transporter involved in cytochrome c biogenesis, ATPase component CcmA | ABC transporter (Heme) | 0.25 | 4.37E-01 | -0.14 | 4.37E-01 | 0.22 | 4.37E-01 | -0.17 | 4.37E-01 | 0.04 | 7.82E-01 | -0.36 | 7.25E-04 | -0.30 | 1.07E-02 | -0.70 | 4.56E-12 |
| SZCCDBA067_00979 | *nuoN* | NADH-ubiquinone oxidoreductase chain N (EC 1.6.5.3) | Complex I: NADH dehydrogenase | -0.05 | 1.08E-04 | 1.24 | 1.08E-04 | -1.30 | 1.08E-04 | -0.01 | 1.08E-04 | 0.14 | 2.56E-01 | -0.28 | 6.39E-03 | -0.42 | 1.10E-04 | -0.83 | 9.77E-19 |
| SZCCDBA067_00978 | *nuoM* | NADH-ubiquinone oxidoreductase chain M (EC 1.6.5.3) | Complex I: NADH dehydrogenase | -0.05 | 6.10E-09 | 0.98 | 6.10E-09 | -1.51 | 6.10E-09 | -0.48 | 6.10E-09 | 0.33 | 5.25E-02 | -0.28 | 6.33E-02 | -0.70 | 8.32E-06 | -1.32 | 1.85E-21 |
| SZCCDBA067_00977 | *nuoL* | NADH-ubiquinone oxidoreductase chain L (EC 1.6.5.3) | Complex I: NADH dehydrogenase | -0.08 | 2.40E-10 | 1.05 | 2.40E-10 | -1.81 | 2.40E-10 | -0.68 | 2.40E-10 | 0.24 | 1.06E-01 | -0.10 | 4.74E-01 | -0.99 | 1.26E-13 | -1.34 | 2.62E-29 |
| SZCCDBA067_00976 | *nuoK* | NADH-ubiquinone oxidoreductase chain K (EC 1.6.5.3) | Complex I: NADH dehydrogenase | -0.12 | 9.68E-05 | 1.30 | 9.68E-05 | -2.35 | 9.68E-05 | -0.93 | 9.68E-05 | 0.09 | 5.37E-01 | -0.42 | 1.59E-04 | -0.96 | 1.09E-16 | -1.46 | 4.83E-46 |
| SZCCDBA067_00975 | *nuoJ* | NADH-ubiquinone oxidoreductase chain J (EC 1.6.5.3) | Complex I: NADH dehydrogenase | -0.08 | 6.01E-04 | 0.90 | 6.01E-04 | -1.89 | 6.01E-04 | -0.91 | 6.01E-04 | 0.22 | 8.02E-02 | -0.60 | 1.02E-08 | -0.66 | 5.17E-09 | -1.48 | 4.89E-50 |
| SZCCDBA067_00973 | *nuoI* | NADH-ubiquinone oxidoreductase chain I (EC 1.6.5.3) | Complex I: NADH dehydrogenase | -0.16 | 1.08E-08 | 0.61 | 1.08E-08 | -1.94 | 1.08E-08 | -1.17 | 1.08E-08 | 0.58 | 1.50E-06 | -0.35 | 2.01E-03 | -0.65 | 6.36E-08 | -1.58 | 5.41E-52 |
| SZCCDBA067_00972 | *nuoH* | NADH-ubiquinone oxidoreductase chain H (EC 1.6.5.3) | Complex I: NADH dehydrogenase | -0.08 | 2.58E-05 | 0.97 | 2.58E-05 | -2.20 | 2.58E-05 | -1.15 | 2.58E-05 | 0.65 | 1.59E-04 | -0.34 | 3.10E-02 | -0.50 | 3.08E-03 | -1.49 | 1.36E-24 |
| SZCCDBA067_00970 | *nuoG* | NADH-ubiquinone oxidoreductase chain G (EC 1.6.5.3) | Complex I: NADH dehydrogenase | 0.04 | 1.15E-19 | 1.17 | 1.15E-19 | -1.68 | 1.15E-19 | -0.55 | 1.15E-19 | 0.76 | 1.45E-07 | 0.11 | 4.73E-01 | -0.70 | 6.87E-07 | -1.35 | 1.25E-27 |
| SZCCDBA067_00966 | *nuoF* | NADH-ubiquinone oxidoreductase chain F (EC 1.6.5.3) | Complex I: NADH dehydrogenase | 0.12 | 5.99E-16 | 1.00 | 5.99E-16 | -1.12 | 5.99E-16 | -0.24 | 5.99E-16 | 0.24 | 1.99E-01 | -0.02 | 9.18E-01 | -0.64 | 1.21E-04 | -0.89 | 9.26E-10 |
| SZCCDBA067_00964 | *nuoE* | NADH-ubiquinone oxidoreductase chain E (EC 1.6.5.3) | Complex I: NADH dehydrogenase | 0.01 | 1.42E-02 | 0.96 | 1.42E-02 | -1.18 | 1.42E-02 | -0.23 | 1.42E-02 | -0.40 | 9.23E-04 | 0.17 | 1.43E-01 | -1.66 | 3.59E-49 | -1.09 | 4.24E-27 |
| SZCCDBA067_00963 | *nuoD* | NADH-ubiquinone oxidoreductase chain D (EC 1.6.5.3) | Complex I: NADH dehydrogenase | -0.15 | 6.04E-25 | 2.00 | 6.04E-25 | -2.23 | 6.04E-25 | -0.08 | 6.04E-25 | -0.52 | 7.54E-03 | 0.38 | 2.90E-02 | -1.60 | 5.88E-19 | -0.70 | 2.22E-05 |
| SZCCDBA067_00960 | *nuoC* | NADH-ubiquinone oxidoreductase chain C (EC 1.6.5.3) | Complex I: NADH dehydrogenase | -0.09 | 3.44E-03 | 0.24 | 3.44E-03 | -1.54 | 3.44E-03 | -1.21 | 3.44E-03 | -1.11 | 2.91E-08 | -0.25 | 1.98E-01 | -1.91 | 9.60E-23 | -1.05 | 3.25E-09 |
| SZCCDBA067_00959 | *nuoB* | NADH-ubiquinone oxidoreductase chain B (EC 1.6.5.3) | Complex I: NADH dehydrogenase | -0.20 | 1.37E-02 | -0.22 | 1.37E-02 | -1.07 | 1.37E-02 | -1.10 | 1.37E-02 | -0.72 | 1.62E-06 | -0.67 | 8.03E-07 | -0.84 | 1.16E-08 | -0.79 | 2.97E-09 |
| SZCCDBA067_00958 | *nuoA* | NADH ubiquinone oxidoreductase chain A (EC 1.6.5.3) | Complex I: NADH dehydrogenase | -0.11 | 1.74E-01 | 0.07 | 1.74E-01 | -1.29 | 1.74E-01 | -1.12 | 1.74E-01 | -0.46 | 3.98E-03 | -0.82 | 2.02E-09 | -0.54 | 4.45E-04 | -0.90 | 1.81E-11 |
| SZCCDBA067_02279 | *Unknown* | NADH:ubiquinone oxidoreductase 17.2 kD subunit | Complex I: NADH dehydrogenase | -0.06 | 1.89E-01 | -0.76 | 1.89E-01 | 0.17 | 1.89E-01 | -0.53 | 1.89E-01 | -0.05 | 7.88E-01 | -0.41 | 1.83E-03 | -0.13 | 4.24E-01 | -0.48 | 1.35E-04 |
| SZCCDBA067_01108 | *fdoI, fdsG* | Formate dehydrogenase -O, gamma subunit (EC 1.2.1.2) | Complex II: formate dehydrogenase | -0.30 | 2.92E-01 | 0.16 | 2.92E-01 | 0.40 | 2.92E-01 | 0.86 | 2.92E-01 | 0.49 | 1.99E-02 | 0.43 | 7.93E-03 | 0.64 | 1.08E-03 | 0.58 | 2.83E-04 |
| SZCCDBA067_01111 | *fdoG, fdhF, fdwA* | Formate dehydrogenase-O, major subunit (EC 1.2.1.2) | Complex II: formate dehydrogenase | -0.69 | 3.84E-03 | 0.30 | 3.84E-03 | 1.25 | 3.84E-03 | 2.24 | 3.84E-03 | 2.17 | 5.27E-27 | 1.05 | 3.48E-12 | 1.85 | 3.57E-20 | 0.73 | 1.36E-06 |
| SZCCDBA067_00500 | *sdhD, frdD* | Succinate dehydrogenase hydrophobic membrane anchor protein | Complex II: succinate dehydrogenase | 0.34 | 6.31E-01 | -0.11 | 6.31E-01 | 0.64 | 6.31E-01 | 0.19 | 6.31E-01 | 1.09 | 1.04E-05 | -0.94 | 3.78E-06 | 3.04 | 2.54E-39 | 1.01 | 7.10E-07 |
| SZCCDBA067_00499 | *sdhC, frdC* | Succinate dehydrogenase cytochrome b-556 subunit | Complex II: succinate dehydrogenase | 0.29 | 8.23E-02 | -0.11 | 8.23E-02 | 1.39 | 8.23E-02 | 0.98 | 8.23E-02 | 1.23 | 3.86E-08 | -0.99 | 2.50E-07 | 3.76 | 3.26E-70 | 1.55 | 1.70E-16 |
| SZCCDBA067_00502 | *sdhB, frdB* | Succinate dehydrogenase iron-sulfur protein (EC 1.3.5.1) | Complex II: succinate dehydrogenase | 0.35 | 5.73E-07 | 1.60 | 5.73E-07 | -0.82 | 5.73E-07 | 0.43 | 5.73E-07 | 0.91 | 1.24E-04 | 0.08 | 7.53E-01 | 0.87 | 2.02E-04 | 0.03 | 8.95E-01 |
| SZCCDBA067_03542 | *UQCRFS1, RIP1, petA* | Ubiquinol-cytochrome C reductase iron-sulfur subunit (EC 1.10.2.2) | Complex III: Cytochrome c reductase | -0.04 | 1.44E-14 | -1.20 | 1.44E-14 | 0.50 | 1.44E-14 | -0.67 | 1.44E-14 | -0.30 | 1.70E-01 | -0.95 | 5.84E-08 | 0.01 | 9.50E-01 | -0.64 | 2.41E-04 |
| SZCCDBA067_01362 | *qcrC* | hypothetical protein | Complex III: Cytochrome c reductase | 0.26 | 1.05E-06 | 1.10 | 1.05E-06 | 0.34 | 1.05E-06 | 1.18 | 1.05E-06 | 0.79 | 7.66E-03 | 0.13 | 6.14E-01 | 0.96 | 6.43E-04 | 0.29 | 2.13E-01 |
| SZCCDBA067_03543 | *CYTB, petB* | Ubiquinol-cytochrome C reductase, cytochrome B subunit (EC 1.10.2.2) | Complex III: Cytochrome c reductase | -0.06 | 2.82E-02 | -0.46 | 2.82E-02 | -0.86 | 2.82E-02 | -1.27 | 2.82E-02 | -1.22 | 5.43E-13 | -0.90 | 5.08E-09 | -1.75 | 3.57E-26 | -1.44 | 7.93E-22 |
| SZCCDBA067_03544 | *CYC1, CYT1, petC* | Ubiquinol-cytochrome C reductase, cytochrome C1 subunit | Complex III: Cytochrome c reductase | -0.06 | 3.96E-05 | 0.12 | 3.96E-05 | -1.30 | 3.96E-05 | -1.12 | 3.96E-05 | -1.28 | 2.01E-11 | -0.51 | 4.03E-03 | -2.15 | 4.57E-31 | -1.39 | 1.53E-16 |
| SZCCDBA067_00255 | *coxC, ctaE* | Cytochrome c oxidase polypeptide III (EC 1.9.3.1) | Complex IV: Cytochrome c oxidase | 0.32 | 4.84E-01 | 0.19 | 4.84E-01 | 0.10 | 4.84E-01 | -0.03 | 4.84E-01 | -1.87 | 8.46E-12 | -1.05 | 2.67E-05 | -0.92 | 9.08E-04 | -0.11 | 6.99E-01 |
| SZCCDBA067_00251 | *coxB, ctaC* | Cytochrome c oxidase polypeptide II (EC 1.9.3.1) | Complex IV: Cytochrome c oxidase | 0.29 | 7.31E-03 | -0.20 | 7.31E-03 | 1.60 | 7.31E-03 | 1.11 | 7.31E-03 | -1.51 | 2.96E-27 | -0.58 | 9.27E-06 | 0.03 | 8.65E-01 | 0.97 | 1.29E-14 |
| SZCCDBA067_02063 | *coxB, ctaC* | Cytochrome c oxidase (B(O/a)3-type) chain II (EC 1.9.3.1) | Complex IV: Cytochrome c oxidase | -0.34 | 2.51E-04 | 0.95 | 2.51E-04 | -1.31 | 2.51E-04 | -0.02 | 2.51E-04 | 1.53 | 1.77E-11 | 1.23 | 9.77E-10 | 0.39 | 1.02E-01 | 0.10 | 6.61E-01 |
| SZCCDBA067_02062 | *coxA, ctaD* | Cytochrome c oxidase (B(O/a)3-type) chain I (EC 1.9.3.1) | Complex IV: Cytochrome c oxidase | -0.31 | 2.22E-10 | 1.17 | 2.22E-10 | -0.59 | 2.22E-10 | 0.90 | 2.22E-10 | 0.49 | 2.92E-03 | 1.26 | 1.24E-19 | 0.05 | 7.77E-01 | 0.82 | 2.99E-09 |
| SZCCDBA067_02431 | *coxA, ctaD* | Cytochrome c oxidase polypeptide I (EC 1.9.3.1) | Complex IV: Cytochrome c oxidase | 0.47 | 1.99E-04 | -0.63 | 1.99E-04 | 2.54 | 1.99E-04 | 1.43 | 1.99E-04 | -3.45 | 8.27E-56 | -0.93 | 6.10E-06 | -0.95 | 2.68E-05 | 1.57 | 2.35E-15 |
| SZCCDBA067_01462 | *COX15, ctaA* | Heme A synthase, cytochrome oxidase biogenesis protein Cox15-CtaA | Complex IV: Cytochrome c oxidase | -0.11 | 2.26E-01 | 0.21 | 2.26E-01 | -0.59 | 2.26E-01 | -0.28 | 2.26E-01 | -0.75 | 1.88E-05 | 0.14 | 4.30E-01 | -1.00 | 3.81E-09 | -0.11 | 5.24E-01 |
| SZCCDBA067_00254 | *COX11, ctaG* | Cytochrome oxidase biogenesis protein Cox11-CtaG, copper delivery to Cox1 | Complex IV: Cytochrome c oxidase | 0.35 | 2.70E-03 | 0.63 | 2.70E-03 | 0.56 | 2.70E-03 | 0.84 | 2.70E-03 | -1.90 | 1.14E-18 | -0.30 | 1.50E-01 | -0.98 | 6.36E-06 | 0.62 | 1.79E-03 |
| SZCCDBA067_00252 | *COX10, ctaB, cyoE* | Heme O synthase, protoheme IX farnesyltransferase, COX10-CtaB | Complex IV: Cytochrome c oxidase | 0.27 | 7.45E-05 | 0.43 | 7.45E-05 | 1.61 | 7.45E-05 | 1.77 | 7.45E-05 | -1.62 | 4.25E-16 | -0.47 | 1.22E-02 | -0.06 | 8.17E-01 | 1.09 | 9.39E-10 |
| SZCCDBA067_02061 | *COX10, ctaB, cyoE* | Heme O synthase, protoheme IX farnesyltransferase, COX10-CtaB | Complex IV: Cytochrome c oxidase | -0.81 | 9.59E-17 | 1.66 | 9.59E-17 | -0.51 | 9.59E-17 | 1.96 | 9.59E-17 | 0.15 | 5.03E-01 | 1.18 | 2.85E-12 | 0.59 | 2.39E-03 | 1.62 | 1.43E-22 |
| SZCCDBA067_02894 | *ccoQ* | Cytochrome c oxidase (cbb3-type) subunit CcoQ (EC 1.9.3.1) | Complex IV: Cytochrome c oxidase (cbb3) | -0.32 | 1.33E-11 | 1.37 | 1.33E-11 | -4.49 | 1.33E-11 | -2.80 | 1.33E-11 | 1.51 | 1.26E-10 | 0.92 | 2.75E-05 | -2.16 | 1.18E-20 | -2.76 | 1.89E-41 |
| SZCCDBA067_02893 | *ccoP* | Cytochrome c oxidase (cbb3-type) subunit CcoP (EC 1.9.3.1) | Complex IV: Cytochrome c oxidase (cbb3) | -0.40 | 5.25E-23 | 1.25 | 5.25E-23 | -3.92 | 5.25E-23 | -2.27 | 5.25E-23 | 1.55 | 3.31E-09 | 0.71 | 3.73E-03 | -1.58 | 1.07E-09 | -2.42 | 3.04E-26 |
| SZCCDBA067_02895 | *ccoO* | Cytochrome c oxidase (cbb3-type) subunit CcoO (EC 1.9.3.1) | Complex IV: Cytochrome c oxidase (cbb3) | -0.37 | 2.59E-42 | 1.79 | 2.59E-42 | -4.90 | 2.59E-42 | -2.74 | 2.59E-42 | 1.56 | 1.35E-09 | 0.91 | 1.29E-04 | -2.32 | 2.68E-20 | -2.97 | 1.78E-40 |
| SZCCDBA067_02896 | *ccoN* | Cytochrome c oxidase (cbb3-type) subunit CcoN (EC 1.9.3.1) | Complex IV: Cytochrome c oxidase (cbb3) | -0.34 | 2.24E-80 | 0.84 | 2.24E-80 | -4.28 | 2.24E-80 | -3.11 | 2.24E-80 | 1.94 | 8.07E-19 | 0.53 | 1.15E-02 | -1.64 | 4.35E-14 | -3.05 | 4.02E-57 |
| SZCCDBA067_02671 | *atpI* | ATP synthase protein I | ATP synthase | 0.01 | 3.19E-11 | -1.25 | 3.19E-11 | -1.07 | 3.19E-11 | -2.33 | 3.19E-11 | 1.65 | 3.51E-29 | -0.57 | 1.37E-05 | 1.13 | 1.46E-14 | -1.09 | 3.74E-18 |
| SZCCDBA067_02406 | ATPF1G, atpG | ATP synthase gamma chain (EC 3.6.3.14) | ATPase (F type) | -0.04 | 1.11E-20 | 0.73 | 1.11E-20 | -2.57 | 1.11E-20 | -1.80 | 1.11E-20 | 0.68 | 4.14E-06 | 0.24 | 8.01E-02 | -1.60 | 6.29E-30 | -2.03 | 2.16E-59 |
| SZCCDBA067_02404 | *ATPF1E, atpC* | ATP synthase epsilon chain (EC 3.6.3.14) | ATPase (F type) | 0.24 | 6.39E-08 | 0.59 | 6.39E-08 | -1.93 | 6.39E-08 | -1.58 | 6.39E-08 | 1.22 | 1.78E-12 | -0.43 | 8.82E-03 | -0.57 | 1.24E-03 | -2.21 | 3.99E-49 |
| SZCCDBA067_02408 | *ATPF1D, atpH* | ATP synthase delta chain (EC 3.6.3.14) | ATPase (F type) | -0.25 | 2.40E-10 | -1.15 | 2.40E-10 | -0.30 | 2.40E-10 | -1.20 | 2.40E-10 | 0.27 | 2.14E-01 | -1.12 | 1.72E-10 | 0.72 | 2.69E-04 | -0.68 | 1.04E-04 |
| SZCCDBA067_02405 | *ATPF1B, atpD* | ATP synthase beta chain (EC 3.6.3.14) | ATPase (F type) | 0.12 | 1.52E-23 | 1.02 | 1.52E-23 | -2.74 | 1.52E-23 | -1.84 | 1.52E-23 | 0.98 | 3.94E-09 | -0.31 | 4.59E-02 | -0.92 | 1.84E-08 | -2.21 | 8.54E-54 |
| SZCCDBA067_02407 | *ATPF1A, atpA* | ATP synthase alpha chain (EC 3.6.3.14) | ATPase (F type) | -0.09 | 1.20E-10 | -0.35 | 1.20E-10 | -1.62 | 1.20E-10 | -1.88 | 1.20E-10 | 0.18 | 3.83E-01 | -0.50 | 2.53E-03 | -1.20 | 7.37E-12 | -1.87 | 7.89E-34 |
| SZCCDBA067_02669 | *ATPF0C, atpE* | ATP synthase F0 sector subunit c (EC 3.6.3.14) | ATPase (F type) | -0.15 | 3.70E-10 | -0.15 | 3.70E-10 | -2.49 | 3.70E-10 | -2.49 | 3.70E-10 | 0.34 | 3.87E-02 | -0.66 | 2.42E-06 | -1.00 | 2.59E-11 | -2.01 | 1.17E-51 |
| SZCCDBA067_01107 | *CYC* | WD domain/cytochrome c family protein | Cytochrome c | 0.12 | 2.53E-01 | -0.33 | 2.53E-01 | 1.08 | 2.53E-01 | 0.63 | 2.53E-01 | 0.96 | 1.06E-03 | 0.53 | 1.45E-02 | 1.57 | 8.44E-09 | 1.13 | 1.07E-07 |
| SZCCDBA067_02199 | *CYC* | Cytochrome c2 | Cytochrome c | -0.40 | 6.05E-03 | -0.91 | 6.05E-03 | 0.50 | 6.05E-03 | -0.02 | 6.05E-03 | -0.29 | 4.38E-01 | -0.54 | 5.50E-02 | 0.36 | 2.88E-01 | 0.11 | 7.27E-01 |
| SZCCDBA067_02911 | *CYC* | membrane c-type cytochrome cy | Cytochrome c | 0.06 | 5.49E-02 | -0.68 | 5.49E-02 | 0.86 | 5.49E-02 | 0.12 | 5.49E-02 | -0.12 | 6.34E-01 | -1.12 | 7.52E-10 | 0.20 | 3.59E-01 | -0.80 | 8.89E-06 |
| SZCCDBA067_03541 | *CYC* | Cytochrome c2 | Cytochrome c | -0.12 | 4.89E-06 | -2.11 | 4.89E-06 | 1.38 | 4.89E-06 | -0.61 | 4.89E-06 | -0.93 | 2.63E-07 | -0.47 | 1.82E-03 | 0.96 | 1.28E-08 | 1.41 | 3.29E-20 |
| SZCCDBA067_00059 | *CYC* | Cytochrome c2 | Cytochrome c | -0.48 | 9.09E-04 | -0.30 | 9.09E-04 | -1.99 | 9.09E-04 | -1.81 | 9.09E-04 | -0.66 | 9.24E-11 | -0.40 | 1.67E-05 | -1.44 | 7.91E-48 | -1.18 | 3.66E-40 |
| SZCCDBA067_02033 | *ccmH* | Cytochrome c heme lyase subunit CcmL, apocytochrome c chaperone | Cytochrome c | -0.35 | 2.98E-04 | -0.20 | 2.98E-04 | -0.80 | 2.98E-04 | -0.65 | 2.98E-04 | -1.01 | 5.21E-09 | -0.97 | 7.11E-10 | -1.17 | 5.87E-12 | -1.12 | 2.23E-13 |
| SZCCDBA067_01314 | *ccmH* | Cytochrome c heme lyase subunit CcmL | Cytochrome c | -0.30 | 7.76E-10 | 1.47 | 7.76E-10 | -2.66 | 7.76E-10 | -0.88 | 7.76E-10 | -0.82 | 9.59E-08 | 0.14 | 3.96E-01 | -2.69 | 2.49E-69 | -1.72 | 1.00E-35 |
| SZCCDBA067_02003 | *ccmG, dsbE* | Cytochrome c-type biogenesis protein CcmG/DsbE, thiol:disulfide oxidoreductase | Cytochrome c | -0.59 | 4.81E-13 | 0.48 | 4.81E-13 | -3.85 | 4.81E-13 | -2.77 | 4.81E-13 | 0.19 | 2.99E-01 | 0.19 | 2.26E-01 | -2.42 | 1.12E-55 | -2.42 | 1.53E-69 |
| SZCCDBA067_01313 | *ccmF* | Cytochrome c heme lyase subunit CcmF | Cytochrome c | -0.37 | 1.11E-04 | 0.37 | 1.11E-04 | -1.66 | 1.11E-04 | -0.92 | 1.11E-04 | -0.85 | 6.51E-08 | -0.28 | 6.48E-02 | -1.63 | 7.85E-27 | -1.06 | 1.61E-14 |
| SZCCDBA067_01310 | *ccmE* | Cytochrome c-type biogenesis protein CcmE, heme chaperone | Cytochrome c | -0.61 | 2.27E-31 | -0.56 | 2.27E-31 | -3.88 | 2.27E-31 | -3.83 | 2.27E-31 | 0.56 | 2.52E-03 | -0.58 | 4.96E-04 | -2.04 | 2.57E-32 | -3.17 | 5.47E-95 |
| SZCCDBA067_02964 | *hcaD* | Ferredoxin reductase | Electron accepting | -0.06 | 6.31E-01 | 0.13 | 6.31E-01 | 0.09 | 6.31E-01 | 0.28 | 6.31E-01 | 0.16 | 3.23E-01 | 0.27 | 3.23E-02 | 0.21 | 1.59E-01 | 0.32 | 9.14E-03 |
| SZCCDBA067_01240 | *fpr* | Ferredoxin--NADP(+) reductase (EC 1.18.1.2) @ Flavodoxin--NADP(+) reductase (EC 1.19.1.1) | Electron accepting | -0.84 | 5.89E-20 | -1.68 | 5.89E-20 | -1.46 | 5.89E-20 | -2.31 | 5.89E-20 | -2.09 | 2.17E-32 | -2.73 | 1.39E-67 | -1.65 | 4.32E-21 | -2.30 | 2.98E-48 |
| SZCCDBA067_00457 | *ETFDH* | Electron transfer flavoprotein-ubiquinone oxidoreductase (EC 1.5.5.1) | Electron accepting | 0.21 | 1.29E-01 | 0.15 | 1.29E-01 | 0.67 | 1.29E-01 | 0.61 | 1.29E-01 | -0.60 | 8.43E-05 | -0.04 | 8.02E-01 | -0.33 | 3.24E-02 | 0.23 | 9.65E-02 |
| SZCCDBA067_02889 | *ccoS* | Type cbb3 cytochrome oxidase biogenesis protein CcoS, involved in heme b insertion | Electron accepting | -0.51 | 1.76E-10 | 1.11 | 1.76E-10 | -2.78 | 1.76E-10 | -1.16 | 1.76E-10 | 1.53 | 1.13E-08 | 0.35 | 1.87E-01 | -0.67 | 1.68E-02 | -1.85 | 9.05E-16 |
| SZCCDBA067_02891 | *ccoH* | Type cbb3 cytochrome oxidase biogenesis protein CcoH | Electron accepting | -0.40 | 8.67E-08 | 0.59 | 8.67E-08 | -2.69 | 8.67E-08 | -1.69 | 8.67E-08 | 1.62 | 2.24E-26 | 0.66 | 4.29E-06 | -1.15 | 1.15E-13 | -2.10 | 3.72E-56 |
| SZCCDBA067_02892 | *ccoG* | Type cbb3 cytochrome oxidase biogenesis protein CcoG, involved in Cu oxidation | Electron accepting | -0.49 | 1.57E-20 | 0.86 | 1.57E-20 | -3.55 | 1.57E-20 | -2.20 | 1.57E-20 | 1.88 | 9.36E-20 | 0.68 | 4.94E-04 | -1.14 | 4.74E-08 | -2.34 | 4.63E-38 |
| SZCCDBA067_02468 | *fdx* | Ferredoxin, 2Fe-2S | Electron carrier | -0.32 | 6.80E-04 | -1.17 | 6.80E-04 | 0.67 | 6.80E-04 | -0.18 | 6.80E-04 | -0.78 | 9.67E-06 | -1.13 | 6.72E-14 | 0.72 | 1.91E-05 | 0.37 | 1.86E-02 |
| SZCCDBA067_01175 | *glpA, glpD* | Aerobic glycerol-3-phosphate dehydrogenase (EC 1.1.5.3) | Electron donating | 0.01 | 1.99E-02 | 0.59 | 1.99E-02 | 0.92 | 1.99E-02 | 1.50 | 1.99E-02 | 0.36 | 1.11E-01 | 1.02 | 8.84E-10 | 0.81 | 6.21E-05 | 1.47 | 9.91E-19 |
| SZCCDBA067_03124 | *nqrF* | Na(+)-translocating NADH-quinone reductase subunit F (EC 1.6.5.8) | Na+-NQR | 0.22 | 1.44E-21 | 0.85 | 1.44E-21 | -1.93 | 1.44E-21 | -1.31 | 1.44E-21 | -1.49 | 2.04E-24 | -0.44 | 1.93E-03 | -3.60 | ####### | -2.55 | 5.42E-85 |
| SZCCDBA067_03125 | *nqrE* | Na(+)-translocating NADH-quinone reductase subunit E (EC 1.6.5.8) | Na+-NQR | 0.10 | 1.24E-03 | 1.25 | 1.24E-03 | -1.38 | 1.24E-03 | -0.23 | 1.24E-03 | -1.62 | 8.77E-18 | -0.32 | 9.86E-02 | -3.18 | 9.36E-65 | -1.87 | 1.02E-27 |
| SZCCDBA067_03126 | *nqrD* | Na(+)-translocating NADH-quinone reductase subunit D (EC 1.6.5.8) | Na+-NQR | -0.22 | 1.81E-07 | 1.86 | 1.81E-07 | -2.75 | 1.81E-07 | -0.68 | 1.81E-07 | -1.42 | 3.83E-10 | -0.25 | 2.68E-01 | -3.14 | 1.01E-45 | -1.96 | 1.12E-22 |
| SZCCDBA067_03127 | *nqrC* | Na(+)-translocating NADH-quinone reductase subunit C (EC 1.6.5.8) | Na+-NQR | -0.34 | 1.40E-12 | 1.43 | 1.40E-12 | -2.28 | 1.40E-12 | -0.51 | 1.40E-12 | -1.43 | 3.10E-17 | 0.15 | 4.07E-01 | -3.57 | ####### | -1.99 | 2.81E-40 |
| SZCCDBA067_03128 | *nqrB* | Na(+)-translocating NADH-quinone reductase subunit B (EC 1.6.5.8) | Na+-NQR | -0.29 | 3.67E-05 | 0.67 | 3.67E-05 | -1.72 | 3.67E-05 | -0.76 | 3.67E-05 | -1.45 | 6.26E-30 | 0.00 | 9.72E-01 | -3.00 | ####### | -1.55 | 3.07E-42 |
| SZCCDBA067_03129 | *nqrA* | Na(+)-translocating NADH-quinone reductase subunit A (EC 1.6.5.8) | Na+-NQR | -0.20 | 3.71E-04 | -0.18 | 3.71E-04 | -1.03 | 3.71E-04 | -1.01 | 3.71E-04 | -0.54 | 1.38E-03 | -0.58 | 1.09E-04 | -1.77 | 1.30E-29 | -1.81 | 4.63E-38 |

Table S5. Relative abundances of oxidative-type *dsrAB* in *TARA* Ocean metagenomes.

| Accession | Oxygen (μmol/kg) | Oxygen category | *dsrA_Rhodobacterales* (Reads) | *dsrA*_all (Reads) | *dsrA_Rhodobacterales* (copy per genome) | *dsrA*_all (copy per | *dsrA* ratio (*dsrA_Rhodobacterales*/all) | *dsrB_Rhodobacterales* (Reads) | *dsrB*_all (Reads) | *dsrB_Rhodobacterales* (copy per genome) | *dsrB*_all (copy per | *dsrB* ratio (*dsrB_Rhodobacterales*/all) | *soxA* (copy per genome) | *soxB* (copy per genome) | *soxC* (copy per genome) |
|---|---|---|---|---|---|---|---|---|---|---|---|---|---|---|---|
| ERR599127 | 0.6 | Nanooxic | 22 | 1243 | 0.0590 | 0.1104 | 1.7699 | 15 | 951 | 0.0486 | 0.0858 | 1.5773 | 0.0807 | 0.2460 | 0.1935 |
| ERR599152 | 0.6 | Nanooxic | 24 | 1219 | 0.0560 | 0.1064 | 1.9688 | 12 | 1026 | 0.0338 | 0.0910 | 1.1696 | 0.0764 | 0.2408 | 0.1806 |
| ERR599076 | 0.6 | Nanooxic | 20 | 1195 | 0.0510 | 0.1081 | 1.6736 | 12 | 989 | 0.0369 | 0.0909 | 1.2133 | 0.0737 | 0.2547 | 0.1934 |
| ERR599015 | 0.6 | Nanooxic | 26 | 1231 | 0.0628 | 0.1113 | 2.1121 | 14 | 989 | 0.0408 | 0.0908 | 1.4156 | 0.0790 | 0.2600 | 0.1800 |
| ERR598953 | 0.7 | Nanooxic | 27 | 1454 | 0.0659 | 0.1268 | 1.8569 | 12 | 1065 | 0.0353 | 0.0944 | 1.1268 | 0.0854 | 0.2009 | 0.1559 |
| ERR598946 | 0.7 | Nanooxic | 64 | 3710 | 0.0562 | 0.1273 | 1.7251 | 43 | 3019 | 0.0456 | 0.1053 | 1.4243 | 0.1075 | 0.2035 | 0.1724 |
| ERR598981 | 0.7 | Nanooxic | 10 | 705 | 0.0470 | 0.1243 | 1.4184 | 5 | 573 | 0.0284 | 0.1026 | 0.8726 | 0.0674 | 0.2234 | 0.1699 |
| ERR599004 | 0.8 | Nanooxic | 15 | 831 | 0.0450 | 0.0912 | 1.8051 | 9 | 679 | 0.0326 | 0.0757 | 1.3255 | 0.0805 | 0.2449 | 0.2677 |
| ERR599051 | 0.8 | Nanooxic | 27 | 876 | 0.0843 | 0.0917 | 3.0822 | 15 | 775 | 0.0565 | 0.0824 | 1.9355 | 0.0748 | 0.2407 | 0.2661 |
| ERR599175 | 0.8 | Nanooxic | 22 | 910 | 0.0667 | 0.0956 | 2.4176 | 12 | 758 | 0.0439 | 0.0809 | 1.5831 | 0.0936 | 0.2615 | 0.2560 |
| ERR599060 | 0.8 | Nanooxic | 23 | 924 | 0.0691 | 0.0973 | 2.4892 | 13 | 725 | 0.0471 | 0.0776 | 1.7931 | 0.0760 | 0.2373 | 0.2622 |
| ERR599020 | 1.3 | Suboxic | 55 | 3981 | 0.0162 | 0.0871 | 1.3816 | 30 | 3028 | 0.0107 | 0.0673 | 0.9908 | 0.1186 | 0.2372 | 0.2706 |
| ERR599037 | 2.3 | Suboxic | 58 | 2765 | 0.0779 | 0.1367 | 2.0976 | 23 | 2260 | 0.0373 | 0.1135 | 1.0177 | 0.0366 | 0.1515 | 0.1015 |
| ERR599172 | 2.3 | Suboxic | 19 | 1355 | 0.0498 | 0.1437 | 1.4022 | 12 | 1092 | 0.0380 | 0.1176 | 1.0989 | 0.0406 | 0.1547 | 0.1359 |
| ERR599167 | 2.4 | Suboxic | 43 | 2312 | 0.0571 | 0.1196 | 1.8599 | 24 | 1899 | 0.0384 | 0.0998 | 1.2638 | 0.0293 | 0.1303 | 0.0970 |
| ERR599109 | 2.4 | Suboxic | 22 | 1214 | 0.0645 | 0.1367 | 1.8122 | 10 | 979 | 0.0354 | 0.1120 | 1.0215 | 0.0504 | 0.1649 | 0.1314 |
| ERR599067 | 2.5 | Suboxic | 6 | 655 | 0.0195 | 0.0784 | 0.9160 | 2 | 538 | 0.0079 | 0.0654 | 0.3717 | 0.0510 | 0.1519 | 0.2036 |
| ERR598971 | 2.5 | Suboxic | 20 | 2870 | 0.0153 | 0.0778 | 0.6969 | 13 | 2384 | 0.0120 | 0.0657 | 0.5453 | 0.0482 | 0.1310 | 0.1591 |
| ERR599031 | 2.6 | Suboxic | 13 | 475 | 0.0729 | 0.0806 | 2.7368 | 10 | 395 | 0.0677 | 0.0681 | 2.5316 | 0.0396 | 0.1580 | 0.1777 |
| ERR599062 | 2.6 | Suboxic | 95 | 2821 | 0.0870 | 0.0820 | 3.3676 | 48 | 2134 | 0.0531 | 0.0630 | 2.2493 | 0.0298 | 0.1541 | 0.1811 |
| ERR599086 | 3.2 | Suboxic | 29 | 5148 | 0.0211 | 0.1140 | 0.5633 | 13 | 3783 | 0.0114 | 0.0851 | 0.3436 | 0.0803 | 0.1712 | 0.2647 |
| ERR599055 | 4.4 | Suboxic | 9 | 481 | 0.0510 | 0.0756 | 1.8711 | 1 | 348 | 0.0068 | 0.0555 | 0.2874 | 0.0608 | 0.1351 | 0.1994 |
| ERR599128 | 4.4 | Suboxic | 12 | 902 | 0.0302 | 0.0703 | 1.3304 | 1 | 645 | 0.0030 | 0.0511 | 0.1550 | 0.0588 | 0.1191 | 0.1928 |
| ERR599132 | 4.4 | Suboxic | 17 | 2395 | 0.0157 | 0.0734 | 0.7098 | 7 | 1782 | 0.0078 | 0.0555 | 0.3928 | 0.0599 | 0.1223 | 0.1836 |
| ERR599115 | 8.6 | Microoxic | 26 | 3538 | 0.0156 | 0.0719 | 0.7349 | 12 | 2822 | 0.0087 | 0.0582 | 0.4252 | 0.0718 | 0.1482 | 0.2316 |
| ERR599096 | 38.1 | Microoxic | 0 | 634 | 0.0000 | 0.0500 | 0.0000 | 1 | 546 | 0.0017 | 0.0438 | 0.1832 | 0.0758 | 0.1642 | 0.3389 |
| ERR599033 | 38.1 | Microoxic | 7 | 678 | 0.0102 | 0.0546 | 1.0324 | 0 | 575 | 0.0000 | 0.0471 | 0.0000 | 0.0850 | 0.1612 | 0.3193 |
| ERR598999 | 38.1 | Microoxic | 8 | 758 | 0.0115 | 0.0611 | 1.0554 | 2 | 559 | 0.0035 | 0.0458 | 0.3578 | 0.0807 | 0.1593 | 0.3350 |
| ERR599083 | 38.1 | Microoxic | 4 | 722 | 0.0055 | 0.0578 | 0.5540 | 2 | 575 | 0.0033 | 0.0468 | 0.3478 | 0.0719 | 0.1807 | 0.3148 |
| ERR599087 | 77 | Oxic | 21 | 1628 | 0.0052 | 0.0396 | 1.2899 | 11 | 1171 | 0.0033 | 0.0289 | 0.9394 | 0.0950 | 0.2563 | 0.3511 |
| ERR599112 | 88.2 | Oxic | 4 | 1094 | 0.0017 | 0.0299 | 0.3656 | 4 | 851 | 0.0020 | 0.0236 | 0.4700 | 0.0512 | 0.2081 | 0.2483 |
| ERR599099 | 95.4 | Oxic | 36 | 1476 | 0.0835 | 0.1229 | 2.4390 | 24 | 1266 | 0.0672 | 0.1071 | 1.8957 | 0.0709 | 0.2362 | 0.1693 |
| ERR599070 | 95.4 | Oxic | 31 | 1443 | 0.0593 | 0.1203 | 2.1483 | 16 | 1233 | 0.0369 | 0.1045 | 1.2976 | 0.0875 | 0.2114 | 0.1709 |
| ERR599147 | 95.4 | Oxic | 44 | 1511 | 0.0916 | 0.1258 | 2.9120 | 19 | 1283 | 0.0477 | 0.1085 | 1.4809 | 0.0774 | 0.2180 | 0.1738 |
| ERR598987 | 95.4 | Oxic | 44 | 1524 | 0.0920 | 0.1269 | 2.8871 | 11 | 1170 | 0.0278 | 0.0990 | 0.9402 | 0.0786 | 0.2164 | 0.1557 |
| ERR599168 | 103.9 | Oxic | 2 | 474 | 0.0010 | 0.0246 | 0.4219 | 0 | 237 | 0.0000 | 0.0125 | 0.0000 | 0.1476 | 0.2409 | 0.2736 |
| ERR599007 | 103.9 | Oxic | 7 | 1203 | 0.0013 | 0.0334 | 0.5819 | 1 | 633 | 0.0002 | 0.0179 | 0.1580 | 0.1835 | 0.3421 | 0.3686 |
| ERR598962 | 103.9 | Oxic | 1 | 228 | 0.0011 | 0.0358 | 0.4386 | 0 | 123 | 0.0000 | 0.0196 | 0.0000 | 0.1956 | 0.3601 | 0.3650 |
| ERR598985 | 108.2 | Oxic | 6 | 2043 | 0.0035 | 0.0529 | 0.2937 | 3 | 1543 | 0.0021 | 0.0406 | 0.1944 | 0.0706 | 0.1541 | 0.2317 |
| ERR599130 | 132.8 | Oxic | 2 | 1165 | 0.0007 | 0.0343 | 0.1717 | 0 | 496 | 0.0000 | 0.0148 | 0.0000 | 0.0821 | 0.2192 | 0.3492 |
| ERR599013 | 132.8 | Oxic | 0 | 225 | 0.0000 | 0.0350 | 0.0000 | 1 | 104 | 0.0024 | 0.0164 | 0.9615 | 0.0760 | 0.2372 | 0.3424 |
| ERR599047 | 138.1 | Oxic | 10 | 1719 | 0.0063 | 0.0498 | 0.5817 | 6 | 1441 | 0.0046 | 0.0424 | 0.4164 | 0.0637 | 0.1794 | 0.3019 |
| ERR598980 | 138.7 | Oxic | 18 | 2618 | 0.0111 | 0.0674 | 0.6875 | 11 | 2188 | 0.0082 | 0.0572 | 0.5027 | 0.0695 | 0.1615 | 0.2643 |
| ERR599048 | 143.2 | Oxic | 4 | 337 | 0.0067 | 0.0383 | 1.1869 | 6 | 220 | 0.0121 | 0.0254 | 2.7273 | 0.0915 | 0.2055 | 0.2736 |
| ERR599005 | 143.2 | Oxic | 15 | 680 | 0.0129 | 0.0388 | 2.2059 | 9 | 436 | 0.0094 | 0.0253 | 2.0642 | 0.0874 | 0.1885 | 0.2650 |
| ERR599014 | 144.6 | Oxic | 8 | 1616 | 0.0012 | 0.0321 | 0.4950 | 5 | 938 | 0.0009 | 0.0189 | 0.5330 | 0.1910 | 0.3636 | 0.3828 |
| ERR598977 | 148.3 | Oxic | 2 | 498 | 0.0027 | 0.0675 | 0.4016 | 0 | 166 | 0.0000 | 0.0229 | 0.0000 | 0.1267 | 0.2937 | 0.3621 |
| ERR599053 | 148.3 | Oxic | 9 | 2707 | 0.0025 | 0.0765 | 0.3325 | 7 | 947 | 0.0024 | 0.0272 | 0.7392 | 0.1327 | 0.2891 | 0.3972 |
| ERR598949 | 154.7 | Oxic | 1 | 270 | 0.0017 | 0.0397 | 0.3704 | 0 | 155 | 0.0000 | 0.0231 | 0.0000 | 0.1817 | 0.3393 | 0.3935 |
| ERR599082 | 154.7 | Oxic | 7 | 1532 | 0.0019 | 0.0447 | 0.4569 | 2 | 730 | 0.0007 | 0.0216 | 0.2740 | 0.1726 | 0.3454 | 0.3891 |
| ERR599071 | 155.6 | Oxic | 10 | 2098 | 0.0066 | 0.0605 | 0.4766 | 10 | 1620 | 0.0080 | 0.0474 | 0.6173 | 0.0622 | 0.1361 | 0.2830 |
| ERR599085 | 155.6 | Oxic | 3 | 497 | 0.0102 | 0.0654 | 0.6036 | 3 | 341 | 0.0123 | 0.0456 | 0.8798 | 0.0576 | 0.1541 | 0.3556 |
| ERR599149 | 161.7 | Oxic | 0 | 154 | 0.0000 | 0.0284 | 0.0000 | 1 | 141 | 0.0058 | 0.0264 | 0.7092 | 0.0876 | 0.1634 | 0.2834 |
| ERR599044 | 161.7 | Oxic | 7 | 943 | 0.0062 | 0.0317 | 0.7423 | 5 | 730 | 0.0053 | 0.0249 | 0.6849 | 0.0687 | 0.1378 | 0.2735 |
| ERR598944 | 175.3 | Oxic | 11 | 1580 | 0.0052 | 0.0368 | 0.6962 | 8 | 1265 | 0.0046 | 0.0299 | 0.6324 | 0.0537 | 0.1240 | 0.3191 |
| ERR599111 | 175.4 | Oxic | 20 | 1302 | 0.0103 | 0.0389 | 1.5361 | 23 | 1016 | 0.0143 | 0.0308 | 2.2638 | 0.0658 | 0.2554 | 0.3831 |
| ERR598975 | 175.4 | Oxic | 3 | 307 | 0.0080 | 0.0435 | 0.9772 | 5 | 210 | 0.0161 | 0.0302 | 2.3810 | 0.0709 | 0.2599 | 0.3837 |
| ERR599032 | 177 | Oxic | 2 | 778 | 0.0004 | 0.0185 | 0.2571 | 2 | 542 | 0.0005 | 0.0131 | 0.3690 | 0.1537 | 0.2919 | 0.3619 |
| ERR599038 | 179.9 | Oxic | 0 | 642 | 0.0000 | 0.0137 | 0.0000 | 1 | 440 | 0.0002 | 0.0095 | 0.2273 | 0.1375 | 0.2748 | 0.3278 |
| ERR598948 | 179.9 | Oxic | 2 | 607 | 0.0007 | 0.0149 | 0.3295 | 5 | 406 | 0.0020 | 0.0101 | 1.2315 | 0.0690 | 0.2644 | 0.3974 |
| ERR599061 | 180 | Oxic | 0 | 190 | 0.0000 | 0.0281 | 0.0000 | 3 | 117 | 0.0077 | 0.0176 | 2.5641 | 0.1266 | 0.2858 | 0.4741 |
| ERR599097 | 180 | Oxic | 7 | 1104 | 0.0024 | 0.0270 | 0.6341 | 15 | 795 | 0.0061 | 0.0198 | 1.8868 | 0.0998 | 0.2859 | 0.4342 |
| ERR599049 | 182.8 | Oxic | 1 | 621 | 0.0004 | 0.0275 | 0.1610 | 1 | 503 | 0.0005 | 0.0227 | 0.1988 | 0.1907 | 0.3467 | 0.4112 |
| ERR599134 | 182.8 | Oxic | 2 | 340 | 0.0015 | 0.0298 | 0.5882 | 1 | 250 | 0.0009 | 0.0222 | 0.4000 | 0.1827 | 0.3524 | 0.4457 |
| ERR598991 | 184.1 | Oxic | 1 | 507 | 0.0004 | 0.0249 | 0.1972 | 0 | 342 | 0.0000 | 0.0170 | 0.0000 | 0.1669 | 0.3594 | 0.4202 |
| ERR598959 | 184.1 | Oxic | 0 | 223 | 0.0000 | 0.0229 | 0.0000 | 2 | 158 | 0.0023 | 0.0165 | 1.2658 | 0.1800 | 0.3577 | 0.4299 |
| ERR599026 | 185 | Oxic | 3 | 1097 | 0.0007 | 0.0238 | 0.2735 | 1 | 771 | 0.0003 | 0.0170 | 0.1297 | 0.1369 | 0.2772 | 0.3094 |
| ERR599054 | 185.2 | Oxic | 1 | 922 | 0.0002 | 0.0205 | 0.1085 | 4 | 761 | 0.0011 | 0.0172 | 0.5256 | 0.1217 | 0.2668 | 0.3586 |
| ERR599045 | 185.2 | Oxic | 0 | 201 | 0.0000 | 0.0218 | 0.0000 | 0 | 162 | 0.0000 | 0.0179 | 0.0000 | 0.1259 | 0.2783 | 0.3631 |
| ERR598992 | 186.2 | Oxic | 0 | 426 | 0.0000 | 0.0092 | 0.0000 | 0 | 323 | 0.0000 | 0.0071 | 0.0000 | 0.0887 | 0.2045 | 0.2619 |
| ERR599019 | 187.3 | Oxic | 0 | 139 | 0.0000 | 0.0137 | 0.0000 | 0 | 124 | 0.0000 | 0.0124 | 0.0000 | 0.0986 | 0.3261 | 0.4675 |
| ERR599138 | 187.3 | Oxic | 1 | 825 | 0.0002 | 0.0154 | 0.1212 | 4 | 617 | 0.0009 | 0.0117 | 0.6483 | 0.0903 | 0.3075 | 0.4783 |
| ERR599091 | 187.3 | Oxic | 0 | 263 | 0.0000 | 0.0182 | 0.0000 | 2 | 237 | 0.0016 | 0.0166 | 0.8439 | 0.1435 | 0.2251 | 0.2672 |
| ERR599066 | 187.3 | Oxic | 1 | 305 | 0.0007 | 0.0214 | 0.3279 | 0 | 244 | 0.0000 | 0.0174 | 0.0000 | 0.1212 | 0.2216 | 0.2647 |
| ERR599119 | 187.3 | Oxic | 0 | 286 | 0.0000 | 0.0189 | 0.0000 | 1 | 227 | 0.0008 | 0.0153 | 0.4405 | 0.1312 | 0.2285 | 0.2526 |
| ERR599114 | 187.3 | Oxic | 1 | 296 | 0.0007 | 0.0207 | 0.3378 | 1 | 263 | 0.0009 | 0.0187 | 0.3802 | 0.1361 | 0.2431 | 0.2684 |
| ERR599074 | 187.4 | Oxic | 0 | 1112 | 0.0000 | 0.0267 | 0.0000 | 2 | 577 | 0.0006 | 0.0141 | 0.3466 | 0.1312 | 0.2932 | 0.3902 |
| ERR599011 | 187.4 | Oxic | 0 | 219 | 0.0000 | 0.0276 | 0.0000 | 1 | 106 | 0.0014 | 0.0136 | 0.9434 | 0.1189 | 0.2650 | 0.4422 |
| ERR599116 | 188.4 | Oxic | 1 | 1036 | 0.0005 | 0.0336 | 0.0965 | 1 | 837 | 0.0005 | 0.0276 | 0.1195 | 0.1254 | 0.2373 | 0.2726 |
| ERR599041 | 188.4 | Oxic | 0 | 394 | 0.0000 | 0.0345 | 0.0000 | 3 | 305 | 0.0047 | 0.0271 | 0.9836 | 0.1206 | 0.2265 | 0.2669 |
| ERR599155 | 188.4 | Oxic | 0 | 154 | 0.0000 | 0.0361 | 0.0000 | 0 | 116 | 0.0000 | 0.0276 | 0.0000 | 0.1169 | 0.2280 | 0.2667 |
| ERR599106 | 188.8 | Oxic | 0 | 1098 | 0.0000 | 0.0262 | 0.0000 | 7 | 845 | 0.0020 | 0.0205 | 0.8284 | 0.1637 | 0.3360 | 0.4368 |
| ERR598969 | 188.8 | Oxic | 0 | 209 | 0.0000 | 0.0255 | 0.0000 | 0 | 143 | 0.0000 | 0.0177 | 0.0000 | 0.1565 | 0.3633 | 0.4174 |
| ERR599141 | 189.3 | Oxic | 0 | 274 | 0.0000 | 0.0301 | 0.0000 | 0 | 209 | 0.0000 | 0.0233 | 0.0000 | 0.1330 | 0.2759 | 0.3285 |
| ERR599075 | 189.3 | Oxic | 2 | 1353 | 0.0005 | 0.0332 | 0.1478 | 6 | 918 | 0.0018 | 0.0229 | 0.6536 | 0.1328 | 0.2716 | 0.3487 |
| ERR599160 | 189.8 | Oxic | 1 | 527 | 0.0002 | 0.0144 | 0.1898 | 1 | 415 | 0.0003 | 0.0116 | 0.2410 | 0.1690 | 0.3231 | 0.4010 |
| ERR599058 | 190.1 | Oxic | 6 | 737 | 0.0015 | 0.0209 | 0.8141 | 1 | 497 | 0.0003 | 0.0143 | 0.2012 | 0.1455 | 0.3038 | 0.3754 |
| ERR599151 | 190.7 | Oxic | 1 | 286 | 0.0005 | 0.0182 | 0.3497 | 0 | 235 | 0.0000 | 0.0152 | 0.0000 | 0.1617 | 0.2513 | 0.3342 |
| ERR599080 | 190.7 | Oxic | 2 | 307 | 0.0012 | 0.0221 | 0.6515 | 1 | 257 | 0.0007 | 0.0188 | 0.3891 | 0.1638 | 0.2984 | 0.3524 |
| ERR599036 | 190.7 | Oxic | 1 | 306 | 0.0006 | 0.0221 | 0.3268 | 3 | 253 | 0.0022 | 0.0186 | 1.1858 | 0.1617 | 0.2892 | 0.3344 |
| ERR599069 | 190.7 | Oxic | 0 | 322 | 0.0000 | 0.0211 | 0.0000 | 1 | 242 | 0.0006 | 0.0161 | 0.4132 | 0.1405 | 0.2759 | 0.3313 |
| ERR588857 | 190.7 | Oxic | 4 | 600 | 0.0014 | 0.0239 | 0.6667 | 6 | 471 | 0.0025 | 0.0191 | 1.2739 | 0.1928 | 0.3007 | 0.3655 |
| ERR599039 | 190.8 | Oxic | 2 | 1355 | 0.0003 | 0.0266 | 0.1476 | 0 | 889 | 0.0000 | 0.0177 | 0.0000 | 0.2100 | 0.4171 | 0.4584 |
| ERR599139 | 191.7 | Oxic | 4 | 432 | 0.0007 | 0.0081 | 0.9259 | 1 | 283 | 0.0002 | 0.0054 | 0.3534 | 0.0993 | 0.2742 | 0.3692 |
| ERR599098 | 191.7 | Oxic | 0 | 80 | 0.0000 | 0.0085 | 0.0000 | 0 | 33 | 0.0000 | 0.0035 | 0.0000 | 0.0943 | 0.2713 | 0.3784 |
| ERR598958 | 191.8 | Oxic | 4 | 972 | 0.0022 | 0.0254 | 0.4115 | 10 | 822 | 0.0065 | 0.0218 | 1.2165 | 0.0521 | 0.1707 | 0.3087 |
| ERR599000 | 192.2 | Oxic | 0 | 1498 | 0.0000 | 0.0422 | 0.0000 | 10 | 1277 | 0.0094 | 0.0365 | 0.7831 | 0.0722 | 0.1789 | 0.3214 |
| ERR599154 | 192.2 | Oxic | 2 | 270 | 0.0111 | 0.0481 | 0.7407 | 1 | 216 | 0.0067 | 0.0391 | 0.4630 | 0.0808 | 0.1754 | 0.3426 |
| ERR599002 | 192.6 | Oxic | 7 | 1317 | 0.0016 | 0.0302 | 0.5315 | 2 | 530 | 0.0005 | 0.0123 | 0.3774 | 0.1178 | 0.2861 | 0.3858 |
| ERR599016 | 192.6 | Oxic | 0 | 309 | 0.0000 | 0.0304 | 0.0000 | 0 | 119 | 0.0000 | 0.0119 | 0.0000 | 0.1187 | 0.2765 | 0.4124 |
| ERR599100 | 193 | Oxic | 2 | 670 | 0.0005 | 0.0151 | 0.2985 | 2 | 470 | 0.0006 | 0.0108 | 0.4255 | 0.0962 | 0.2620 | 0.3400 |
| ERR599145 | 193 | Oxic | 1 | 1656 | 0.0002 | 0.0463 | 0.0604 | 2 | 584 | 0.0005 | 0.0166 | 0.3425 | 0.2098 | 0.4475 | 0.5278 |
| ERR599164 | 193 | Oxic | 2 | 628 | 0.0029 | 0.0383 | 0.3185 | 6 | 538 | 0.0104 | 0.0333 | 1.1152 | 0.0443 | 0.1366 | 0.2486 |
| ERR599021 | 193 | Oxic | 2 | 415 | 0.0048 | 0.0390 | 0.4819 | 1 | 332 | 0.0029 | 0.0317 | 0.3012 | 0.0508 | 0.1529 | 0.2768 |
| ERR599057 | 193.1 | Oxic | 2 | 1022 | 0.0005 | 0.0241 | 0.1957 | 2 | 800 | 0.0006 | 0.0192 | 0.2500 | 0.1204 | 0.2531 | 0.3131 |
| ERR599136 | 194.3 | Oxic | 2 | 760 | 0.0005 | 0.0162 | 0.2632 | 4 | 544 | 0.0011 | 0.0118 | 0.7353 | 0.1066 | 0.2526 | 0.3277 |
| ERR599137 | 194.4 | Oxic | 6 | 268 | 0.0051 | 0.0214 | 2.2388 | 2 | 185 | 0.0020 | 0.0150 | 1.0811 | 0.1206 | 0.2792 | 0.4023 |
| ERR599133 | 194.4 | Oxic | 14 | 729 | 0.0050 | 0.0256 | 1.9204 | 5 | 516 | 0.0022 | 0.0184 | 0.9690 | 0.1154 | 0.2937 | 0.4478 |
| ERR598989 | 195.1 | Oxic | 1 | 958 | 0.0001 | 0.0165 | 0.1044 | 0 | 696 | 0.0000 | 0.0121 | 0.0000 | 0.1444 | 0.2858 | 0.3472 |
| ERR599131 | 195.2 | Oxic | 4 | 501 | 0.0152 | 0.0718 | 0.7984 | 1 | 395 | 0.0046 | 0.0575 | 0.2532 | 0.0505 | 0.1924 | 0.3027 |
| ERR598947 | 195.2 | Oxic | 13 | 2814 | 0.0086 | 0.0759 | 0.4620 | 5 | 2148 | 0.0040 | 0.0589 | 0.2328 | 0.0543 | 0.1584 | 0.2887 |
| ERR599029 | 195.5 | Oxic | 1 | 896 | 0.0002 | 0.0195 | 0.1116 | 1 | 574 | 0.0002 | 0.0127 | 0.1742 | 0.2236 | 0.4404 | 0.5175 |
| ERR599030 | 196.9 | Oxic | 0 | 684 | 0.0000 | 0.0139 | 0.0000 | 0 | 546 | 0.0000 | 0.0113 | 0.0000 | 0.1195 | 0.2606 | 0.3438 |
| ERR599142 | 197.7 | Oxic | 0 | 976 | 0.0000 | 0.0204 | 0.0000 | 3 | 842 | 0.0008 | 0.0179 | 0.3563 | 0.0896 | 0.2537 | 0.3397 |
| ERR598960 | 198 | Oxic | 15 | 1664 | 0.0085 | 0.0415 | 0.9014 | 6 | 1259 | 0.0041 | 0.0319 | 0.4766 | 0.0438 | 0.1399 | 0.2489 |
| ERR599034 | 198 | Oxic | 2 | 311 | 0.0071 | 0.0475 | 0.6431 | 0 | 191 | 0.0000 | 0.0297 | 0.0000 | 0.0480 | 0.1804 | 0.3012 |
| ERR598997 | 198.6 | Oxic | 0 | 165 | 0.0000 | 0.0204 | 0.0000 | 1 | 133 | 0.0010 | 0.0167 | 0.7519 | 0.2260 | 0.4330 | 0.4458 |
| ERR599118 | 198.6 | Oxic | 2 | 991 | 0.0003 | 0.0206 | 0.2018 | 2 | 625 | 0.0003 | 0.0132 | 0.3200 | 0.2019 | 0.4027 | 0.4686 |
| ERR598984 | 199.1 | Oxic | 3 | 1019 | 0.0011 | 0.0365 | 0.2944 | 0 | 461 | 0.0000 | 0.0168 | 0.0000 | 0.1218 | 0.2738 | 0.3722 |
| ERR599105 | 199.1 | Oxic | 0 | 214 | 0.0000 | 0.0371 | 0.0000 | 1 | 90 | 0.0021 | 0.0158 | 1.1111 | 0.1224 | 0.2909 | 0.3886 |
| ERR599072 | 199.2 | Oxic | 4 | 1170 | 0.0022 | 0.0341 | 0.3419 | 3 | 1009 | 0.0020 | 0.0298 | 0.2973 | 0.0872 | 0.1830 | 0.3550 |
| ERR599012 | 199.9 | Oxic | 3 | 882 | 0.0008 | 0.0228 | 0.3401 | 0 | 513 | 0.0000 | 0.0135 | 0.0000 | 0.1295 | 0.2595 | 0.3272 |
| ERR599158 | 199.9 | Oxic | 0 | 475 | 0.0000 | 0.0537 | 0.0000 | 0 | 257 | 0.0000 | 0.0295 | 0.0000 | 0.1984 | 0.3791 | 0.4650 |
| ERR599102 | 199.9 | Oxic | 1 | 939 | 0.0005 | 0.0463 | 0.1065 | 1 | 555 | 0.0006 | 0.0278 | 0.1802 | 0.1925 | 0.3492 | 0.4000 |
| ERR599163 | 200.2 | Oxic | 0 | 97 | 0.0000 | 0.0024 | 0.0000 | 0 | 61 | 0.0000 | 0.0015 | 0.0000 | 0.1705 | 0.3515 | 0.4095 |
| ERR599169 | 200.2 | Oxic | 0 | 35 | 0.0000 | 0.0023 | 0.0000 | 0 | 28 | 0.0000 | 0.0019 | 0.0000 | 0.1540 | 0.3415 | 0.4156 |
| ERR599063 | 200.2 | Oxic | 0 | 22 | 0.0000 | 0.0027 | 0.0000 | 0 | 13 | 0.0000 | 0.0016 | 0.0000 | 0.1446 | 0.3433 | 0.3941 |
| ERR599120 | 200.5 | Oxic | 0 | 378 | 0.0000 | 0.0268 | 0.0000 | 2 | 286 | 0.0019 | 0.0206 | 0.6993 | 0.0944 | 0.2390 | 0.2968 |
| ERR599093 | 200.5 | Oxic | 0 | 699 | 0.0000 | 0.0270 | 0.0000 | 1 | 595 | 0.0005 | 0.0233 | 0.1681 | 0.0951 | 0.2285 | 0.3002 |
| ERR598954 | 202.2 | Oxic | 2 | 1057 | 0.0005 | 0.0201 | 0.1892 | 3 | 679 | 0.0008 | 0.0131 | 0.4418 | 0.0930 | 0.2162 | 0.2969 |
| ERR599108 | 203.1 | Oxic | 0 | 297 | 0.0000 | 0.0240 | 0.0000 | 0 | 163 | 0.0000 | 0.0134 | 0.0000 | 0.2155 | 0.4066 | 0.4798 |
| ERR599065 | 203.1 | Oxic | 1 | 748 | 0.0002 | 0.0247 | 0.1337 | 1 | 445 | 0.0002 | 0.0150 | 0.2247 | 0.2108 | 0.4217 | 0.4633 |
| ERR598952 | 203.1 | Oxic | 0 | 160 | 0.0000 | 0.0278 | 0.0000 | 0 | 79 | 0.0000 | 0.0140 | 0.0000 | 0.2505 | 0.4401 | 0.4956 |
| ERR599040 | 203.8 | Oxic | 3 | 85 | 0.0162 | 0.0293 | 3.5294 | 0 | 30 | 0.0000 | 0.0105 | 0.0000 | 0.0988 | 0.2101 | 0.3382 |
| ERR599148 | 203.8 | Oxic | 4 | 562 | 0.0031 | 0.0283 | 0.7117 | 7 | 275 | 0.0066 | 0.0141 | 2.5455 | 0.0982 | 0.2058 | 0.3205 |
| ERR599008 | 203.8 | Oxic | 8 | 794 | 0.0108 | 0.0318 | 1.0076 | 6 | 633 | 0.0098 | 0.0258 | 0.9479 | 0.0435 | 0.1484 | 0.2820 |
| ERR599125 | 203.8 | Oxic | 3 | 209 | 0.0153 | 0.0324 | 1.4354 | 0 | 167 | 0.0000 | 0.0263 | 0.0000 | 0.0357 | 0.1567 | 0.3347 |
| ERR599024 | 204 | Oxic | 0 | 1248 | 0.0000 | 0.0239 | 0.0000 | 2 | 1092 | 0.0005 | 0.0212 | 0.1832 | 0.0773 | 0.2208 | 0.3203 |
| ERR598967 | 204.1 | Oxic | 0 | 977 | 0.0000 | 0.0161 | 0.0000 | 2 | 863 | 0.0004 | 0.0144 | 0.2317 | 0.0375 | 0.1825 | 0.3073 |
| ERR599162 | 205.3 | Oxic | 2 | 890 | 0.0005 | 0.0176 | 0.2247 | 1 | 523 | 0.0003 | 0.0105 | 0.1912 | 0.1188 | 0.2462 | 0.2911 |
| ERR598943 | 206 | Oxic | 1 | 372 | 0.0006 | 0.0313 | 0.2688 | 0 | 189 | 0.0000 | 0.0161 | 0.0000 | 0.2039 | 0.3752 | 0.3958 |
| ERR598978 | 206 | Oxic | 4 | 1332 | 0.0007 | 0.0326 | 0.3003 | 0 | 676 | 0.0000 | 0.0168 | 0.0000 | 0.1766 | 0.3726 | 0.4139 |
| ERR599126 | 206.2 | Oxic | 2 | 1130 | 0.0008 | 0.0390 | 0.1770 | 3 | 776 | 0.0014 | 0.0272 | 0.3866 | 0.1424 | 0.3424 | 0.4598 |
| ERR599010 | 206.2 | Oxic | 0 | 166 | 0.0000 | 0.0487 | 0.0000 | 0 | 102 | 0.0000 | 0.0304 | 0.0000 | 0.1149 | 0.3756 | 0.4652 |
| ERR598990 | 206.4 | Oxic | 0 | 368 | 0.0000 | 0.0230 | 0.0000 | 1 | 291 | 0.0007 | 0.0185 | 0.3436 | 0.1234 | 0.3166 | 0.4447 |
| ERR599110 | 206.4 | Oxic | 7 | 811 | 0.0017 | 0.0223 | 0.8631 | 7 | 694 | 0.0020 | 0.0194 | 1.0086 | 0.1410 | 0.3225 | 0.4492 |
| ERR599018 | 206.4 | Oxic | 0 | 161 | 0.0000 | 0.0231 | 0.0000 | 0 | 141 | 0.0000 | 0.0205 | 0.0000 | 0.1679 | 0.3343 | 0.4351 |
| ERR598979 | 207 | Oxic | 1 | 287 | 0.0010 | 0.0291 | 0.3484 | 1 | 184 | 0.0012 | 0.0190 | 0.5435 | 0.1112 | 0.2851 | 0.3690 |
| ERR599146 | 207 | Oxic | 1 | 677 | 0.0004 | 0.0273 | 0.1477 | 6 | 437 | 0.0029 | 0.0179 | 1.3730 | 0.1009 | 0.2481 | 0.3666 |
| ERR598957 | 207.1 | Oxic | 3 | 716 | 0.0008 | 0.0154 | 0.4190 | 8 | 484 | 0.0027 | 0.0106 | 1.6529 | 0.0710 | 0.2224 | 0.3342 |
| ERR599025 | 207.4 | Oxic | 2 | 226 | 0.0024 | 0.0219 | 0.8850 | 0 | 192 | 0.0000 | 0.0189 | 0.0000 | 0.1064 | 0.2493 | 0.3810 |
| ERR598972 | 207.4 | Oxic | 0 | 223 | 0.0000 | 0.0212 | 0.0000 | 1 | 216 | 0.0013 | 0.0209 | 0.4630 | 0.1391 | 0.2772 | 0.4235 |
| ERR599023 | 207.4 | Oxic | 4 | 600 | 0.0019 | 0.0232 | 0.6667 | 4 | 482 | 0.0023 | 0.0189 | 0.8299 | 0.1311 | 0.2765 | 0.4135 |
| ERR315862 | 207.6 | Oxic | 2 | 486 | 0.0009 | 0.0213 | 0.4115 | 1 | 235 | 0.0005 | 0.0104 | 0.4255 | 0.0577 | 0.2155 | 0.7944 |
| ERR315863 | 207.6 | Oxic | 3 | 767 | 0.0012 | 0.0299 | 0.3911 | 7 | 467 | 0.0033 | 0.0185 | 1.4989 | 0.0876 | 0.1260 | 0.1313 |
| ERR599140 | 207.8 | Oxic | 0 | 300 | 0.0000 | 0.0426 | 0.0000 | 2 | 167 | 0.0034 | 0.0241 | 1.1976 | 0.1123 | 0.2032 | 0.2910 |
| ERR598993 | 207.8 | Oxic | 9 | 2184 | 0.0019 | 0.0408 | 0.4121 | 16 | 1301 | 0.0041 | 0.0247 | 1.2298 | 0.1094 | 0.2059 | 0.2656 |
| ERR599077 | 208.9 | Oxic | 2 | 818 | 0.0003 | 0.0157 | 0.2445 | 0 | 667 | 0.0000 | 0.0130 | 0.0000 | 0.1564 | 0.2937 | 0.3806 |
| ERR598970 | 210 | Oxic | 0 | 278 | 0.0000 | 0.0224 | 0.0000 | 2 | 222 | 0.0024 | 0.0182 | 0.9009 | 0.0990 | 0.2211 | 0.3017 |
| ERR599088 | 210 | Oxic | 1 | 231 | 0.0010 | 0.0193 | 0.4329 | 2 | 199 | 0.0023 | 0.0169 | 1.0050 | 0.1145 | 0.2496 | 0.2927 |
| ERR599150 | 210 | Oxic | 5 | 1474 | 0.0009 | 0.0227 | 0.3392 | 5 | 1055 | 0.0010 | 0.0165 | 0.4739 | 0.1114 | 0.2452 | 0.3066 |
| ERR599079 | 210.6 | Oxic | 2 | 339 | 0.0012 | 0.0154 | 0.5900 | 1 | 274 | 0.0007 | 0.0126 | 0.3650 | 0.0596 | 0.2342 | 0.3605 |
| ERR599042 | 210.6 | Oxic | 0 | 177 | 0.0000 | 0.0163 | 0.0000 | 3 | 133 | 0.0049 | 0.0124 | 2.2556 | 0.0545 | 0.2394 | 0.3666 |
| ERR598961 | 211.5 | Oxic | 9 | 1267 | 0.0018 | 0.0242 | 0.7103 | 2 | 1002 | 0.0005 | 0.0195 | 0.1996 | 0.1203 | 0.2857 | 0.3975 |
| ERR599028 | 211.6 | Oxic | 3 | 2050 | 0.0006 | 0.0533 | 0.1463 | 2 | 1222 | 0.0005 | 0.0323 | 0.1637 | 0.2162 | 0.4312 | 0.4693 |
| ERR598974 | 211.6 | Oxic | 0 | 427 | 0.0000 | 0.0519 | 0.0000 | 2 | 257 | 0.0026 | 0.0317 | 0.7782 | 0.2019 | 0.3995 | 0.4406 |
| ERR599123 | 212.6 | Oxic | 8 | 1306 | 0.0018 | 0.0278 | 0.6126 | 14 | 963 | 0.0038 | 0.0208 | 1.4538 | 0.1201 | 0.2874 | 0.4140 |
| ERR598968 | 214.4 | Oxic | 10 | 1271 | 0.0023 | 0.0249 | 0.7868 | 18 | 918 | 0.0049 | 0.0183 | 1.9608 | 0.0896 | 0.2861 | 0.4358 |
| ERR599159 | 214.8 | Oxic | 37 | 1776 | 0.0195 | 0.0511 | 2.0833 | 27 | 1413 | 0.0171 | 0.0413 | 1.9108 | 0.0658 | 0.1868 | 0.2750 |
| ERR599124 | 214.8 | Oxic | 7 | 373 | 0.0224 | 0.0586 | 1.8767 | 5 | 306 | 0.0193 | 0.0488 | 1.6340 | 0.0995 | 0.2254 | 0.3488 |
| ERR599135 | 215.7 | Oxic | 0 | 95 | 0.0000 | 0.0213 | 0.0000 | 1 | 81 | 0.0025 | 0.0184 | 1.2346 | 0.1815 | 0.3898 | 0.4746 |
| ERR599165 | 215.7 | Oxic | 2 | 236 | 0.0013 | 0.0193 | 0.8475 | 2 | 187 | 0.0016 | 0.0155 | 1.0695 | 0.1691 | 0.3773 | 0.4537 |
| ERR599113 | 216.8 | Oxic | 1 | 146 | 0.0011 | 0.0192 | 0.6849 | 1 | 151 | 0.0013 | 0.0201 | 0.6623 | 0.1379 | 0.3343 | 0.4411 |
| ERR599081 | 216.8 | Oxic | 6 | 888 | 0.0013 | 0.0211 | 0.6757 | 5 | 763 | 0.0013 | 0.0184 | 0.6553 | 0.1370 | 0.3494 | 0.4476 |
| ERR599046 | 217.1 | Oxic | 2 | 144 | 0.0037 | 0.0285 | 1.3889 | 2 | 96 | 0.0045 | 0.0193 | 2.0833 | 0.1571 | 0.3699 | 0.5290 |
| ERR599101 | 217.1 | Oxic | 10 | 1051 | 0.0024 | 0.0247 | 0.9515 | 8 | 797 | 0.0023 | 0.0190 | 1.0038 | 0.1370 | 0.3443 | 0.4958 |
| ERR599043 | 218 | Oxic | 2 | 1775 | 0.0003 | 0.0342 | 0.1127 | 4 | 1139 | 0.0007 | 0.0223 | 0.3512 | 0.2416 | 0.3881 | 0.3784 |
| ERR598951 | 218 | Oxic | 1 | 311 | 0.0009 | 0.0371 | 0.3215 | 0 | 189 | 0.0000 | 0.0229 | 0.0000 | 0.2933 | 0.4418 | 0.4318 |
| ERR318620 | 218.1 | Oxic | 2 | 1148 | 0.0007 | 0.0409 | 0.1742 | 4 | 626 | 0.0018 | 0.0227 | 0.6390 | 0.1270 | 0.2167 | 0.2658 |
| ERR318618 | 218.1 | Oxic | 5 | 1092 | 0.0018 | 0.0386 | 0.4579 | 4 | 693 | 0.0017 | 0.0249 | 0.5772 | 0.1249 | 0.2293 | 0.2715 |
| ERR318619 | 218.1 | Oxic | 1 | 63 | 0.0087 | 0.0466 | 1.5873 | 0 | 39 | 0.0000 | 0.0293 | 0.0000 | 0.1371 | 0.3122 | 0.4482 |
| ERR318621 | 218.1 | Oxic | 0 | 49 | 0.0000 | 0.0448 | 0.0000 | 0 | 33 | 0.0000 | 0.0307 | 0.0000 | 0.1953 | 0.3251 | 0.3802 |
| ERR599050 | 219.2 | Oxic | 3 | 1578 | 0.0003 | 0.0208 | 0.1901 | 0 | 1084 | 0.0000 | 0.0145 | 0.0000 | 0.1624 | 0.3398 | 0.3820 |
| ERR315858 | 220 | Oxic | 1 | 631 | 0.0006 | 0.0393 | 0.1585 | 1 | 445 | 0.0007 | 0.0282 | 0.2247 | 0.1870 | 0.2464 | 0.3051 |
| ERR315861 | 220 | Oxic | 0 | 601 | 0.0000 | 0.0383 | 0.0000 | 1 | 425 | 0.0008 | 0.0275 | 0.2353 | 0.2079 | 0.2392 | 0.3184 |
| ERR598963 | 220.2 | Oxic | 13 | 1386 | 0.0029 | 0.0296 | 0.9380 | 9 | 1082 | 0.0025 | 0.0235 | 0.8318 | 0.1267 | 0.2979 | 0.4225 |
| ERR599022 | 221.5 | Oxic | 2 | 145 | 0.0028 | 0.0257 | 1.3793 | 1 | 109 | 0.0017 | 0.0196 | 0.9174 | 0.1687 | 0.4625 | 0.5662 |
| ERR599006 | 221.5 | Oxic | 5 | 1060 | 0.0010 | 0.0276 | 0.4717 | 3 | 768 | 0.0007 | 0.0203 | 0.3906 | 0.1557 | 0.4220 | 0.5412 |
| ERR315860 | 224.3 | Oxic | 0 | 1233 | 0.0000 | 0.0779 | 0.0000 | 2 | 657 | 0.0017 | 0.0422 | 0.3044 | 0.2075 | 0.2969 | 0.3437 |
| ERR315859 | 224.3 | Oxic | 1 | 1211 | 0.0007 | 0.0758 | 0.0826 | 1 | 680 | 0.0008 | 0.0433 | 0.1471 | 0.2124 | 0.3044 | 0.3515 |
| ERR599052 | 224.4 | Oxic | 9 | 2676 | 0.0011 | 0.0388 | 0.3363 | 7 | 1914 | 0.0010 | 0.0282 | 0.3657 | 0.1795 | 0.3533 | 0.3945 |
| ERR598995 | 225.5 | Oxic | 41 | 1646 | 0.0136 | 0.0384 | 2.4909 | 23 | 1287 | 0.0092 | 0.0305 | 1.7871 | 0.0874 | 0.2919 | 0.4544 |
| ERR598942 | 227.7 | Oxic | 30 | 2022 | 0.0056 | 0.0505 | 1.4837 | 8 | 1156 | 0.0018 | 0.0293 | 0.6920 | 0.2064 | 0.3314 | 0.3703 |
| ERR599170 | 228.4 | Oxic | 8 | 2003 | 0.0016 | 0.0419 | 0.3994 | 6 | 1421 | 0.0014 | 0.0302 | 0.4222 | 0.1556 | 0.3498 | 0.4345 |
| ERR598986 | 228.5 | Oxic | 11 | 1786 | 0.0021 | 0.0400 | 0.6159 | 8 | 1342 | 0.0019 | 0.0305 | 0.5961 | 0.1439 | 0.3431 | 0.4942 |
| ERR599094 | 229.5 | Oxic | 14 | 2683 | 0.0046 | 0.0686 | 0.5218 | 6 | 1307 | 0.0024 | 0.0340 | 0.4591 | 0.1324 | 0.2956 | 0.4771 |
| ERR599153 | 229.5 | Oxic | 4 | 451 | 0.0078 | 0.0599 | 0.8869 | 2 | 246 | 0.0047 | 0.0332 | 0.8130 | 0.1367 | 0.2981 | 0.4702 |
| ERR598996 | 230.1 | Oxic | 3 | 2186 | 0.0007 | 0.0471 | 0.1372 | 3 | 1420 | 0.0008 | 0.0311 | 0.2113 | 0.1440 | 0.3179 | 0.3900 |
| ERR599166 | 230.2 | Oxic | 16 | 1681 | 0.0089 | 0.0411 | 0.9518 | 7 | 1420 | 0.0047 | 0.0353 | 0.4930 | 0.0644 | 0.1569 | 0.2973 |
| ERR599017 | 231.7 | Oxic | 9 | 336 | 0.0086 | 0.0300 | 2.6786 | 4 | 217 | 0.0046 | 0.0197 | 1.8433 | 0.0991 | 0.2593 | 0.3310 |
| ERR599103 | 231.7 | Oxic | 6 | 214 | 0.0087 | 0.0273 | 2.8037 | 2 | 136 | 0.0035 | 0.0176 | 1.4706 | 0.1006 | 0.2415 | 0.3436 |
| ERR599056 | 231.7 | Oxic | 5 | 216 | 0.0074 | 0.0274 | 2.3148 | 0 | 162 | 0.0000 | 0.0209 | 0.0000 | 0.1024 | 0.2416 | 0.3239 |
| ERR599129 | 231.9 | Oxic | 3 | 228 | 0.0040 | 0.0262 | 1.3158 | 4 | 149 | 0.0064 | 0.0174 | 2.6846 | 0.1048 | 0.2593 | 0.3468 |
| ERR599174 | 231.9 | Oxic | 5 | 374 | 0.0043 | 0.0280 | 1.3369 | 5 | 260 | 0.0052 | 0.0198 | 1.9231 | 0.1142 | 0.2881 | 0.3626 |
| ERR599171 | 231.9 | Oxic | 4 | 212 | 0.0046 | 0.0237 | 1.8868 | 4 | 145 | 0.0056 | 0.0165 | 2.7586 | 0.0998 | 0.2649 | 0.3638 |
| ERR598976 | 232.1 | Oxic | 8 | 2438 | 0.0014 | 0.0475 | 0.3281 | 6 | 1613 | 0.0013 | 0.0319 | 0.3720 | 0.1639 | 0.3531 | 0.4304 |
| ERR598983 | 233.9 | Oxic | 26 | 1090 | 0.0061 | 0.0236 | 2.3853 | 13 | 831 | 0.0037 | 0.0183 | 1.5644 | 0.0960 | 0.2107 | 0.3264 |
| ERR598965 | 237.2 | Oxic | 7 | 1327 | 0.0008 | 0.0312 | 0.5275 | 3 | 963 | 0.0004 | 0.0230 | 0.3115 | 0.3013 | 0.5253 | 0.5622 |
| ERR599073 | 237.9 | Oxic | 2 | 423 | 0.0023 | 0.0399 | 0.4728 | 3 | 240 | 0.0042 | 0.0230 | 1.2500 | 0.1036 | 0.2085 | 0.2626 |
| ERR599092 | 237.9 | Oxic | 4 | 2051 | 0.0009 | 0.0405 | 0.1950 | 5 | 1126 | 0.0013 | 0.0226 | 0.4440 | 0.1055 | 0.2001 | 0.2625 |
| ERR599068 | 238.9 | Oxic | 0 | 274 | 0.0000 | 0.0278 | 0.0000 | 1 | 189 | 0.0013 | 0.0195 | 0.5291 | 0.1187 | 0.2556 | 0.3406 |
| ERR599173 | 238.9 | Oxic | 6 | 628 | 0.0028 | 0.0299 | 0.9554 | 6 | 439 | 0.0033 | 0.0212 | 1.3667 | 0.1248 | 0.2694 | 0.3531 |
| ERR598973 | 238.9 | Oxic | 0 | 297 | 0.0000 | 0.0278 | 0.0000 | 3 | 186 | 0.0033 | 0.0177 | 1.6129 | 0.1153 | 0.2546 | 0.3247 |
| ERR599107 | 240.4 | Oxic | 3 | 316 | 0.0028 | 0.0300 | 0.9494 | 2 | 214 | 0.0023 | 0.0206 | 0.9346 | 0.1115 | 0.2753 | 0.3583 |
| ERR598982 | 240.4 | Oxic | 2 | 295 | 0.0020 | 0.0274 | 0.6780 | 6 | 254 | 0.0071 | 0.0239 | 2.3622 | 0.1195 | 0.2553 | 0.3606 |
| ERR599078 | 243.1 | Oxic | 19 | 1453 | 0.0029 | 0.0304 | 1.3076 | 15 | 1073 | 0.0028 | 0.0228 | 1.3979 | 0.1536 | 0.3632 | 0.4588 |
| ERR599064 | 244.9 | Oxic | 4 | 1223 | 0.0007 | 0.0322 | 0.3271 | 3 | 738 | 0.0007 | 0.0197 | 0.4065 | 0.2362 | 0.3549 | 0.3762 |
| ERR599144 | 249.4 | Oxic | 1 | 254 | 0.0006 | 0.0250 | 0.3937 | 1 | 228 | 0.0007 | 0.0228 | 0.4386 | 0.4635 | 0.5844 | 0.6056 |
| ERR598994 | 249.4 | Oxic | 1 | 311 | 0.0005 | 0.0333 | 0.3215 | 2 | 226 | 0.0013 | 0.0246 | 0.8850 | 0.4757 | 0.6007 | 0.6350 |
| ERR599035 | 305 | Oxic | 0 | 134 | 0.0000 | 0.0094 | 0.0000 | 0 | 115 | 0.0000 | 0.0082 | 0.0000 | 0.2871 | 0.3373 | 0.4796 |
| ERR599009 | 305 | Oxic | 1 | 209 | 0.0002 | 0.0096 | 0.4785 | 0 | 175 | 0.0000 | 0.0082 | 0.0000 | 0.2674 | 0.3505 | 0.4663 |
| ERR599027 | 306.2 | Oxic | 0 | 157 | 0.0000 | 0.0135 | 0.0000 | 0 | 117 | 0.0000 | 0.0102 | 0.0000 | 0.2592 | 0.3261 | 0.4678 |
| ERR599122 | 306.2 | Oxic | 0 | 700 | 0.0000 | 0.0148 | 0.0000 | 0 | 561 | 0.0000 | 0.0121 | 0.0000 | 0.2870 | 0.3605 | 0.4513 |
| ERR599104 | 325.4 | Oxic | 0 | 81 | 0.0000 | 0.0095 | 0.0000 | 0 | 62 | 0.0000 | 0.0074 | 0.0000 | 0.2745 | 0.3772 | 0.4960 |
| ERR599121 | 325.4 | Oxic | 2 | 362 | 0.0007 | 0.0093 | 0.5525 | 1 | 303 | 0.0004 | 0.0079 | 0.3300 | 0.2525 | 0.3477 | 0.4425 |
| ERR598945 | 338.3 | Oxic | 0 | 3 | 0.0000 | 0.0004 | 0.0000 | 0 | 4 | 0.0000 | 0.0005 | 0.0000 | 0.2125 | 0.2835 | 0.3881 |
| ERR599059 | 338.3 | Oxic | 0 | 32 | 0.0000 | 0.0008 | 0.0000 | 0 | 33 | 0.0000 | 0.0009 | 0.0000 | 0.2303 | 0.2610 | 0.3459 |
| ERR599176 | 343.4 | Oxic | 0 | 13 | 0.0000 | 0.0010 | 0.0000 | 0 | 7 | 0.0000 | 0.0006 | 0.0000 | 0.1865 | 0.2392 | 0.3711 |
| ERR599090 | 343.4 | Oxic | 0 | 16 | 0.0000 | 0.0004 | 0.0000 | 0 | 10 | 0.0000 | 0.0002 | 0.0000 | 0.1655 | 0.2325 | 0.3474 |

Table S6. Relative abundances of oxidative-type *dsrAB* in marine sediments.

| Accession | *dsrA_Rhodobacterales* (Reads) | *dsrA*_all (Reads) | *dsrA_Rhodobacterales* (copy per genome) | *dsrA*_all (copy per | *dsrA* ratio (dsrA_Rhodobacterales/all) | *dsrB*_Rhodobacterales (Reads) | *dsrB*_all (Reads) | *dsrB*_Rhodobacterales (copy per genome) | *dsrB*_all (copy per genome) | *dsrB* ratio (dsrB_Rhodobacterales/all) | *soxA*_all (copy per genome) | *soxB*_all (copy per genome) | NCBI Bioproject | Type | Depth | Geographic Location | Sample description | Latitude | Longitude |
|---|---|---|---|---|---|---|---|---|---|---|---|---|---|---|---|---|---|---|---|
| SRR7152120 | 81 | 25129 | 0.0121 | 0.3135 | 0.3223 | 907 | 17788 | 0.1640 | 0.2255 | 5.0989 | 0.5445 | 0.8439 | PRJNA465668 | Salt marsh sediment | 0 | Georgia, United States | Coastal salt marsh | 31.972 | -81.028 |
| SRR6824232 | 83 | 24287 | 0.0141 | 0.3165 | 0.3417 | 868 | 17294 | 0.1785 | 0.2290 | 5.0191 | 0.5499 | 0.8159 | PRJNA406784 | Salt marsh sediment | 0 | Georgia, United States | Coastal salt marsh | 31.972 | -81.028 |
| SRR7152353 | 82 | 19980 | 0.0106 | 0.3105 | 0.4104 | 790 | 15333 | 0.1236 | 0.2421 | 5.1523 | 0.6778 | 1.0387 | PRJNA465672 | Salt marsh sediment | 0 | Georgia, United States | Coastal salt marsh | 31.972 | -81.028 |
| SRR7151429 | 86 | 13972 | 0.0159 | 0.1988 | 0.6155 | 757 | 11189 | 0.1687 | 0.1617 | 6.7656 | 0.5301 | 0.6728 | PRJNA465665 | Salt marsh sediment | 0 | Georgia, United States | Coastal salt marsh | 31.972 | -81.028 |
| SRR7152439 | 41 | 22706 | 0.0078 | 0.3452 | 0.1806 | 734 | 14712 | 0.1676 | 0.2272 | 4.9891 | 0.5205 | 0.7963 | PRJNA465673 | Salt marsh sediment | 0 | Georgia, United States | Coastal salt marsh | 31.972 | -81.028 |
| SRR6978759 | 48 | 21946 | 0.0089 | 0.3147 | 0.2187 | 720 | 15322 | 0.1606 | 0.2232 | 4.6991 | 0.5512 | 0.8163 | PRJNA443837 | Salt marsh sediment | 0 | Georgia, United States | Coastal salt marsh | 31.972 | -81.028 |
| SRR7152452 | 167 | 17121 | 0.0351 | 0.2886 | 0.9754 | 679 | 12172 | 0.1722 | 0.2084 | 5.5784 | 0.5512 | 0.8421 | PRJNA465674 | Salt marsh sediment | 0 | Georgia, United States | Coastal salt marsh | 31.972 | -81.028 |
| SRR6977655 | 59 | 11631 | 0.0118 | 0.1716 | 0.5073 | 626 | 9431 | 0.1505 | 0.1413 | 6.6377 | 0.4587 | 0.6016 | PRJNA443834 | Salt marsh sediment | 0 | Georgia, United States | Coastal salt marsh | 31.972 | -81.028 |
| SRR7152119 | 52 | 17718 | 0.0100 | 0.2828 | 0.2935 | 606 | 12622 | 0.1402 | 0.2047 | 4.8011 | 0.5416 | 0.8312 | PRJNA465669 | Salt marsh sediment | 0 | Georgia, United States | Coastal salt marsh | 31.972 | -81.028 |
| SRR7152676 | 33 | 20181 | 0.0065 | 0.3134 | 0.1635 | 581 | 13510 | 0.1381 | 0.2131 | 4.3005 | 0.6151 | 0.9119 | PRJNA465678 | Salt marsh sediment | 0 | Georgia, United States | Coastal salt marsh | 31.972 | -81.028 |
| SRR7154838 | 45 | 20684 | 0.0076 | 0.3009 | 0.2176 | 556 | 14754 | 0.1135 | 0.2180 | 3.7685 | 0.6964 | 1.0342 | PRJNA465684 | Salt marsh sediment | 0 | Georgia, United States | Coastal salt marsh | 31.972 | -81.028 |
| SRR7152297 | 42 | 15665 | 0.0077 | 0.2567 | 0.2681 | 508 | 10618 | 0.1122 | 0.1768 | 4.7843 | 0.5434 | 0.8426 | PRJNA465671 | Salt marsh sediment | 0 | Georgia, United States | Coastal salt marsh | 31.972 | -81.028 |
| SRR7154831 | 43 | 19233 | 0.0079 | 0.2988 | 0.2236 | 501 | 11338 | 0.1104 | 0.1789 | 4.4188 | 0.5894 | 0.9201 | PRJNA465686 | Salt marsh sediment | 0 | Georgia, United States | Coastal salt marsh | 31.972 | -81.028 |
| SRR7154800 | 34 | 19881 | 0.0068 | 0.3221 | 0.1710 | 496 | 11338 | 0.1192 | 0.1866 | 4.3747 | 0.5500 | 0.8445 | PRJNA465685 | Salt marsh sediment | 0 | Georgia, United States | Coastal salt marsh | 31.972 | -81.028 |
| SRR7152869 | 53 | 17853 | 0.0089 | 0.3225 | 0.2969 | 424 | 11261 | 0.0860 | 0.2067 | 3.7652 | 0.8299 | 1.1853 | PRJNA465679 | Salt marsh sediment | 0 | Georgia, United States | Coastal salt marsh | 31.972 | -81.028 |
| SRR7154798 | 27 | 16860 | 0.0053 | 0.3004 | 0.1601 | 416 | 9680 | 0.0982 | 0.1752 | 4.2975 | 0.5707 | 0.8980 | PRJNA465687 | Salt marsh sediment | 0 | Georgia, United States | Coastal salt marsh | 31.972 | -81.028 |
| SRR7152503 | 140 | 8518 | 0.0156 | 0.1810 | 1.6436 | 396 | 6468 | 0.0531 | 0.1396 | 6.1224 | 0.8338 | 1.3779 | PRJNA465675 | Salt marsh sediment | 0 | Georgia, United States | Coastal salt marsh | 31.972 | -81.028 |
| SRR7152871 | 34 | 14103 | 0.0080 | 0.2531 | 0.2411 | 395 | 9868 | 0.1118 | 0.1799 | 4.0028 | 0.6254 | 0.9250 | PRJNA465681 | Salt marsh sediment | 0 | Georgia, United States | Coastal salt marsh | 31.972 | -81.028 |
| ERR2179510 | 519 | 8578 | 0.1563 | 0.0669 | 6.0504 | 376 | 6511 | 0.1366 | 0.0516 | 5.7748 | 0.4492 | 0.5358 | PRJEB22775 | Mangrove sediment | | Zini Mangrove Nature Reserve, China | Mgv, Enrichment | 24.33333 | 118.95 |
| SRR7151887 | 211 | 4884 | 0.0339 | 0.0860 | 4.3202 | 358 | 4098 | 0.0693 | 0.0733 | 8.7360 | 0.6161 | 0.8559 | PRJNA465666 | Salt marsh sediment | 0 | Georgia, United States | Coastal salt marsh | 31.972 | -81.028 |
| SRR7148981 | 133 | 5985 | 0.0250 | 0.0958 | 2.2222 | 354 | 4711 | 0.0803 | 0.0766 | 7.5143 | 0.4575 | 0.6219 | PRJNA465660 | Salt marsh sediment | 0 | Georgia, United States | Coastal salt marsh | 31.972 | -81.028 |
| SRR6977656 | 332 | 2208 | 0.0405 | 0.0481 | 15.0362 | 338 | 1708 | 0.0498 | 0.0378 | 19.7892 | 0.8496 | 1.1321 | PRJNA443833 | Salt marsh sediment | 0 | Georgia, United States | Coastal salt marsh | 31.972 | -81.028 |
| SRR7152675 | 24 | 12334 | 0.0050 | 0.1902 | 0.1946 | 322 | 8220 | 0.0810 | 0.1287 | 3.9173 | 0.4609 | 0.6637 | PRJNA465677 | Salt marsh sediment | 0 | Georgia, United States | Coastal salt marsh | 31.972 | -81.028 |
| ERR2179512 | 430 | 9469 | 0.1766 | 0.0895 | 4.5411 | 305 | 7429 | 0.1511 | 0.0714 | 4.1055 | 0.5462 | 0.6710 | PRJEB22775 | Mangrove sediment | | Zini Mangrove Nature Reserve, China | CK, Control | 24.33333 | 118.95 |
| SRR7153117 | 42 | 8862 | 0.0078 | 0.1470 | 0.4739 | 298 | 6894 | 0.0665 | 0.1161 | 4.3226 | 0.6372 | 0.9416 | PRJNA465683 | Salt marsh sediment | | Georgia, United States | Coastal salt marsh | 31.972 | -81.028 |
| SRR6978757 | 307 | 1501 | 0.0524 | 0.0476 | 20.4530 | 255 | 1015 | 0.0525 | 0.0327 | 25.1232 | 0.9219 | 1.3719 | PRJNA443838 | Salt marsh sediment | 0 | Georgia, United States | Coastal salt marsh | 31.972 | -81.028 |
| SRR7152251 | 39 | 9073 | 0.0082 | 0.1415 | 0.4298 | 253 | 5008 | 0.0640 | 0.0793 | 5.0519 | 0.4295 | 0.6330 | PRJNA465670 | Salt marsh sediment | 0 | Georgia, United States | Coastal salt marsh | 31.972 | -81.028 |
| SRR7149135 | 147 | 3515 | 0.0162 | 0.0556 | 4.1821 | 250 | 2706 | 0.0333 | 0.0435 | 9.2387 | 0.8649 | 1.1423 | PRJNA465663 | Salt marsh sediment | 0 | Georgia, United States | Coastal salt marsh | 31.972 | -81.028 |
| SRR7151428 | 132 | 3960 | 0.0165 | 0.0644 | 3.3333 | 247 | 2951 | 0.0374 | 0.0488 | 8.3700 | 0.7443 | 1.0506 | PRJNA465664 | Salt marsh sediment | 0 | Georgia, United States | Coastal salt marsh | 31.972 | -81.028 |
| SRR6978420 | 193 | 1472 | 0.0190 | 0.0301 | 13.1114 | 233 | 1111 | 0.0276 | 0.0231 | 20.9721 | 0.9551 | 1.2892 | PRJNA443835 | Salt marsh sediment | 0 | Georgia, United States | Coastal salt marsh | 31.972 | -81.028 |
| SRR7149136 | 176 | 1866 | 0.0263 | 0.0327 | 9.4319 | 223 | 1527 | 0.0402 | 0.0272 | 14.6038 | 0.6987 | 0.9967 | PRJNA465662 | Salt marsh sediment | 0 | Georgia, United States | Coastal salt marsh | 31.972 | -81.028 |
| SRR7152995 | 32 | 8147 | 0.0077 | 0.1266 | 0.3928 | 204 | 5456 | 0.0591 | 0.0862 | 3.7390 | 0.4322 | 0.6187 | PRJNA465682 | Salt marsh sediment | 0 | Georgia, United States | Coastal salt marsh | 31.972 | -81.028 |
| SRR7154899 | 78 | 8921 | 0.0119 | 0.1306 | 0.8743 | 190 | 6048 | 0.0350 | 0.0900 | 3.1415 | 0.7106 | 1.0781 | PRJNA465688 | Salt marsh sediment | 0 | Georgia, United States | Coastal salt marsh | 31.972 | -81.028 |
| SRR6978760 | 155 | 1781 | 0.0240 | 0.0321 | 8.7030 | 177 | 1357 | 0.0331 | 0.0249 | 13.0435 | 0.7468 | 0.9958 | PRJNA443836 | Salt marsh sediment | 0 | Georgia, United States | Coastal salt marsh | 31.972 | -81.028 |
| SRR6978756 | 111 | 4036 | 0.0246 | 0.0864 | 2.7502 | 170 | 2287 | 0.0454 | 0.0497 | 7.4333 | 0.6007 | 0.9072 | PRJNA443839 | Salt marsh sediment | 0 | Georgia, United States | Coastal salt marsh | 31.972 | -81.028 |
| SRR6978758 | 59 | 5161 | 0.0151 | 0.1044 | 1.1432 | 164 | 3166 | 0.0507 | 0.0650 | 5.1800 | 0.4823 | 0.7119 | PRJNA443840 | Salt marsh sediment | 0 | Georgia, United States | Coastal salt marsh | 31.972 | -81.028 |
| SRR7152121 | 138 | 1910 | 0.0289 | 0.0298 | 7.2251 | 162 | 1564 | 0.0410 | 0.0248 | 10.3581 | 0.4208 | 0.5581 | PRJNA465667 | Salt marsh sediment | 0 | Georgia, United States | Coastal salt marsh | 31.972 | -81.028 |
| SRR7149128 | 109 | 1831 | 0.0193 | 0.0310 | 5.9530 | 130 | 1475 | 0.0277 | 0.0253 | 8.8136 | 0.5811 | 0.7396 | PRJNA465661 | Salt marsh sediment | 0 | Georgia, United States | Coastal salt marsh | 31.972 | -81.028 |
| SRR6979551 | 133 | 3071 | 0.0247 | 0.0730 | 4.3308 | 99 | 2158 | 0.0222 | 0.0521 | 4.5876 | 0.7685 | 1.2770 | PRJNA443843 | Salt marsh sediment | 0 | Georgia, United States | Coastal salt marsh | 31.972 | -81.028 |
| SRR7245658 | 122 | 5901 | 0.1808 | 0.1803 | 2.0674 | 98 | 4972 | 0.1753 | 0.1543 | 1.9710 | 0.9520 | 1.0070 | PRJNA473166 | Mangrove sediment | | | GBE_NI_sed | 43.06847 | -70.8633 |
| SRR6979289 | 116 | 2230 | 0.0227 | 0.0598 | 5.2018 | 98 | 1693 | 0.0232 | 0.0461 | 5.7885 | 0.7940 | 1.3170 | PRJNA443842 | Salt marsh sediment | 0 | Georgia, United States | Coastal salt marsh | 31.972 | -81.028 |
| SRR7148980 | 26 | 2042 | 0.0066 | 0.0377 | 1.2733 | 97 | 1548 | 0.0297 | 0.0290 | 6.2661 | 0.3780 | 0.4604 | PRJNA465659 | Salt marsh sediment | 0 | Georgia, United States | Coastal salt marsh | 31.972 | -81.028 |
| SRR6979426 | 71 | 3630 | 0.0148 | 0.0893 | 1.9559 | 89 | 2652 | 0.0224 | 0.0663 | 3.3560 | 0.7865 | 1.2056 | PRJNA443841 | Salt marsh sediment | 0 | Georgia, United States | Coastal salt marsh | 31.972 | -81.028 |
| SRR12659818 | 88 | 1904 | 0.1098 | 0.1101 | 4.6218 | 82 | 1489 | 0.1235 | 0.0875 | 5.5071 | 0.7375 | 0.7520 | PRJNA654835 | Salt marsh sediment | 0 | Rowley, Massachusetts, United States | untreated salt marsh sediment | 42.7489 | -70.8243 |
| ERR2564009 | 71 | 1460 | 0.0632 | 0.0626 | 4.8630 | 78 | 1186 | 0.0837 | 0.0517 | 6.5767 | 0.4119 | 0.5866 | PRJEB26615 | Intertidal Sediment | 5m | the Gulfs of Kathiawar Peninsula and Arabian Sea | GOKS4 | 22.95139 | 70.40639 |
| SRR6980411 | 68 | 4381 | 0.0075 | 0.0678 | 1.5522 | 74 | 3640 | 0.0099 | 0.0572 | 2.0330 | 0.7333 | 0.9462 | PRJNA443852 | Salt marsh sediment | 0 | Georgia, United States | Coastal salt marsh | 31.972 | -81.028 |
| SRR18461109 | 75 | 757 | 0.1824 | 0.1341 | 9.9075 | 73 | 649 | 0.2143 | 0.1168 | 11.2481 | 0.9252 | 0.8417 | PRJNA819259 | Coastal surface sediments | 10 ~ 15 cm | the Shores of Kuwait | Site 10 | 29.34742 | 48.08315 |
| SRR12659808 | 106 | 1617 | 0.1152 | 0.0932 | 6.5553 | 73 | 1384 | 0.0957 | 0.0811 | 5.2746 | 0.6722 | 0.6150 | PRJNA654833 | Salt marsh sediment | | Rowley, Massachusetts, United States | untreated salt marsh sediment | 42.7489 | -70.8243 |
| SRR18461112 | 99 | 803 | 0.1876 | 0.1326 | 12.3288 | 66 | 625 | 0.1509 | 0.1049 | 10.5600 | 0.8263 | 0.7050 | PRJNA819259 | Coastal surface sediments | 13 ~ 15 cm | the Shores of Kuwait | Site 7 | 29.17297 | 48.12441 |
| SRR6979552 | 17 | 5128 | 0.0030 | 0.0793 | 0.3315 | 65 | 4117 | 0.0141 | 0.0647 | 1.5788 | 0.5822 | 0.7480 | PRJNA443844 | Salt marsh sediment | 0 | Georgia, United States | Coastal salt marsh | 31.972 | -81.028 |
| SRR11827248 | 60 | 1573 | 0.1509 | 0.1200 | 3.8144 | 63 | 1254 | 0.1912 | 0.0972 | 5.0239 | 0.7729 | 0.6626 | PRJNA620414 | Salt marsh sediment | 0 | Rowley, Massachusetts, United States | untreated salt marsh sediment | 42.7489 | -70.8243 |
| SRR11828899 | 51 | 1379 | 0.0627 | 0.1082 | 3.6983 | 63 | 1191 | 0.0935 | 0.0949 | 5.2897 | 0.7949 | 0.8311 | PRJNA620423 | Salt marsh sediment | 0 | Rowley, Massachusetts, United States | untreated salt marsh sediment | 42.7489 | -70.8243 |
| SRR18461119 | 35 | 419 | 0.0314 | 0.0640 | 8.3532 | 61 | 363 | 0.0661 | 0.0563 | 16.8044 | 0.7847 | 0.7890 | PRJNA819259 | Coastal surface sediments | 20 ~ 15 cm | the Shores of Kuwait | Site 2 | 29.39002 | 47.98736 |
| SRR11828457 | 76 | 991 | 0.1780 | 0.0818 | 7.6690 | 61 | 804 | 0.1724 | 0.0675 | 7.5871 | 0.5225 | 0.5686 | PRJNA620416 | Salt marsh sediment | 0 | Rowley, Massachusetts, United States | untreated salt marsh sediment | 42.7489 | -70.8243 |
| SRR11828906 | 63 | 1042 | 0.1381 | 0.1007 | 6.0461 | 61 | 824 | 0.1613 | 0.0809 | 7.4029 | 0.6057 | 0.7136 | PRJNA620424 | Salt marsh sediment | 0 | Rowley, Massachusetts, United States | untreated salt marsh sediment | 42.7489 | -70.8243 |
| SRR11829000 | 58 | 1330 | 0.0783 | 0.1043 | 4.3609 | 61 | 1153 | 0.0994 | 0.0919 | 5.2905 | 0.6984 | 0.8456 | PRJNA620428 | Salt marsh sediment | 0 | Rowley, Massachusetts, United States | untreated salt marsh sediment | 42.7489 | -70.8243 |
| SRR11828890 | 52 | 1491 | 0.0844 | 0.1077 | 3.4876 | 57 | 1209 | 0.1117 | 0.0887 | 4.7146 | 0.5851 | 0.6452 | PRJNA620421 | Salt marsh sediment | 0 | Rowley, Massachusetts, United States | untreated salt marsh sediment | 42.7489 | -70.8243 |
| SRR11828957 | 48 | 1090 | 0.0597 | 0.1191 | 4.4037 | 57 | 927 | 0.0855 | 0.1029 | 6.1489 | 0.8429 | 0.8855 | PRJNA620426 | Salt marsh sediment | 0 | Rowley, Massachusetts, United States | untreated salt marsh sediment | 42.7489 | -70.8243 |
| SRR7162752 | 38 | 4865 | 0.0049 | 0.0816 | 0.7811 | 56 | 3849 | 0.0087 | 0.0656 | 1.4549 | 0.7228 | 0.9072 | PRJNA465691 | Salt marsh sediment | 0 | Georgia, United States | Coastal salt marsh | 31.972 | -81.028 |
| SRR7162755 | 36 | 3661 | 0.0047 | 0.0600 | 0.9833 | 56 | 2872 | 0.0087 | 0.0478 | 1.9499 | 0.6118 | 0.7977 | PRJNA465690 | Salt marsh sediment | 0 | Georgia, United States | Coastal salt marsh | 31.972 | -81.028 |
| SRR19627612 | 76 | 1212 | 0.1009 | 0.0993 | 6.2706 | 55 | 1002 | 0.0881 | 0.0834 | 5.4890 | 0.7940 | 0.9136 | PRJNA847709 | Mangrove sediment | | | DC | 22.49444 | 114.575 |
| ERR2179511 | 50 | 960 | 0.1120 | 0.0680 | 5.2083 | 54 | 797 | 0.1460 | 0.0574 | 6.7754 | 0.4302 | 0.5272 | PRJEB22775 | Mangrove sediment | | Zini Mangrove Nature Reserve, China | Mgv-MP, Enrichment | 24.33333 | 118.95 |
| SRR18461111 | 40 | 780 | 0.1622 | 0.1523 | 5.1282 | 50 | 615 | 0.2447 | 0.1220 | 8.1301 | 0.8971 | 0.7850 | PRJNA819259 | Coastal surface sediments | 12 ~ 15 cm | the Shores of Kuwait | Site 8 | 29.35584 | 47.9426 |
| SRR7543971 | 62 | 2797 | 0.1903 | 0.1080 | 2.2167 | 49 | 2052 | 0.1815 | 0.0805 | 2.3879 | 0.6750 | 0.8096 | PRJNA407863 | Mangrove sediment | | Yunxiao Mangrove National Nature Reserve, China. | | 23.38333 | 118.95 |
| SRR14844110 | 63 | 650 | 0.0666 | 0.0462 | 9.6923 | 48 | 471 | 0.0613 | 0.0340 | 10.1911 | 0.2631 | 0.3021 | PRJNA724996 | Reef sediment | | Xisha islands, China | | 16.49877 | 111.693 |
| SRR6980406 | 51 | 1553 | 0.0122 | 0.0326 | 3.2840 | 47 | 1178 | 0.0136 | 0.0251 | 3.9898 | 0.4454 | 0.6527 | PRJNA443850 | Salt marsh sediment | 0 | Georgia, United States | Coastal salt marsh | 31.972 | -81.028 |
| SRR6979585 | 42 | 1432 | 0.0075 | 0.0222 | 2.9330 | 47 | 1123 | 0.0101 | 0.0177 | 4.1852 | 0.4318 | 0.5653 | PRJNA443846 | Salt marsh sediment | 0 | Georgia, United States | Coastal salt marsh | 31.972 | -81.028 |
| SRR11829259 | 41 | 646 | 0.0978 | 0.0451 | 6.3467 | 47 | 500 | 0.1353 | 0.0355 | 9.4000 | 0.3136 | 0.3674 | PRJNA620436 | Salt marsh sediment | 0 | Rowley, Massachusetts, United States | untreated salt marsh sediment | 42.7489 | -70.8243 |
| SRR7162754 | 31 | 4239 | 0.0041 | 0.0637 | 0.7313 | 46 | 3360 | 0.0074 | 0.0513 | 1.3690 | 0.5619 | 0.7440 | PRJNA465692 | Salt marsh sediment | 0 | Georgia, United States | Coastal salt marsh | 31.972 | -81.028 |
| SRR19627615 | 79 | 1347 | 0.2648 | 0.1161 | 5.8649 | 45 | 1114 | 0.1820 | 0.0975 | 4.0395 | 0.7931 | 0.8786 | PRJNA847709 | Mangrove sediment | | | DZG4B | 19.98364 | 110.5689 |
| SRR11828804 | 47 | 833 | 0.0938 | 0.0837 | 5.6423 | 44 | 705 | 0.1059 | 0.0720 | 6.2411 | 0.5977 | 0.6268 | PRJNA620420 | Salt marsh sediment | 0 cm | Rowley, Massachusetts, United States | untreated salt marsh sediment | 42.7489 | -70.8243 |
| SRR8556771 | 36 | 724 | 0.1023 | 0.0561 | 4.9724 | 43 | 596 | 0.1474 | 0.0469 | 7.2148 | 0.2675 | 0.3500 | PRJNA518275 | Salt marsh sediment | 40 cm | Massachusetts, United States | Salt marsh sediment | 42.759 | -70.891 |
| SRR18461114 | 50 | 603 | 0.1827 | 0.1138 | 8.2919 | 42 | 494 | 0.1852 | 0.0947 | 8.5020 | 0.7316 | 0.6423 | PRJNA819259 | Coastal surface sediments | 15 ~ 15 cm | the Shores of Kuwait | Site 5 | 29.08565 | 48.14047 |
| SRR9710542 | 28 | 554 | 0.1051 | 0.0259 | 5.0542 | 41 | 408 | 0.1856 | 0.0194 | 10.0490 | 0.0888 | 0.2025 | PRJNA539501 | Salt marsh sediment | 40 cm | Massachusetts, United States | Salt marsh sediment | 42.722 | -70.847 |
| SRR18461113 | 44 | 364 | 0.0516 | 0.0590 | 12.0879 | 39 | 256 | 0.0552 | 0.0422 | 15.2344 | 0.5808 | 0.6146 | PRJNA819259 | Coastal surface sediments | 14 ~ 15 cm | the Shores of Kuwait | Site 6 | 29.14462 | 48.13162 |
| ERR2564006 | 39 | 1032 | 0.1114 | 0.0570 | 3.7791 | 39 | 962 | 0.1344 | 0.0540 | 4.0541 | 0.2666 | 0.5400 | PRJEB26615 | Intertidal Sediment | 32 m | the Gulfs of Kathiawar Peninsula and Arabian Sea | GOKS1 | 22.6925 | 69.24944 |
| SRR6979550 | 49 | 1480 | 0.0103 | 0.0296 | 3.3108 | 38 | 1199 | 0.0097 | 0.0244 | 3.1693 | 0.4950 | 0.6755 | PRJNA443845 | Salt marsh sediment | 0 | Georgia, United States | Coastal salt marsh | 31.972 | -81.028 |
| SRR11829002 | 48 | 1270 | 0.0880 | 0.0772 | 3.7795 | 38 | 1032 | 0.0840 | 0.0638 | 3.6822 | 0.4448 | 0.4826 | PRJNA620430 | Salt marsh sediment | 0 | Rowley, Massachusetts, United States | untreated salt marsh sediment | 42.7489 | -70.8243 |
| SRR8556609 | 26 | 348 | 0.1872 | 0.0254 | 7.4713 | 38 | 293 | 0.3301 | 0.0218 | 12.9693 | 0.0743 | 0.1814 | PRJNA518265 | Salt marsh sediment | 40 cm | Massachusetts, United States | Salt marsh sediment | 42.759 | -70.891 |
| SRR10156268 | 72 | 1523 | 0.2374 | 0.1034 | 4.7275 | 37 | 1283 | 0.1472 | 0.0885 | 2.8839 | 0.3847 | 0.5163 | PRJNA573088 | Marine sediment (permanent) | 223 - 820 m | the continental shelf of Antarctica | Amundsen Sea 2 | -72.3 | -104.3 |
| SRR11492610 | 45 | 1025 | 0.0388 | 0.1408 | 4.3902 | 37 | 832 | 0.0385 | 0.1161 | 4.4471 | 0.9589 | 1.0092 | PRJNA623061 | Permeable sediment | 0 ~ 2 cm | Port Phillip Bay, Australia | | -37.8631 | 144.971 |
| SRR11828999 | 32 | 579 | 0.1462 | 0.0780 | 5.5268 | 37 | 476 | 0.2039 | 0.0651 | 7.7731 | 0.4883 | 0.5247 | PRJNA620429 | Salt marsh sediment | 0 | Rowley, Massachusetts, United States | untreated salt marsh sediment | 42.7489 | -70.8243 |
| SRR5824292 | 53 | 952 | 0.1786 | 0.0931 | 5.5672 | 36 | 733 | 0.1464 | 0.0728 | 4.9113 | 0.5788 | 0.6874 | PRJNA393787 | Mangrove sediment | | Longhai City, Fujian, China | | 24.4 | 118.95 |
| SRR6980357 | 40 | 1014 | 0.0072 | 0.0204 | 3.9448 | 36 | 784 | 0.0078 | 0.0160 | 4.5918 | 0.5285 | 0.7130 | PRJNA443849 | Salt marsh sediment | 0 | Georgia, United States | Coastal salt marsh | 31.972 | -81.028 |
| SRR11828800 | 37 | 1182 | 0.0487 | 0.0881 | 3.1303 | 34 | 955 | 0.0540 | 0.0723 | 3.5602 | 0.6493 | 0.7646 | PRJNA620419 | Salt marsh sediment | 0 | Rowley, Massachusetts, United States | untreated salt marsh sediment | 42.7489 | -70.8243 |
| SRR11829146 | 34 | 864 | 0.0846 | 0.0961 | 3.9352 | 34 | 693 | 0.1021 | 0.0783 | 4.9062 | 0.6231 | 0.7049 | PRJNA620435 | Salt marsh sediment | | Rowley, Massachusetts, United States | untreated salt marsh sediment | 42.7489 | -70.8243 |
| SRR19627616 | 38 | 1291 | 0.2085 | 0.1162 | 2.9435 | 33 | 1089 | 0.2185 | 0.0995 | 3.0303 | 0.6725 | 0.7702 | PRJNA847709 | Mangrove sediment | | | XY4B | 19.74028 | 109.2792 |
| SRR11195982 | 46 | 1318 | 0.0855 | 0.0856 | 3.4901 | 33 | 1100 | 0.0740 | 0.0726 | 3.0000 | 0.4800 | 0.5233 | PRJNA609151 | Permeable sediment | 30 cm | Middle Park Beach, Port Phillip Bay, Australia | | -37.8513 | 144.9544 |
| SRR11492609 | 37 | 2136 | 0.0708 | 0.1928 | 1.7322 | 33 | 1773 | 0.0762 | 0.1626 | 1.8613 | 1.1430 | 0.9501 | PRJNA623061 | Permeable sediment | 6 ~ 8 cm | Port Phillip Bay, Australia | | -37.8631 | 144.971 |
| SRR14844112 | 29 | 992 | 0.0476 | 0.0490 | 2.9234 | 33 | 868 | 0.0654 | 0.0436 | 3.8018 | 0.2692 | 0.2873 | PRJNA724996 | Reef sediment | | Xisha islands, China | | 17.07702 | 111.4736 |
| SRR11829098 | 61 | 931 | 0.0948 | 0.0877 | 6.5521 | 32 | 753 | 0.0600 | 0.0721 | 4.2497 | 0.5878 | 0.6483 | PRJNA620431 | Salt marsh sediment | 0 | Rowley, Massachusetts, United States | untreated salt marsh sediment | 42.7489 | -70.8243 |
| SRR9700627 | 26 | 2139 | 0.1578 | 0.1357 | 1.2155 | 32 | 1773 | 0.2343 | 0.1143 | 1.8049 | 0.5191 | 0.5825 | PRJNA539488 | Salt marsh sediment | 10 cm | Massachusetts, United States | Salt marsh sediment | 42.722 | -70.847 |
| SRR6979610 | 22 | 4308 | 0.0044 | 0.0824 | 0.5107 | 32 | 3447 | 0.0077 | 0.0669 | 0.9283 | 0.6208 | 0.8199 | PRJNA443848 | Salt marsh sediment | 0 | Georgia, United States | Coastal salt marsh | 31.972 | -81.028 |
| SRR11195980 | 27 | 1257 | 0.0165 | 0.0616 | 2.1480 | 31 | 957 | 0.0228 | 0.0477 | 3.2393 | 0.5279 | 0.6099 | PRJNA609151 | Permeable sediment | 15 cm | Middle Park Beach, Port Phillip Bay, Australia | | -37.8513 | 144.9544 |
| SRR6979553 | 28 | 1601 | 0.0077 | 0.0362 | 1.7489 | 31 | 1326 | 0.0103 | 0.0304 | 2.3379 | 0.4655 | 0.6008 | PRJNA443847 | Salt marsh sediment | | Georgia, United States | Coastal salt marsh | 31.972 | -81.028 |
| SRR19627611 | 62 | 1065 | 0.2382 | 0.0915 | 5.8216 | 30 | 861 | 0.1391 | 0.0751 | 3.4843 | 0.5686 | 0.6137 | PRJNA847709 | Mangrove sediment | | | FT | 22.52861 | 114.0028 |
| ERR257714 | 37 | 490 | 0.1299 | 0.1232 | 7.5510 | 30 | 396 | 0.1271 | 0.1011 | 7.5758 | 1.2280 | 0.9117 | PRJEB1760 | Salt marsh sediment | 1 cm | | DMS12 | 52.96528 | 0.925833 |
| SRR9709000 | 18 | 451 | 0.1468 | 0.0267 | 3.9911 | 30 | 316 | 0.2953 | 0.0190 | 9.4937 | 0.0814 | 0.2024 | PRJNA539498 | Salt marsh sediment | 90 cm | Massachusetts, United States | Salt marsh sediment | 42.722 | -70.847 |
| SRR11492613 | 39 | 1146 | 0.0431 | 0.0899 | 3.4031 | 29 | 922 | 0.0387 | 0.0734 | 3.1453 | 0.6499 | 0.6801 | PRJNA623061 | Permeable sediment | 0 ~ 2 cm | Port Phillip Bay, Australia | | -37.8562 | 144.9642 |
| SRR11828972 | 42 | 948 | 0.0972 | 0.0735 | 4.4304 | 29 | 745 | 0.0810 | 0.0587 | 3.8926 | 0.4184 | 0.5055 | PRJNA620427 | Salt marsh sediment | 0 | Rowley, Massachusetts, United States | untreated salt marsh sediment | 42.7489 | -70.8243 |
| SRR7152504 | 32 | 1613 | 0.0077 | 0.0542 | 1.9839 | 29 | 1200 | 0.0084 | 0.0409 | 2.4167 | 0.8016 | 1.3340 | PRJNA465676 | Salt marsh sediment | 0 | Georgia, United States | Coastal salt marsh | 31.972 | -81.028 |
| SRR6980407 | 21 | 955 | 0.0026 | 0.0147 | 2.1990 | 29 | 727 | 0.0044 | 0.0114 | 3.9890 | 0.4765 | 0.6455 | PRJNA443851 | Salt marsh sediment | 0 | Georgia, United States | Coastal salt marsh | 31.972 | -81.028 |
| SRR7154837 | 17 | 4025 | 0.0032 | 0.0837 | 0.4224 | 29 | 3162 | 0.0066 | 0.0668 | 0.9171 | 0.6385 | 0.7898 | PRJNA465689 | Salt marsh sediment | 0 | Georgia, United States | Coastal salt marsh | 31.972 | -81.028 |
| ERR2564008 | 76 | 1710 | 0.0568 | 0.0789 | 4.4444 | 28 | 1388 | 0.0252 | 0.0650 | 2.0173 | 0.3867 | 0.6188 | PRJEB26615 | Intertidal Sediment | 15 m | the Gulfs of Kathiawar Peninsula and Arabian Sea | GOKS3 | 22.81833 | 70.12556 |
| SRR14844116 | 22 | 852 | 0.0309 | 0.0513 | 2.5822 | 26 | 716 | 0.0441 | 0.0438 | 3.6313 | 0.3402 | 0.3733 | PRJNA724996 | Reef sediment | | Xisha islands, China | | 16.04884 | 112.4702 |
| SRR18461116 | 50 | 524 | 0.0712 | 0.0894 | 9.5420 | 24 | 407 | 0.0412 | 0.0705 | 5.8968 | 0.7970 | 0.7041 | PRJNA819259 | Coastal surface sediments | 17 ~ 15 cm | the Shores of Kuwait | Site 3 | 29.39191 | 47.99833 |
| SRR19627618 | 60 | 1114 | 0.3642 | 0.0929 | 5.3860 | 24 | 854 | 0.1758 | 0.0724 | 2.8103 | 0.6107 | 0.7340 | PRJNA847709 | Mangrove sediment | | | ZJK2B | 23.91944 | 117.4189 |
| SRR19627617 | 47 | 970 | 0.2841 | 0.0805 | 4.8454 | 24 | 770 | 0.1750 | 0.0649 | 3.1169 | 0.5222 | 0.6210 | PRJNA847709 | Mangrove sediment | | | ZJK4B | 23.91778 | 117.4211 |
| SRR14844115 | 22 | 1192 | 0.0542 | 0.0446 | 1.8456 | 24 | 895 | 0.0714 | 0.0340 | 2.6816 | 0.1661 | 0.1806 | PRJNA724996 | Reef sediment | | Xisha islands, China | | 16.50555 | 111.5853 |
| SRR11828592 | 42 | 518 | 0.1564 | 0.0607 | 8.1081 | 24 | 410 | 0.1078 | 0.0488 | 5.8537 | 0.3928 | 0.4225 | PRJNA620417 | Salt marsh sediment | 0 | Rowley, Massachusetts, United States | untreated salt marsh sediment | 42.7489 | -70.8243 |
| SRR8556705 | 18 | 132 | 0.0988 | 0.0098 | 13.6364 | 24 | 91 | 0.1589 | 0.0069 | 26.3736 | 0.0868 | 0.1261 | PRJNA518273 | Salt marsh sediment | 220 cm | Massachusetts, United States | Salt marsh sediment | 42.759 | -70.891 |
| SRR19627610 | 58 | 1663 | 0.3853 | 0.1339 | 3.4877 | 23 | 1220 | 0.1844 | 0.0998 | 1.8852 | 0.8301 | 0.9825 | PRJNA847709 | Mangrove sediment | | | FT4B | 22.52611 | 114.0036 |
| SRR11492612 | 21 | 885 | 0.0373 | 0.0902 | 2.3729 | 23 | 624 | 0.0494 | 0.0646 | 3.6859 | 0.5836 | 0.6253 | PRJNA623061 | Permeable sediment | 6 ~ 8 cm | Port Phillip Bay, Australia | | -37.8562 | 144.9642 |
| SRR11827249 | 24 | 884 | 0.0420 | 0.0761 | 2.7149 | 23 | 699 | 0.0486 | 0.0611 | 3.2904 | 0.4646 | 0.4858 | PRJNA620415 | Salt marsh sediment | 0 | Rowley, Massachusetts, United States | untreated salt marsh sediment | 42.7489 | -70.8243 |
| SRR2134631 | 33 | 432 | 0.4458 | 0.0936 | 7.6389 | 22 | 405 | 0.3586 | 0.0892 | 5.4321 | 0.5083 | 0.6393 | PRJNA283251 | Intertidal Sediment | 0 ~ 10 cm | Hong Kong | | 22.49883 | 113.9878 |
| SRR19627608 | 42 | 893 | 0.2122 | 0.0820 | 4.7032 | 22 | 758 | 0.1341 | 0.0707 | 2.9024 | 0.7190 | 0.8654 | PRJNA847709 | Mangrove sediment | | | SK2B | 21.55417 | 109.6792 |
| SRR3620787 | 36 | 561 | 0.1580 | 0.0583 | 6.4171 | 22 | 445 | 0.1165 | 0.0470 | 4.9438 | 0.2717 | 0.2844 | PRJNA321092 | Mangrove sediment | | Rabigh coast in Saudi Arabia | MNK (mangrove mud from Al Kharrar lagoon) | 22.91396 | 38.91101 |
| SRR11195983 | 30 | 1290 | 0.0474 | 0.0922 | 2.3256 | 22 | 932 | 0.0419 | 0.0676 | 2.3605 | 0.5487 | 0.6214 | PRJNA609151 | Permeable sediment | 15 cm | Middle Park Beach, Port Phillip Bay, Australia | | -37.8513 | 144.9544 |
| SRR9700626 | 13 | 1060 | 0.1299 | 0.0682 | 1.2264 | 22 | 990 | 0.2654 | 0.0647 | 2.2222 | 0.2237 | 0.3188 | PRJNA539483 | Salt marsh sediment | 20 cm | Massachusetts, United States | Salt marsh sediment | 42.722 | -70.847 |
| SRR19627609 | 26 | 693 | 0.0634 | 0.0580 | 3.7518 | 21 | 579 | 0.0618 | 0.0492 | 3.6269 | 0.5631 | 0.5897 | PRJNA847709 | Mangrove sediment | | | JHWB | 21.42139 | 109.2053 |
| SRR8557363 | 17 | 575 | 0.1343 | 0.0459 | 2.9565 | 21 | 397 | 0.2003 | 0.0322 | 5.2897 | 0.0955 | 0.1565 | PRJNA518281 | Salt marsh sediment | 30 cm | Massachusetts, United States | Salt marsh sediment | 42.759 | -70.891 |
| SRR9710701 | 8 | 2175 | 0.0612 | 0.1737 | 0.3678 | 21 | 1785 | 0.1939 | 0.1448 | 1.1765 | 0.6911 | 0.7179 | PRJNA539504 | Salt marsh sediment | 10 cm | Massachusetts, United States | Salt marsh sediment | 42.722 | -70.847 |
| ERR257715 | 19 | 279 | 0.1699 | 0.1085 | 6.8100 | 20 | 323 | 0.2157 | 0.1276 | 6.1920 | 1.1610 | 1.0165 | PRJEB1760 | Intertidal Sediment | 1 cm | Norwich, United Kingdom | Coastal intertidal sediment | 52.96528 | 0.925833 |
| SRR8651100 | 30 | 534 | 0.2633 | 0.0968 | 5.6180 | 20 | 446 | 0.2118 | 0.0822 | 4.4843 | 0.6815 | 0.7646 | PRJNA505165 | Mangrove sediment | | Haikou, China | HKC | 19.31667 | 111.8333 |
| SRR11492608 | 22 | 1371 | 0.0837 | 0.1184 | 1.6047 | 20 | 1147 | 0.0918 | 0.1006 | 1.7437 | 0.7537 | 0.5837 | PRJNA623061 | Permeable sediment | 14 ~ 16 cm | Port Phillip Bay, Australia | | -37.8631 | 144.971 |
| SRR11829104 | 28 | 1014 | 0.0613 | 0.0773 | 2.7613 | 20 | 812 | 0.0528 | 0.0629 | 2.4631 | 0.4337 | 0.5847 | PRJNA620433 | Salt marsh sediment | 0 | Rowley, Massachusetts, United States | untreated salt marsh sediment | 42.7489 | -70.8243 |
| SRR11829102 | 18 | 1016 | 0.0553 | 0.0938 | 1.7717 | 20 | 838 | 0.0742 | 0.0786 | 2.3866 | 0.5478 | 0.5595 | PRJNA620432 | Salt marsh sediment | 0 | Rowley, Massachusetts, United States | untreated salt marsh sediment | 42.7489 | -70.8243 |
| SRR6194934 | 20 | 815 | 0.1013 | 0.0992 | 2.4540 | 20 | 623 | 0.1222 | 0.0770 | 3.2103 | 0.5462 | 0.7636 | PRJNA366577 | Salt pond sediment | 11 cm | San Francisco, United States | Eden Landing Ponds | 37.56902 | -122.102 |
| SRR11195981 | 32 | 1314 | 0.0172 | 0.0591 | 2.4353 | 19 | 1096 | 0.0123 | 0.0501 | 1.7336 | 0.5146 | 0.6265 | PRJNA609151 | Permeable sediment | 0 | Middle Park Beach, Port Phillip Bay, Australia | | -37.8513 | 144.9544 |
| SRR11829145 | 29 | 1445 | 0.0436 | 0.0963 | 2.0069 | 19 | 1129 | 0.0345 | 0.0764 | 1.6829 | 0.4695 | 0.5342 | PRJNA620434 | Salt marsh sediment | 0 | Rowley, Massachusetts, United States | untreated salt marsh sediment | 42.7489 | -70.8243 |
| SRR8556696 | 9 | 97 | 0.0541 | 0.0078 | 9.2784 | 19 | 99 | 0.1377 | 0.0081 | 19.1919 | 0.0704 | 0.1305 | PRJNA518268 | Salt marsh sediment | 230 cm | Massachusetts, United States | Salt marsh sediment | 42.759 | -70.891 |
| SRR11829268 | 9 | 1188 | 0.0381 | 0.0937 | 0.7576 | 19 | 985 | 0.0970 | 0.0789 | 1.9289 | 0.4693 | 0.5330 | PRJNA620437 | Salt marsh sediment | | Rowley, Massachusetts, United States | untreated salt marsh sediment | 42.7489 | -70.8243 |
| SRR3620786 | 30 | 1036 | 0.0236 | 0.0408 | 2.8958 | 18 | 719 | 0.0171 | 0.0288 | 2.5035 | 0.3148 | 0.3742 | PRJNA321092 | Microbial mat | | Rabigh coast in Saudi Arabia | MMK (microbial mat from Al Kharrar lagoon) | 22.91396 | 38.91101 |
| SRR12659820 | 21 | 1664 | 0.0323 | 0.0972 | 1.2620 | 18 | 1272 | 0.0334 | 0.0755 | 1.4151 | 0.5116 | 0.5609 | PRJNA654834 | Salt marsh sediment | 0 | Rowley, Massachusetts, United States | untreated salt marsh sediment | 42.7489 | -70.8243 |
| SRR18461110 | 38 | 381 | 0.1047 | 0.0626 | 9.9738 | 17 | 313 | 0.0565 | 0.0522 | 5.4313 | 0.4796 | 0.4904 | PRJNA819259 | Coastal surface sediments | 11 ~ 15 cm | the Shores of Kuwait | Site 9 | 29.35687 | 47.90808 |
| SRR14844113 | 38 | 827 | 0.0699 | 0.0494 | 4.5949 | 17 | 673 | 0.0377 | 0.0409 | 2.5260 | 0.2307 | 0.3081 | PRJNA724996 | Reef sediment | | Xisha islands, China | | 17.09958 | 111.4644 |
| SRR11828907 | 16 | 1147 | 0.0268 | 0.0861 | 1.3949 | 17 | 987 | 0.0343 | 0.0753 | 1.7224 | 0.6184 | 0.5652 | PRJNA620425 | Salt marsh sediment | 0 | Rowley, Massachusetts, United States | untreated salt marsh sediment | 42.7489 | -70.8243 |
| SRR19627606 | 42 | 1146 | 0.2374 | 0.1004 | 3.6649 | 16 | 934 | 0.1091 | 0.0831 | 1.7131 | 0.5806 | 0.6480 | PRJNA847709 | Mangrove sediment | | | BLHB | 21.54389 | 108.0525 |
| SRR19627614 | 24 | 680 | 0.0874 | 0.0566 | 3.5294 | 16 | 644 | 0.0703 | 0.0545 | 2.4845 | 0.4144 | 0.4628 | PRJNA847709 | Mangrove sediment | | | QMG3B | 18.22389 | 109.6186 |
| SRR11195984 | 21 | 721 | 0.0269 | 0.0590 | 2.9126 | 16 | 528 | 0.0247 | 0.0439 | 3.0303 | 0.4876 | 0.5553 | PRJNA609151 | Permeable sediment | 0 | Middle Park Beach, Port Phillip Bay, Australia | | -37.8513 | 144.9544 |
| ERR1341885 | 11 | 225 | 0.1763 | 0.1230 | 4.8889 | 16 | 180 | 0.3095 | 0.1000 | 8.8889 | 0.4897 | 0.5891 | PRJEB13222 | Reef sediment | | Atlantic Ocean off the coast of Faro, Algarve, Portugal | sediment metagenome | 36.97978 | -7.98911 |
| SRR8557365 | 13 | 281 | 0.1269 | 0.0239 | 4.6263 | 16 | 195 | 0.1884 | 0.0169 | 8.2051 | 0.0663 | 0.1680 | PRJNA518283 | Salt marsh sediment | 40 cm | Massachusetts, United States | Salt marsh sediment | 42.722 | -70.891 |
| SRR19627613 | 41 | 865 | 0.1186 | 0.0778 | 4.7399 | 15 | 700 | 0.0524 | 0.0639 | 2.1429 | 0.7268 | 0.7797 | PRJNA847709 | Mangrove sediment | | | BG | 22.65194 | 114.5561 |
| SRR7244169 | 9 | 1371 | 0.0569 | 0.1012 | 0.6565 | 15 | 974 | 0.1144 | 0.0730 | 1.5400 | 0.3555 | 0.4895 | PRJNA473166 | Mangrove sediment | | | Sed_Coco | 43.19744 | -70.8676 |
| SRR11492614 | 22 | 1330 | 0.0329 | 0.0701 | 1.6541 | 15 | 1131 | 0.0271 | 0.0605 | 1.3263 | 0.4861 | 0.4875 | PRJNA623061 | Permeable sediment | 14 ~ 16 cm | Port Phillip Bay, Australia | | -37.8513 | 144.9543 |
| SRR8467675 | 17 | 352 | 0.0369 | 0.0636 | 4.8295 | 15 | 271 | 0.0392 | 0.0498 | 5.5351 | 0.6242 | 0.6609 | PRJNA515295 | Permeable sediment | 0 ~ 10 cm | Melbourne, Middle Park Beach, Australia | | -37.8524 | 144.9575 |
| SRR14844106 | 19 | 895 | 0.0677 | 0.0548 | 2.1229 | 15 | 653 | 0.0645 | 0.0406 | 2.2971 | 0.1677 | 0.2090 | PRJNA724996 | Reef sediment | | Xisha islands, China | | 16.23362 | 111.8157 |
| SRR14844107 | 9 | 1179 | 0.0349 | 0.0571 | 0.7634 | 15 | 989 | 0.0701 | 0.0487 | 1.5167 | 0.1319 | 0.1803 | PRJNA724996 | Reef sediment | | Xisha islands, China | | 16.07455 | 111.7974 |
| SRR11828889 | 31 | 858 | 0.0538 | 0.0754 | 3.6131 | 15 | 722 | 0.0314 | 0.0644 | 2.0776 | 0.4514 | 0.5579 | PRJNA620422 | Salt marsh sediment | 0 | Rowley, Massachusetts, United States | untreated salt marsh sediment | 42.7489 | -70.8243 |
| SRR11829269 | 25 | 1127 | 0.0374 | 0.0792 | 2.2183 | 15 | 829 | 0.0271 | 0.0591 | 1.8094 | 0.5093 | 0.6328 | PRJNA620476 | Salt marsh sediment | 0 | Rowley, Massachusetts, United States | untreated salt marsh sediment | 42.7489 | -70.8243 |
| SRR18461115 | 7 | 217 | 0.0153 | 0.0344 | 3.2258 | 14 | 185 | 0.0368 | 0.0298 | 7.5676 | 0.3117 | 0.3778 | PRJNA819259 | Coastal surface sediments | 16 ~ 15 cm | the Shores of Kuwait | Site 4 | 29.34656 | 48.08096 |
| SRR14844114 | 22 | 1224 | 0.1786 | 0.0475 | 1.7974 | 14 | 1058 | 0.1372 | 0.0417 | 1.3233 | 0.1665 | 0.2165 | PRJNA724996 | Reef sediment | | Xisha islands, China | | 16.5251 | 111.768 |
| SRR8651098 | 27 | 270 | 0.1126 | 0.0497 | 10.0000 | 13 | 204 | 0.0654 | 0.0382 | 6.3725 | 0.3743 | 0.4310 | PRJNA505165 | Mangrove sediment | | Sanya, China | SYC | 18.25403 | 110.8167 |
| SRR8651101 | 22 | 319 | 0.2082 | 0.0570 | 6.8966 | 13 | 325 | 0.1484 | 0.0590 | 4.0000 | 0.6566 | 0.7188 | PRJNA505165 | Mangrove sediment | | Haikou, China | HKNC | 19.31667 | 111.8333 |
| SRR19627607 | 13 | 560 | 0.1859 | 0.0497 | 2.3214 | 13 | 439 | 0.2243 | 0.0396 | 2.9613 | 0.2697 | 0.3558 | PRJNA847709 | Mangrove sediment | | | KXLB | 21.86861 | 108.4883 |
| ERR1341884 | 14 | 184 | 0.1030 | 0.1011 | 7.6087 | 13 | 146 | 0.1154 | 0.0815 | 8.9041 | 0.5622 | 0.6813 | PRJEB13222 | Reef sediment | | Atlantic Ocean off the coast of Faro, Algarve, Portugal | sediment metagenome | 36.97978 | -7.98911 |
| SRR9710038 | 11 | 555 | 0.0852 | 0.0241 | 1.9820 | 13 | 445 | 0.1215 | 0.0196 | 2.9213 | 0.1009 | 0.1868 | PRJNA539499 | Salt marsh sediment | 90 cm | Massachusetts, United States | Salt marsh sediment | 42.722 | -70.847 |
| SRR11492615 | 18 | 846 | 0.0412 | 0.0763 | 2.1277 | 12 | 705 | 0.0332 | 0.0646 | 1.7021 | 0.5570 | 0.5629 | PRJNA623061 | Permeable sediment | 6 ~ 8 cm | Port Phillip Bay, Australia | | -37.8513 | 144.9543 |
| SRR14844104 | 26 | 960 | 0.0701 | 0.0615 | 2.7083 | 12 | 772 | 0.0391 | 0.0502 | 1.5544 | 0.2977 | 0.3525 | PRJNA724996 | Reef sediment | | Xisha islands, China | | 16.04229 | 112.5107 |
| SRR14844105 | 25 | 858 | 0.0721 | 0.0520 | 2.9138 | 12 | 758 | 0.0418 | 0.0467 | 1.5831 | 0.2032 | 0.2360 | PRJNA724996 | Reef sediment | | Xisha islands, China | | 16.07205 | 112.5171 |
| SRR8556611 | 12 | 91 | 0.0994 | 0.0066 | 13.1868 | 12 | 93 | 0.1199 | 0.0069 | 12.9032 | 0.0407 | 0.0783 | PRJNA518267 | Salt marsh sediment | 240 cm | Massachusetts, United States | Salt marsh sediment | 42.759 | -70.891 |
| SRR11828593 | 5 | 696 | 0.0202 | 0.0765 | 0.7184 | 12 | 661 | 0.0584 | 0.0738 | 1.8154 | 0.4177 | 0.4988 | PRJNA620418 | Salt marsh sediment | 0 | Rowley, Massachusetts, United States | untreated salt marsh sediment | 42.7489 | -70.8243 |
| SRR8651099 | 18 | 277 | 0.0588 | 0.0522 | 6.4982 | 10 | 215 | 0.0394 | 0.0411 | 4.6512 | 0.5010 | 0.5536 | PRJNA505165 | Mangrove sediment | | Sanya, China | SYNC | 18.25403 | 110.8167 |
| SRR11492616 | 15 | 843 | 0.0260 | 0.0720 | 1.7794 | 10 | 691 | 0.0209 | 0.0600 | 1.4472 | 0.4828 | 0.5406 | PRJNA623061 | Permeable sediment | 0 ~ 2 cm | Port Phillip Bay, Australia | | -37.8513 | 144.9543 |
| SRR14844118 | 19 | 806 | 0.0460 | 0.0486 | 2.3573 | 10 | 646 | 0.0292 | 0.0396 | 1.5480 | 0.2340 | 0.2670 | PRJNA724996 | Reef sediment | | Xisha islands, China | | 15.80074 | 111.2123 |
| SRR14844117 | 17 | 987 | 0.0403 | 0.0664 | 1.7224 | 10 | 731 | 0.0286 | 0.0499 | 1.3680 | 0.2589 | 0.2737 | PRJNA724996 | Reef sediment | | Xisha islands, China | | 16.06074 | 112.4689 |
| SRR6394728 | 10 | 226 | 0.0547 | 0.0520 | 4.4248 | 10 | 187 | 0.0660 | 0.0437 | 5.3476 | 0.3618 | 0.4669 | PRJNA4229171 | Salt marsh sediment | | Sydney, Australia | regenerated saltmarsh | -33.8402 | 151.0588 |
| SRR18461118 | 9 | 230 | 0.0465 | 0.0423 | 3.9130 | 9 | 179 | 0.0561 | 0.0335 | 5.0279 | 0.4085 | 0.3280 | PRJNA819259 | Coastal surface sediments | 19 ~ 15 cm | the Shores of Kuwait | Site 11 | 28.64354 | 48.36044 |
| SRR8556601 | 6 | 421 | 0.0927 | 0.0344 | 1.4252 | 9 | 388 | 0.1678 | 0.0322 | 2.3196 | 0.1573 | 0.1820 | PRJNA518263 | Salt marsh sediment | 20 cm | Massachusetts, United States | Salt marsh sediment | 42.759 | -70.891 |
| SRR6207736 | 12 | 2664 | 0.1067 | 0.0695 | 0.4505 | 8 | 2275 | 0.0858 | 0.0603 | 0.3516 | 0.4902 | 0.5816 | PRJNA366768 | Estuary sediment | | White Oak River estuary, North Carolina, United States | | 34.6402 | -77.1094 |
| SRR5210214 | 11 | 270 | 0.0186 | 0.0356 | 4.0741 | 8 | 215 | 0.0163 | 0.0288 | 3.7209 | 0.4205 | 0.4856 | PRJNA365351 | Salt marsh sediment | 0 | South San Francisco, United States | Tidal salt marsh | 37.4962 | -122.133 |
| SRR8556704 | 4 | 52 | 0.1241 | 0.0034 | 7.6923 | 8 | 55 | 0.2995 | 0.0037 | 14.5455 | 0.0151 | 0.1026 | PRJNA518271 | Salt marsh sediment | 150 cm | Massachusetts, United States | Salt marsh sediment | 42.759 | -70.891 |
| SRR11195979 | 16 | 638 | 0.1149 | 0.0471 | 2.5078 | 7 | 507 | 0.0607 | 0.0380 | 1.3807 | 0.2636 | 0.2869 | PRJNA609151 | Permeable sediment | 30 cm | Middle Park Beach, Port Phillip Bay, Australia | | -37.8513 | 144.9544 |
| SRR11492611 | 7 | 653 | 0.0474 | 0.0862 | 1.0720 | 7 | 492 | 0.0571 | 0.0660 | 1.4228 | 0.5167 | 0.4787 | PRJNA623061 | Permeable sediment | 14 ~ 16 cm | Port Phillip Bay, Australia | | -37.8562 | 144.9642 |
| SRR14844119 | 11 | 841 | 0.0264 | 0.0524 | 1.3080 | 7 | 776 | 0.0203 | 0.0491 | 0.9021 | 0.2061 | 0.2213 | PRJNA724996 | Reef sediment | | Xisha islands, China | | 16.2704 | 111.7241 |
| ERR1811638 | 3 | 372 | 0.0344 | 0.1262 | 0.8065 | 7 | 325 | 0.0970 | 0.1120 | 2.1538 | 0.8897 | 0.7295 | PRJEB19235 | Salt marsh sediment | | Abbotts Hall, United Kingdom | Natural salt marsh sediment | 51.7865 | 0.86085 |
| SRR9707467 | 1 | 477 | 0.0080 | 0.0201 | 0.2096 | 7 | 359 | 0.0676 | 0.0154 | 1.9499 | 0.1044 | 0.2128 | PRJNA539495 | Salt marsh sediment | 240 cm | Massachusetts, United States | Salt marsh sediment | 42.722 | -70.847 |
| SRR8467674 | 21 | 438 | 0.0538 | 0.0777 | 4.7945 | 7 | 350 | 0.0216 | 0.0631 | 2.0000 | 0.5555 | 0.6888 | PRJNA515295 | Subtidal sediment | 15 ~ 18 cm | Middle Park Beach, Melbourne, Australia | | -37.8524 | 144.9575 |
| SRR16005300 | 5 | 1896 | 0.0353 | 0.0994 | 0.2637 | 6 | 1435 | 0.0512 | 0.0764 | 0.4181 | 0.5257 | 0.6753 | PRJNA765124 | Estuary sediment | 0 ~ 1 cm | Yangtze Estuary, China | Sum_13C_meta | 31 | 122 |
| SRR2134641 | 7 | 167 | 0.3613 | 0.0809 | 4.1916 | 6 | 176 | 0.3737 | 0.0867 | 3.4091 | 0.4147 | 0.4753 | PRJNA283251 | Intertidal Sediment | 0 ~ 10 cm | Hong Kong | | 22.15326 | 114.39 |
| ERR1811643 | 10 | 277 | 0.1889 | 0.0790 | 3.6101 | 6 | 258 | 0.1367 | 0.0747 | 2.3256 | 0.5309 | 0.5815 | PRJEB19235 | Salt marsh sediment | | | W AHNM 2B | 51.7865 | 0.86085 |
| ERR1811653 | 7 | 250 | 0.1517 | 0.0877 | 2.8000 | 6 | 187 | 0.1569 | 0.0666 | 3.2086 | 0.5795 | 0.7027 | PRJEB19235 | Salt marsh sediment | | Fingringhoe Range, United Kingdom | Natural salt marsh sediment | 51.83182 | 0.9687 |
| ERR1811656 | 6 | 104 | 0.0798 | 0.0285 | 5.7692 | 6 | 83 | 0.0963 | 0.0231 | 7.2289 | 0.1998 | 0.2953 | PRJEB19235 | Salt marsh sediment | | Fingringhoe Range, United Kingdom | Realigned salt marsh sediment | 51.83015 | 0.9503 |
| SRR8556600 | 1 | 484 | 0.0155 | 0.0432 | 0.2066 | 6 | 400 | 0.1122 | 0.0363 | 1.5000 | 0.2142 | 0.2229 | PRJNA518262 | Salt marsh sediment | 10 cm | Massachusetts, United States | Salt marsh sediment | 42.759 | -70.891 |
| SRR8557214 | 1 | 988 | 0.0109 | 0.1078 | 0.1012 | 6 | 853 | 0.0790 | 0.0946 | 0.7034 | 0.4398 | 0.5813 | PRJNA518279 | Salt marsh sediment | 10 cm | Massachusetts, United States | Salt marsh sediment | 42.759 | -70.891 |
| ERR2564007 | 6 | 433 | 0.0349 | 0.0327 | 1.3857 | 5 | 409 | 0.0351 | 0.0313 | 1.2225 | 0.1447 | 0.3323 | PRJEB26615 | Intertidal Sediment | 28 m | the Gulfs of Kathiawar Peninsula and Arabian Sea | GOKS2 | 22.62778 | 69.70806 |
| SRR10156264 | 3 | 147 | 0.0424 | 0.0568 | 2.0408 | 5 | 136 | 0.0852 | 0.0534 | 3.6765 | 0.2968 | 0.4640 | PRJNA573088 | Marine sediment (permanent) | 223 - 820 m | the continental shelf of Antarctica | Ross Sea 1 | -75.2 | -176.59 |
| SRR11195985 | 3 | 367 | 0.0691 | 0.0944 | 0.8174 | 5 | 417 | 0.1390 | 0.1090 | 1.1990 | 0.4478 | 0.6023 | PRJNA609151 | Permeable sediment | 30 cm | Middle Park Beach, Port Phillip Bay, Australia | | -37.8513 | 144.9544 |
| ERR1811626 | 8 | 179 | 0.0770 | 0.0507 | 4.4693 | 5 | 147 | 0.0580 | 0.0423 | 3.4014 | 0.3917 | 0.4174 | PRJEB19235 | Salt marsh sediment | | Mersea Island, United Kingdom | Natural salt marsh sediment | 51.79717 | 0.920967 |
| SRR8556602 | 4 | 110 | 0.0700 | 0.0087 | 3.6364 | 5 | 113 | 0.1055 | 0.0090 | 4.4248 | 0.0326 | 0.1165 | PRJNA518264 | Salt marsh sediment | 30 cm | Massachusetts, United States | Salt marsh sediment | 42.759 | -70.891 |
| SRR9700688 | 4 | 141 | 0.0778 | 0.0071 | 2.8369 | 5 | 102 | 0.1173 | 0.0052 | 4.9020 | 0.0248 | 0.1129 | PRJNA539485 | Salt marsh sediment | 70 cm | Massachusetts, United States | Salt marsh sediment | 42.722 | -70.847 |
| SRR10156265 | 3 | 150 | 0.0207 | 0.0377 | 2.0000 | 4 | 117 | 0.0332 | 0.0299 | 3.4188 | 0.2045 | 0.2614 | PRJNA573088 | Marine sediment (permanent) | 223 - 820 m | the continental shelf of Antarctica | Wrights Gulf 2 | -73.16 | -129.27 |
| SRR10156273 | 3 | 229 | 0.0227 | 0.0438 | 1.3100 | 4 | 183 | 0.0365 | 0.0356 | 2.1858 | 0.2397 | 0.3134 | PRJNA573088 | Marine sediment (permanent) | 223 - 820 m | the continental shelf of Antarctica | Near Ross Shelf 1 | -76.55 | -170 |
| SRR14844108 | 14 | 822 | 0.0208 | 0.0563 | 1.7032 | 4 | 669 | 0.0072 | 0.0465 | 0.5979 | 0.3316 | 0.3367 | PRJNA724996 | Reef sediment | | Xisha islands, China | | 16.46878 | 111.616 |
| SRR14844111 | 12 | 967 | 0.0361 | 0.0609 | 1.2410 | 4 | 780 | 0.0145 | 0.0499 | 0.5128 | 0.1596 | 0.1703 | PRJNA724996 | Reef sediment | | Xisha islands, China | | 16.97752 | 112.3099 |
| ERR1811650 | 4 | 279 | 0.1366 | 0.0850 | 1.4337 | 4 | 185 | 0.1649 | 0.0572 | 2.1622 | 0.4628 | 0.4598 | PRJEB19235 | Salt marsh sediment | | Fingringhoe Range, United Kingdom | Natural salt marsh sediment | 51.83182 | 0.9687 |
| SRR9710702 | 4 | 60 | 0.0833 | 0.0032 | 6.6667 | 4 | 50 | 0.1006 | 0.0027 | 8.0000 | 0.0140 | 0.1010 | PRJNA539503 | Salt marsh sediment | 80 cm | Massachusetts, United States | Salt marsh sediment | 42.722 | -70.847 |
| ERR1811661 | 2 | 112 | 0.0735 | 0.0453 | 1.7857 | 4 | 123 | 0.1775 | 0.0506 | 3.2520 | 0.2732 | 0.2844 | PRJEB19235 | Salt marsh sediment | | | Sp MIRM 2A | 51.79675 | 0.921833 |
| SRR6394718 | 2 | 96 | 0.0930 | 0.0839 | 2.0833 | 4 | 60 | 0.2245 | 0.0532 | 6.6667 | 0.5876 | 0.4565 | PRJNA4229171 | Salt marsh sediment | | Sydney, Australia | natural saltmarsh | -33.8402 | 151.0588 |
| SRR18461117 | 5 | 318 | 0.0315 | 0.0550 | 1.5723 | 3 | 248 | 0.0228 | 0.0436 | 1.2097 | 0.3810 | 0.4248 | PRJNA819259 | Coastal surface sediments | 18 ~ 15 cm | the Shores of Kuwait | Site 12 | 28.66439 | 48.38989 |
| SRR6207735 | 20 | 3003 | 0.1476 | 0.0699 | 0.6660 | 3 | 2368 | 0.0267 | 0.0560 | 0.1267 | 0.4774 | 0.5501 | PRJNA366768 | Estuary sediment | | White Oak River estuary, North Carolina, United States | | 34.6402 | -77.1094 |
| SRR14844109 | 11 | 777 | 0.0426 | 0.0505 | 1.4157 | 3 | 645 | 0.0140 | 0.0426 | 0.4651 | 0.1401 | 0.1617 | PRJNA724996 | Reef sediment | | Xisha islands, China | | 16.65982 | 112.7551 |
| SRR9710522 | 10 | 252 | 0.1696 | 0.0168 | 3.9683 | 3 | 208 | 0.0614 | 0.0141 | 1.4423 | 0.0617 | 0.1424 | PRJNA539500 | Salt marsh sediment | 70 cm | Massachusetts, United States | Salt marsh sediment | 42.722 | -70.847 |
| ERR1811659 | 6 | 101 | 0.1247 | 0.0446 | 5.9406 | 3 | 75 | 0.0752 | 0.0337 | 4.0000 | 0.4578 | 0.4249 | PRJEB19235 | Salt marsh sediment | | Mersea Island, United Kingdom | Natural salt marsh sediment | 51.79717 | 0.920967 |
| SRR6394731 | 5 | 234 | 0.0293 | 0.0860 | 2.1368 | 3 | 174 | 0.0212 | 0.0650 | 1.7241 | 0.4802 | 0.4243 | PRJNA4229171 | Salt marsh sediment | | Sydney, Australia | regenerated saltmarsh | -33.8402 | 151.0588 |
| SRR8556254 | 5 | 357 | 0.0411 | 0.0212 | 1.4006 | 3 | 297 | 0.0297 | 0.0179 | 1.0101 | 0.0455 | 0.0978 | PRJNA518258 | Salt marsh sediment | 30 cm | Massachusetts, United States | Salt marsh sediment | 42.759 | -70.891 |
| SRR6394725 | 5 | 94 | 0.0450 | 0.0560 | 5.3191 | 3 | 72 | 0.0326 | 0.0436 | 4.1667 | 0.4332 | 0.4414 | PRJNA4229171 | Salt marsh sediment | | Sydney, Australia | regenerated saltmarsh | -33.8402 | 151.0588 |
| ERR1811648 | 4 | 158 | 0.2167 | 0.0914 | 2.5316 | 3 | 114 | 0.1961 | 0.0670 | 2.6316 | 0.4866 | 0.4536 | PRJEB19235 | Salt marsh sediment | | Fingringhoe Range, United Kingdom | Natural salt marsh sediment | 51.83015 | 0.9503 |
| SRR8557366 | 2 | 252 | 0.0767 | 0.0271 | 0.7937 | 3 | 229 | 0.1389 | 0.0251 | 1.3100 | 0.1287 | 0.2095 | PRJNA518280 | Salt marsh sediment | 20 cm | Massachusetts, United States | Salt marsh sediment | 42.759 | -70.891 |
| ERR1811652 | 1 | 274 | 0.0207 | 0.1037 | 0.3650 | 3 | 155 | 0.0748 | 0.0596 | 1.9355 | 0.5386 | 0.5465 | PRJEB19235 | Salt marsh sediment | | Fingringhoe Range, United Kingdom | Realigned salt marsh sediment | 51.83015 | 0.9503 |
| SRR8557033 | 1 | 84 | 0.0242 | 0.0063 | 1.1905 | 3 | 54 | 0.0874 | 0.0041 | 5.5556 | 0.0348 | 0.0918 | PRJNA518278 | Salt marsh sediment | 90 cm | Massachusetts, United States | Salt marsh sediment | 42.759 | -70.891 |
| ERR1811651 | 1 | 144 | 0.0066 | 0.0612 | 0.6944 | 3 | 115 | 0.0241 | 0.0496 | 2.6087 | 0.3573 | 0.4387 | PRJEB19235 | Salt marsh sediment | | Fingringhoe Range, United Kingdom | Realigned salt marsh sediment | 51.83015 | 0.9503 |
| ERR1811628 | 0 | 128 | 0.0000 | 0.0388 | 0.0000 | 3 | 126 | 0.0997 | 0.0388 | 2.3810 | 0.2098 | 0.2294 | PRJEB19235 | Salt marsh sediment | | Mersea Island, United Kingdom | Realigned salt marsh sediment | 51.79675 | 0.921833 |
| SRR8467673 | 4 | 210 | 0.0146 | 0.0599 | 1.9048 | 3 | 173 | 0.0132 | 0.0501 | 1.7341 | 0.5728 | 0.6497 | PRJNA515295 | Subtidal sediment | 2 ~ 5 cm | Middle Park Beach, Melbourne, Australia | | -37.8524 | 144.9575 |
| SRR18461120 | 3 | 231 | 0.0089 | 0.0347 | 1.2987 | 2 | 170 | 0.0071 | 0.0259 | 1.1765 | 0.3980 | 0.5081 | PRJNA819259 | Coastal surface sediments | 21 ~ 15 cm | the Shores of Kuwait | Site 1 | 29.33482 | 47.90238 |
| SRR5248173 | 3 | 46 | 0.0597 | 0.0007 | 6.5217 | 2 | 45 | 0.0480 | 0.0007 | 4.4444 | 0.0041 | 0.4460 | PRJNA366183 | Estuary sediment | | White Oak River estuary, North Carolina, United States | | 34.64781 | -77.1112 |
| SRR5248174 | 3 | 212 | 0.0502 | 0.0036 | 1.4151 | 2 | 192 | 0.0404 | 0.0033 | 1.0417 | 0.0180 | 0.2300 | PRJNA366184 | Estuary sediment | | White Oak River estuary, North Carolina, United States | | 34.6402 | -77.1094 |
| ERR257716 | 3 | 79 | 0.0095 | 0.0636 | 3.7975 | 2 | 59 | 0.0076 | 0.0482 | 3.3898 | 2.0612 | 1.8929 | PRJEB1760 | Intertidal Sediment | 1 cm | Norwich, United Kingdom | Coastal intertidal sediment | 52.96528 | 0.925833 |
| SRR10156269 | 12 | 387 | 0.1987 | 0.0976 | 3.1008 | 2 | 341 | 0.0400 | 0.0874 | 0.5865 | 0.4553 | 0.5070 | PRJNA573088 | Marine sediment (permanent) | 223 - 820 m | the continental shelf of Antarctica | Amundsen Sea 1 | -71.45 | -102.16 |
| SRR10156272 | 1 | 149 | 0.0100 | 0.0250 | 0.6711 | 2 | 124 | 0.0241 | 0.0211 | 1.6129 | 0.1744 | 0.2095 | PRJNA573088 | Marine sediment (permanent) | 223 - 820 m | the continental shelf of Antarctica | Near Ross Shelf 2 | -75.5 | -166.3 |
| ERR1341886 | 6 | 111 | 0.0895 | 0.0955 | 5.4054 | 2 | 63 | 0.0360 | 0.0551 | 3.1746 | 0.4400 | 0.5382 | PRJEB13222 | Reef sediment | | Atlantic Ocean off the coast of Faro, Algarve, Portugal | sediment metagenome | 36.97978 | -7.98911 |
| ERR1811630 | 8 | 222 | 0.4898 | 0.1032 | 3.6036 | 2 | 170 | 0.1478 | 0.0803 | 1.1765 | 0.8770 | 0.6471 | PRJEB19235 | Salt marsh sediment | | Mersea Island, United Kingdom | Natural salt marsh sediment | 51.79717 | 0.920967 |
| SRR5468418 | 7 | 123 | 0.0191 | 0.0227 | 5.6911 | 2 | 98 | 0.0066 | 0.0183 | 2.0408 | 0.3266 | 0.3326 | PRJNA365353 | Salt marsh sediment | 0 | South San Francisco, United States | Tidal salt marsh | 37.496 | -122.133 |
| ERR1811642 | 4 | 244 | 0.0992 | 0.0921 | 1.6393 | 2 | 209 | 0.0599 | 0.0801 | 0.9569 | 0.7465 | 0.7129 | PRJEB19235 | Salt marsh sediment | | Abbotts Hall, United Kingdom | Natural salt marsh sediment | 51.7865 | 0.86085 |
| SRR8557488 | 4 | 259 | 0.0730 | 0.0250 | 1.5444 | 2 | 208 | 0.0440 | 0.0204 | 0.9615 | 0.0994 | 0.1618 | PRJNA518285 | Salt marsh sediment | | the Plum Island Ecosystem LTER, Massachusetts, United States | SW1603-20 | 42.722 | -70.891 |
| SRR8556772 | 4 | 71 | 0.1134 | 0.0059 | 5.6338 | 2 | 48 | 0.0684 | 0.0041 | 4.1667 | 0.0227 | 0.0850 | PRJNA518277 | Salt marsh sediment | 110 cm | Massachusetts, United States | Salt marsh sediment | 42.759 | -70.891 |
| ERR1811649 | 3 | 205 | 0.1528 | 0.0979 | 1.4634 | 2 | 153 | 0.1229 | 0.0742 | 1.3072 | 0.4869 | 0.4684 | PRJEB19235 | Salt marsh sediment | | Fingringhoe Range, United Kingdom | Natural salt marsh sediment | 51.83182 | 0.9687 |
| ERR1811657 | 3 | 208 | 0.0247 | 0.0851 | 1.4423 | 2 | 139 | 0.0198 | 0.0578 | 1.4388 | 0.5750 | 0.6627 | PRJEB19235 | Salt marsh sediment | | Fingringhoe Range, United Kingdom | Realigned salt marsh sediment | 51.83015 | 0.9503 |
| SRR8556703 | 3 | 135 | 0.0279 | 0.0115 | 2.2222 | 2 | 128 | 0.0225 | 0.0111 | 1.5625 | 0.1004 | 0.1599 | PRJNA518274 | Salt marsh sediment | 70 cm | Massachusetts, United States | Salt marsh sediment | 42.759 | -70.891 |
| SRR6394716 | 1 | 12 | 0.0189 | 0.0104 | 8.3333 | 2 | 16 | 0.0456 | 0.0141 | 12.5000 | 0.1734 | 0.2130 | PRJNA4229171 | Salt marsh sediment | | Sydney, Australia | natural saltmarsh | -33.8402 | 151.0588 |
| ERR1811655 | 0 | 85 | 0.0000 | 0.0432 | 0.0000 | 2 | 71 | 0.1758 | 0.0367 | 2.8169 | 0.3347 | 0.2900 | PRJEB19235 | Salt marsh sediment | | Fingringhoe Range, United Kingdom | Natural salt marsh sediment | 51.83182 | 0.9687 |
| SRR6394715 | 0 | 33 | 0.0000 | 0.0399 | 0.0000 | 2 | 43 | 0.1502 | 0.0528 | 4.6512 | 0.4545 | 0.4312 | PRJNA4229171 | Salt marsh sediment | | Sydney, Australia | natural saltmarsh | -33.8402 | 151.0588 |
| SRR8556428 | 0 | 83 | 0.0000 | 0.0053 | 0.0000 | 2 | 74 | 0.0330 | 0.0048 | 2.7027 | 0.0264 | 0.1068 | PRJNA518260 | Salt marsh sediment | 70 cm | Massachusetts, United States | Salt marsh sediment | 42.759 | -70.891 |
| SRR5248164 | 9 | 1279 | 0.1992 | 0.0668 | 0.7037 | 1 | 1082 | 0.0267 | 0.0574 | 0.0924 | 0.4345 | 0.5024 | PRJNA366769 | Estuary sediment | | White Oak River estuary, North Carolina, United States | | 34.6402 | -77.1094 |
| SRR5112543 | 0 | 80 | 0.0000 | 0.0068 | 0.0000 | 1 | 73 | 0.0133 | 0.0063 | 1.3699 | 0.0515 | 0.1829 | PRJNA357337 | Intertidal Sediment | | China | | 37.31 | 122 |
| SRR2844601 | 3 | 108 | 0.1054 | 0.0901 | 2.7778 | 1 | 66 | 0.0424 | 0.0560 | 1.5152 | 0.6745 | 0.6528 | PRJNA298972 | Mangrove sediment | 20 cm | Kerala, India | PYN01 | 12.10507 | 75.2058 |
| SRR2844600 | 1 | 61 | 0.0254 | 0.0586 | 1.6393 | 1 | 52 | 0.0307 | 0.0507 | 1.9231 | 0.3773 | 0.5284 | PRJNA298972 | Mangrove sediment | 20 cm | Kerala, India | MAL01 | 9.90919 | 76.30629 |
| SRR2844602 | 0 | 109 | 0.0000 | 0.0845 | 0.0000 | 1 | 57 | 0.0874 | 0.0449 | 1.7544 | 0.4011 | 0.5846 | PRJNA298972 | Mangrove sediment | 20 cm | Kerala, India | VL101 | 9.999414 | 76.25371 |
| SRR10156271 | 4 | 192 | 0.0487 | 0.0408 | 2.0833 | 1 | 141 | 0.0147 | 0.0304 | 0.7092 | 0.2200 | 0.3231 | PRJNA573088 | Marine sediment (permanent) | 223 - 820 m | the continental shelf of Antarctica | Near Ross Shelf 3 | -74.4 | -168.3 |
| SRR10156274 | 2 | 325 | 0.0306 | 0.1104 | 0.6154 | 1 | 217 | 0.0185 | 0.0749 | 0.4608 | 0.3288 | 0.5557 | PRJNA573088 | Marine sediment (permanent) | 223 - 820 m | the continental shelf of Antarctica | Bellingshausen Sea 2 | -71.42 | -93.48 |
| SRR10156263 | 1 | 138 | 0.0163 | 0.0449 | 0.7246 | 1 | 109 | 0.0197 | 0.0360 | 0.9174 | 0.1798 | 0.2989 | PRJNA573088 | Marine sediment (permanent) | 223 - 820 m | the continental shelf of Antarctica | Ross Sea 2 | -76.2 | -170.51 |
| SRR10156267 | 1 | 155 | 0.0111 | 0.0281 | 0.6452 | 1 | 134 | 0.0135 | 0.0247 | 0.7463 | 0.2161 | 0.3001 | PRJNA573088 | Marine sediment (permanent) | 223 - 820 m | the continental shelf of Antarctica | Amundsen Sea 3 | -72.46 | -104.36 |
| SRR10156275 | 1 | 25 | 0.0484 | 0.0275 | 4.0000 | 1 | 26 | 0.0584 | 0.0291 | 3.8462 | 0.2515 | 0.2984 | PRJNA573088 | Marine sediment (permanent) | 223 - 820 m | the continental shelf of Antarctica | Bellingshausen Sea 1 | -71.42 | -91.43 |
| SRR10156270 | 0 | 211 | 0.0000 | 0.0365 | 0.0000 | 1 | 177 | 0.0112 | 0.0311 | 0.5650 | 0.1991 | 0.3169 | PRJNA573088 | Marine sediment (permanent) | 223 - 820 m | the continental shelf of Antarctica | Bellingshausen Sea 3 | -70.48 | -95.2 |
| SRR9694931 | 6 | 71 | 0.1112 | 0.0044 | 8.4507 | 1 | 72 | 0.0224 | 0.0045 | 1.3889 | 0.0245 | 0.0963 | PRJNA539480 | Salt marsh sediment | 70 cm | Massachusetts, United States | Salt marsh sediment | 42.722 | -70.847 |
| ERR1811654 | 5 | 230 | 0.2168 | 0.1062 | 2.1739 | 1 | 175 | 0.0523 | 0.0821 | 0.5714 | 0.7504 | 0.6177 | PRJEB19235 | Salt marsh sediment | | Fingringhoe Range, United Kingdom | Natural salt marsh sediment | 51.83182 | 0.9687 |
| ERR1811658 | 4 | 221 | 0.1459 | 0.0924 | 1.8100 | 1 | 171 | 0.0440 | 0.0727 | 0.5848 | 0.6010 | 0.5775 | PRJEB19235 | Salt marsh sediment | | Mersea Island, United Kingdom | Natural salt marsh sediment | 51.79717 | 0.920967 |
| ERR1811632 | 3 | 174 | 0.1017 | 0.0787 | 1.7241 | 1 | 108 | 0.0409 | 0.0496 | 0.9259 | 0.4057 | 0.3914 | PRJEB19235 | Salt marsh sediment | | Mersea Island, United Kingdom | Natural salt marsh sediment | 51.79717 | 0.920967 |
| SRR9700687 | 3 | 228 | 0.0189 | 0.0146 | 1.3158 | 1 | 210 | 0.0076 | 0.0137 | 0.4762 | 0.0884 | 0.1461 | PRJNA539482 | Salt marsh sediment | 40 cm | Massachusetts, United States | Salt marsh sediment | 42.722 | -70.847 |
| SRR6394724 | 3 | 68 | 0.0727 | 0.0458 | 4.4118 | 1 | 54 | 0.0292 | 0.0370 | 1.8519 | 0.4351 | 0.4763 | PRJNA4229171 | Salt marsh sediment | | Sydney, Australia | regenerated saltmarsh | -33.8402 | 151.0588 |
| ERR1811633 | 2 | 160 | 0.0404 | 0.0667 | 1.2500 | 1 | 158 | 0.0244 | 0.0669 | 0.6329 | 0.5301 | 0.4996 | PRJEB19235 | Salt marsh sediment | | Mersea Island, United Kingdom | Realigned salt marsh sediment | 51.79675 | 0.921833 |
| SRR9700623 | 2 | 59 | 0.0965 | 0.0052 | 3.3898 | 1 | 56 | 0.0582 | 0.0050 | 1.7857 | 0.0184 | 0.0875 | PRJNA539484 | Salt marsh sediment | 30 cm | Massachusetts, United States | Salt marsh sediment | 42.722 | -70.847 |
| SRR9707989 | 2 | 70 | 0.0390 | 0.0022 | 2.8571 | 1 | 56 | 0.0235 | 0.0018 | 1.7857 | 0.0087 | 0.0805 | PRJNA539497 | Salt marsh sediment | 200 cm | Massachusetts, United States | Salt marsh sediment | 42.722 | -70.847 |
| SRR9710561 | 2 | 46 | 0.0990 | 0.0033 | 4.3478 | 1 | 41 | 0.0597 | 0.0030 | 2.4390 | 0.0120 | 0.1228 | PRJNA539502 | Salt marsh sediment | 70 cm | Massachusetts, United States | Salt marsh sediment | 42.759 | -70.847 |
| ERR1811624 | 1 | 208 | 0.1096 | 0.1434 | 0.4808 | 1 | 153 | 0.1323 | 0.1071 | 0.6536 | 0.9449 | 0.8349 | PRJEB19235 | Salt marsh sediment | | Mersea Island, United Kingdom | Natural salt marsh sediment | 51.79717 | 0.920967 |
| ERR1811634 | 1 | 42 | 0.0805 | 0.0170 | 2.3810 | 1 | 35 | 0.0971 | 0.0144 | 2.8571 | 0.0537 | 0.1500 | PRJEB19235 | Salt marsh sediment | | Mersea Island, United Kingdom | Realigned salt marsh sediment | 51.79675 | 0.921833 |
| ERR1811635 | 1 | 119 | 0.0466 | 0.0619 | 0.8403 | 1 | 108 | 0.0562 | 0.0571 | 0.9259 | 0.3595 | 0.4298 | PRJEB19235 | Salt marsh sediment | | Mersea Island, United Kingdom | Realigned salt marsh sediment | 51.79675 | 0.921833 |
| SRR9705735 | 1 | 42 | 0.0085 | 0.0018 | 2.3810 | 1 | 44 | 0.0103 | 0.0019 | 2.2727 | 0.0191 | 0.0854 | PRJNA539491 | Salt marsh sediment | 210 cm | Massachusetts, United States | Salt marsh sediment | 42.722 | -70.847 |
| SRR9705674 | 0 | 37 | 0.0000 | 0.0020 | 0.0000 | 1 | 29 | 0.0332 | 0.0016 | 3.4483 | 0.0113 | 0.0952 | PRJNA539489 | Salt marsh sediment | 190 cm | Massachusetts, United States | Salt marsh sediment | 42.722 | -70.847 |
| SRR9705733 | 0 | 13 | 0.0000 | 0.0008 | 0.0000 | 1 | 12 | 0.0234 | 0.0008 | 8.3333 | 0.0069 | 0.0907 | PRJNA539492 | Salt marsh sediment | 130 cm | Massachusetts, United States | Salt marsh sediment | 42.722 | -70.847 |
| SRR9707468 | 0 | 4 | 0.0000 | 0.0002 | 0.0000 | 1 | 8 | 0.0608 | 0.0004 | 12.5000 | 0.0021 | 0.0747 | PRJNA539496 | Salt marsh sediment | 110 cm | Massachusetts, United States | Salt marsh sediment | 42.722 | -70.847 |
| SRR6195247 | 2 | 268 | 0.0264 | 0.0439 | 0.7463 | 1 | 233 | 0.0159 | 0.0387 | 0.4292 | 0.2773 | 0.5041 | PRJNA366574 | Salt pond sediment | 32 cm | San Francisco, United States | Eden Landing Ponds | 37.56908 | -122.103 |
| SRR16005299 | 10 | 2007 | 0.0522 | 0.1165 | 0.4983 | 0 | 1437 | 0.0000 | 0.0847 | 0.0000 | 0.5689 | 0.7168 | PRJNA765123 | Estuary sediment | 0 ~ 1 cm | Yangtze Estuary, China | Win_13C_meta | 31 | 122 |
| SRR5248172 | 1 | 214 | 0.0211 | 0.0037 | 0.4673 | 0 | 184 | 0.0000 | 0.0032 | 0.0000 | 0.0161 | 0.3009 | PRJNA366355 | Estuary sediment | | White Oak River estuary, North Carolina, United States | | 34.6402 | -77.1094 |
| ERR257713 | 4 | 34 | 0.1599 | 0.0497 | 11.7647 | 0 | 23 | 0.0000 | 0.0342 | 0.0000 | 2.0306 | 2.6615 | PRJEB1760 | Intertidal Sediment | 1 cm | Norwich, United Kingdom | Coastal intertidal sediment | 52.96528 | 0.925833 |
| SRR2134644 | 4 | 202 | 0.1879 | 0.0535 | 1.9802 | 0 | 165 | 0.0000 | 0.0444 | 0.0000 | 0.2723 | 0.3492 | PRJNA283251 | Intertidal Sediment | 0 ~ 10 cm | Hong Kong | | 22.43833 | 114.2167 |
| SRR5112544 | 2 | 157 | 0.0098 | 0.0109 | 1.2739 | 0 | 129 | 0.0000 | 0.0091 | 0.0000 | 0.0986 | 0.2798 | PRJNA357337 | Intertidal Sediment | | China | | 37.31 | 122 |
| SRR5112542 | 0 | 69 | 0.0000 | 0.0069 | 0.0000 | 0 | 61 | 0.0000 | 0.0062 | 0.0000 | 0.0374 | 0.2061 | PRJNA357337 | Intertidal Sediment | | China | | 37.31 | 122 |
| SRR2844616 | 0 | 21 | 0.0000 | 0.0176 | 0.0000 | 0 | 13 | 0.0000 | 0.0111 | 0.0000 | 0.3466 | 0.4492 | PRJNA298972 | Mangrove sediment | 20 cm | Kerala, India | PGD01 | 9.895994 | 76.32609 |
| SRR3620785 | 0 | 4 | 0.0000 | 0.0015 | 0.0000 | 0 | 1 | 0.0000 | 0.0004 | 0.0000 | 0.1740 | 0.1922 | PRJNA321092 | Mangrove sediment | | Rabigh coast in Saudi Arabia | MNR (mangrove mud from Rabigh harbor lagoon) | 22.75155 | 39.00993 |
| SRR10156266 | 1 | 184 | 0.0126 | 0.0530 | 0.5435 | 0 | 146 | 0.0000 | 0.0427 | 0.0000 | 0.3151 | 0.4165 | PRJNA573088 | Marine sediment (permanent) | 223 - 820 m | the continental shelf of Antarctica | Wrights Gulf 1 | -73.3 | -129.56 |
| SRR3620783 | 0 | 23 | 0.0000 | 0.0028 | 0.0000 | 0 | 21 | 0.0000 | 0.0026 | 0.0000 | 0.1280 | 0.2212 | PRJNA321092 | Microbial mat | | Rabigh coast in Saudi Arabia | MMR (microbial mat from Rabigh harbor lagoon) | 22.75155 | 39.00993 |
| SRR8556257 | 5 | 405 | 0.0622 | 0.0464 | 1.2346 | 0 | 355 | 0.0000 | 0.0414 | 0.0000 | 0.2200 | 0.2804 | PRJNA518257 | Salt marsh sediment | 20 | Massachusetts, United States | Salt marsh sediment | 42.759 | -70.891 |
| ERR1811644 | 5 | 417 | 0.1961 | 0.1372 | 1.1990 | 0 | 271 | 0.0000 | 0.0906 | 0.0000 | 0.7971 | 0.5753 | PRJEB19235 | Salt marsh sediment | | Abbotts Hall, United Kingdom | Natural salt marsh sediment | 51.7865 | 0.86085 |
| SRR9705675 | 5 | 207 | 0.1214 | 0.0168 | 2.4155 | 0 | 152 | 0.0000 | 0.0125 | 0.0000 | 0.0731 | 0.1603 | PRJNA539490 | Salt marsh sediment | 20 cm | Massachusetts, United States | Salt marsh sediment | 42.722 | -70.847 |
| ERR1811627 | 4 | 69 | 0.1379 | 0.0326 | 5.7971 | 0 | 69 | 0.0000 | 0.0331 | 0.0000 | 0.1868 | 0.2801 | PRJEB19235 | Salt marsh sediment | | Mersea Island, United Kingdom | Realigned salt marsh sediment | 51.79675 | 0.921833 |
| ERR1811637 | 4 | 89 | 0.2833 | 0.0862 | 4.4944 | 0 | 58 | 0.0000 | 0.0571 | 0.0000 | 0.4410 | 0.3925 | PRJEB19235 | Salt marsh sediment | | Abbotts Hall, United Kingdom | Natural salt marsh sediment | 51.7865 | 0.86085 |
| SRR6394726 | 2 | 101 | 0.0287 | 0.0673 | 1.9802 | 0 | 69 | 0.0000 | 0.0467 | 0.0000 | 0.6442 | 0.6017 | PRJNA4229171 | Salt marsh sediment | | Sydney, Australia | regenerated saltmarsh | -33.8402 | 151.0588 |
| SRR9700529 | 2 | 32 | 0.1088 | 0.0030 | 6.2500 | 0 | 17 | 0.0000 | 0.0016 | 0.0000 | 0.0074 | 0.1249 | PRJNA539486 | Salt marsh sediment | 230 cm | Massachusetts, United States | Salt marsh sediment | 42.722 | -70.847 |
| ERR1811625 | 1 | 94 | 0.0479 | 0.0549 | 1.0638 | 0 | 78 | 0.0000 | 0.0463 | 0.0000 | 0.3737 | 0.3399 | PRJEB19235 | Salt marsh sediment | | Mersea Island, United Kingdom | Natural salt marsh sediment | 51.79717 | 0.920967 |
| ERR1811631 | 1 | 116 | 0.0581 | 0.0691 | 0.8621 | 0 | 110 | 0.0000 | 0.0666 | 0.0000 | 0.4517 | 0.4060 | PRJEB19235 | Salt marsh sediment | | Mersea Island, United Kingdom | Natural salt marsh sediment | 51.79717 | 0.920967 |
| ERR1811645 | 1 | 4 | 0.0516 | 0.0033 | 25.0000 | 0 | 1 | 0.0000 | 0.0008 | 0.0000 | 0.2187 | 0.2119 | PRJEB19235 | Salt marsh sediment | | | W AHRM 1A | 51.78783 | 0.855517 |
| SRR6394721 | 1 | 15 | 0.0292 | 0.0228 | 6.6667 | 0 | 9 | 0.0000 | 0.0139 | 0.0000 | 0.3244 | 0.2376 | PRJNA4229171 | Salt marsh sediment | | Sydney, Australia | natural saltmarsh | -33.8402 | 151.0588 |
| SRR8556701 | 1 | 27 | 0.0831 | 0.0018 | 3.7037 | 0 | 19 | 0.0000 | 0.0013 | 0.0000 | 0.0038 | 0.0984 | PRJNA518270 | Salt marsh sediment | 170 cm | Massachusetts, United States | Salt marsh sediment | 42.759 | -70.891 |
| SRR9700624 | 1 | 38 | 0.0181 | 0.0023 | 2.6316 | 0 | 25 | 0.0000 | 0.0015 | 0.0000 | 0.0142 | 0.0903 | PRJNA539481 | Salt marsh sediment | 30 cm | Massachusetts, United States | Salt marsh sediment | 42.722 | -70.847 |
| SRR9700625 | 1 | 7 | 0.0561 | 0.0004 | 14.2857 | 0 | 12 | 0.0000 | 0.0008 | 0.0000 | 0.0011 | 0.0933 | PRJNA539487 | Salt marsh sediment | 150 cm | Massachusetts, United States | Salt marsh sediment | 42.722 | -70.847 |
| ERR1811629 | 0 | 13 | 0.0000 | 0.0129 | 0.0000 | 0 | 7 | 0.0000 | 0.0071 | 0.0000 | 0.0484 | 0.2168 | PRJEB19235 | Salt marsh sediment | | Mersea Island, United Kingdom | Realigned salt marsh sediment | 51.79675 | 0.921833 |
| ERR1811639 | 0 | 14 | 0.0000 | 0.0167 | 0.0000 | 0 | 6 | 0.0000 | 0.0073 | 0.0000 | 0.3091 | 0.3255 | PRJEB19235 | Salt marsh sediment | 5 cm | | S AHRM 1A | 51.78783 | 0.855517 |
| ERR1811640 | 0 | 41 | 0.0000 | 0.0213 | 0.0000 | 0 | 25 | 0.0000 | 0.0132 | 0.0000 | 0.2190 | 0.2756 | PRJEB19235 | Salt marsh sediment | | | S AHRM 2B | 51.78783 | 0.855517 |
| ERR1811641 | 0 | 70 | 0.0000 | 0.0362 | 0.0000 | 0 | 50 | 0.0000 | 0.0263 | 0.0000 | 0.2468 | 0.3144 | PRJEB19235 | Salt marsh sediment | | | S AHRM 3B | 51.78783 | 0.855517 |
| ERR1811646 | 0 | 11 | 0.0000 | 0.0067 | 0.0000 | 0 | 5 | 0.0000 | 0.0031 | 0.0000 | 0.2223 | 0.2543 | PRJEB19235 | Salt marsh sediment | | | W AHRM 2A | 51.78783 | 0.855517 |
| ERR1811647 | 0 | 45 | 0.0000 | 0.0226 | 0.0000 | 0 | 39 | 0.0000 | 0.0199 | 0.0000 | 0.2323 | 0.2975 | PRJEB19235 | Salt marsh sediment | | Abbotts Hall, United Kingdom | Realigned salt marsh sediment | 51.78783 | 0.855517 |
| ERR1811660 | 0 | 2 | 0.0000 | 0.0024 | 0.0000 | 0 | 1 | 0.0000 | 0.0012 | 0.0000 | 0.2121 | 0.2101 | PRJEB19235 | Salt marsh sediment | | Mersea Island, United Kingdom | Realigned salt marsh sediment | 51.79675 | 0.921833 |
| SRR6394717 | 0 | 41 | 0.0000 | 0.0639 | 0.0000 | 0 | 29 | 0.0000 | 0.0459 | 0.0000 | 0.3215 | 0.3537 | PRJNA4229171 | Salt marsh sediment | | Sydney, Australia | natural saltmarsh | -33.8402 | 151.0588 |
| SRR6394722 | 0 | 2 | 0.0000 | 0.0007 | 0.0000 | 0 | 2 | 0.0000 | 0.0007 | 0.0000 | 0.2268 | 0.1767 | PRJNA4229171 | Salt marsh sediment | | Sydney, Australia | natural saltmarsh | -33.8402 | 151.0588 |
| SRR6394727 | 0 | 62 | 0.0000 | 0.0477 | 0.0000 | 0 | 58 | 0.0000 | 0.0454 | 0.0000 | 0.4719 | 0.4896 | PRJNA4229171 | Salt marsh sediment | | Sydney, Australia | regenerated saltmarsh | -33.8402 | 151.0588 |
| SRR8556479 | 0 | 95 | 0.0000 | 0.0290 | 0.0000 | 0 | 58 | 0.0000 | 0.0180 | 0.0000 | 0.0711 | 0.1084 | PRJNA518261 | Salt marsh sediment | 90 cm | Massachusetts, United States | Salt marsh sediment | 42.759 | -70.891 |
| SRR8556606 | 0 | 8 | 0.0000 | 0.0006 | 0.0000 | 0 | 17 | 0.0000 | 0.0013 | 0.0000 | 0.0010 | 0.1005 | PRJNA518266 | Salt marsh sediment | 190 cm | Massachusetts, United States | Salt marsh sediment | 42.759 | -70.891 |
| SRR8556697 | 0 | 32 | 0.0000 | 0.0022 | 0.0000 | 0 | 13 | 0.0000 | 0.0009 | 0.0000 | 0.0046 | 0.0712 | PRJNA518269 | Salt marsh sediment | 130 cm | Massachusetts, United States | Salt marsh sediment | 42.759 | -70.891 |
| SRR8556769 | 0 | 24 | 0.0000 | 0.0017 | 0.0000 | 0 | 21 | 0.0000 | 0.0015 | 0.0000 | 0.0088 | 0.0972 | PRJNA518276 | Salt marsh sediment | 90 cm | Massachusetts, United States | Salt marsh sediment | 42.759 | -70.891 |
| SRR8557270 | 0 | 12 | 0.0000 | 0.0010 | 0.0000 | 0 | 10 | 0.0000 | 0.0008 | 0.0000 | 0.0055 | 0.0782 | PRJNA518282 | Salt marsh sediment | 220 cm | Massachusetts, United States | Salt marsh sediment | 42.722 | -70.891 |
| SRR8557364 | 0 | 176 | 0.0000 | 0.0120 | 0.0000 | 0 | 132 | 0.0000 | 0.0092 | 0.0000 | 0.0192 | 0.0937 | PRJNA518284 | Salt marsh sediment | 30 cm | Massachusetts, United States | Salt marsh sediment | 42.722 | -70.891 |
| SRR9705745 | 0 | 5 | 0.0000 | 0.0004 | 0.0000 | 0 | 5 | 0.0000 | 0.0005 | 0.0000 | 0.0057 | 0.1121 | PRJNA539494 | Salt marsh sediment | 170 cm | Massachusetts, United States | Salt marsh sediment | 42.722 | -70.847 |
| SRR9706179 | 0 | 25 | 0.0000 | 0.0009 | 0.0000 | 0 | 13 | 0.0000 | 0.0005 | 0.0000 | 0.0047 | 0.0810 | PRJNA539493 | Salt marsh sediment | 10 cm | Massachusetts, United States | Salt marsh sediment | 42.722 | -70.847 |
| SRR8556427 | 0 | 49 | 0.0000 | 0.0038 | 0.0000 | 0 | 61 | 0.0000 | 0.0048 | 0.0000 | 0.0327 | 0.0920 | PRJNA518259 | Salt marsh sediment | 40 cm | Massachusetts, United States | Salt marsh sediment | 42.759 | -70.891 |

**Table S7. Quantitative PCR primers**

| Gene | Locus id | Forward primer (5’ to 3’) | Reverse primer (5’ to 3’) |
|---|---|---|---|
| *narG* | SZCCDBA067_01139 | GACCCTGTGTCTGGATCAGC | TGTGGGTTGGTTTCAGGACC |
| *nirS* | SZCCDBA067_00039 | GAGCCCCACTGGAAGAAGAC | CGGCTTCCTCGACATAGACC |
| *nosZ* | SZCCDBA067_00053 | CTGTTCCTCGACAGCCAGG | ATCGAGATCAGCCACTTGCC |
| *ccoN* | SZCCDBA067_02896 | TCCAAGGAATATGCCGAGCC | ACCAGTTGGCGACGTAGATG |
| *ctaB* | SZCCDBA067_00252 | CATCGTCTGGGTCGCCTATC | ACGCCAGGTAGAACCAGTTG |
| *dsrA* | SZCCDBA067_02173 | GACATCATGTTCCAGGG | CGTCAAAGCAGGACTG |
| *dsrB* | SZCCDBA067_02172 | TGGCTGCATTGCGACATTCCC | GGTTGATCTTGGGCGGCTTGGTGTA |
| *dsrJ* | SZCCDBA067_02165 | GCAACTTCGAGGAACGCATC | GAAGATCACGCAGTTGTCGC |
| *sqr* | SZCCDBA067_02259 | GTCAAGGTGATCTGCAACCG | TTCTCCGGGGTCATCGTCT |
| *pdo* | SZCCDBA067_02258 | GTGTCCTATCTTGTCCGCGA | CTTTCGAGGACCCACTCCAC |
| *sqr_2* | SZCCDBA067_01474 | ATCAAGTGGATCACCTCGGC | TACCAACCCTTCGATGCCAG |
| *Rhodanese_2* | SZCCDBA067_01274 | CATCCAAGGCGTCAATGCTG | TTGTGGAAATCGGGATGCG |
| *soeA* | SZCCDBA067_02936 | CACAGCAGCCGAGGATAGAG | TTCATGTGCACGTTGATGCC |
| *rpoB* | SZCCDBA067_02879 | ACCTATGACCTGATCGACGC | GTCCTTGTCGTATTCCAGCG |
| *01184* | SZCCDBA067_01184 | TGGCGCAAACTATCACGAAGG | GAAAATGCTCCGGGGATCTGG |
| *03117* | SZCCDBA067_03117 | ATCTGCGGATTCTGGATGGG | CATGAACTTTTCGACCGGGG |
| *00803* | SZCCDBA067_00803 | ATCTGCGGATTCTGGATGGG | CATGAACTTTTCGACCGGGG |
| *02466* | SZCCDBA067_02466 | ACGGTCGAACTTCACGATCC | CCCGGAATGTGACCCGAATC |
| *00795* | SZCCDBA067_00795 | TCATTCTGCGCCCTTGATCC | TCTTCAAACGCTCGGCATCC |
| *00756* | SZCCDBA067_00756 | GTCGGGTCGCATCATATCCC | CGCCTAGATTGAAGACTTGCG |

Table S8. Ruegeria strains used in this study.

| Name | Features | Source |
|---|---|---|
| *E. coli DH5α* | Wild type | Takara |
| *R. pomeroyi* | Wild type strain (DSM15171) | China General Microbiological |
| SZCCDBA06 | Wild type strain | This study |
| DSS-3-1 | *R. pomeroyi DSS-3 Wild type, pcaF* | This study |
| DSS-3-2 | *R. pomeroyi DSS-3 Wild type, pcaF* | This study |
| DSS-3-3 | *R. pomeroyi DSS-3 Wild type, pcaF* | This study |
| DSS-3-4 | *R. pomeroyi DSS-3 Wild type, pcaF* | This study |
| DSS-3-5 | *R. pomeroyi DSS-3 Wild type, soxB* | This study |
| A067-1 | SZCCDBA067 Wild type, rhd | This study |
| A067-2 | SZCCDBA067 Wild type, dsrA | This study |

Table S9. Plasmids used in this study.

| Name | Description | Source |
|---|---|---|
| pTemplate | pUC ori, bla, gRNA scaffold | Lab stock |
| pBE | pBBR1MCS-5, lacI-Ptrc, | Lab stock |
| pgRNA-pcaF1 | pTemplate, gRNA01 | This study |
| pgRNA-pcaF2 | pTemplate, gRNA02 | This study |
| pgRNA-pcaF3 | pTemplate, gRNA03 | This study |
| pgRNA-pcaF4 | pTemplate, gRNA04 | This study |
| pgRNA-rhd | pTemplate, gRNA05 | This study |
| pgRNA-dsrA | pTemplate, gRNA06 | This study |
| pBE-pcaF1 | pBE, gRNA01 | This study |
| pBE-pcaF2 | pBE, gRNA02 | This study |
| pBE-pcaF3 | pBE, gRNA03 | This study |
| pBE-pcaF4 | pBE, gRNA04 | This study |
| pBE-rhd | pBE, gRNA05 | This study |
| pBE-dsrA | pBE, gRNA06 | This study |

Table S10. Primers used in this study.

| Name | Description |
|---|---|
| **Primers for** | **DNA assembly** |
| XIA-ZWQ-01 | ACCGGAAGCAGTGTTCTAGATTGTAAAACGACGGCCAGTC |
| XIA-ZWQ-02 | ATTTGAGAAGCACACGGTCACAGGAAACAGCTATGACCGT |
| XIA-ZWQ-03 | GACTGGCCGTCGTTTTACAATCTAGAACACTGCTTCCGGT |
| XIA-ZWQ-04 | ACGGTCATAGCTGTTTCCTGTGACCGTGTGCTTCTCAAAT |
| **Primers for inverse PCR** | |
| XIA-ZWQ-05 | GCCAGGGGCCGTCCTCCAGCGTTTTAGAGCTAGAAATAGC |
| XIA-ZWQ-06 | GCTGGAGGACGGCCCCTGGCGCTAGCATTATACCTAGGAC |
| XIA-ZWQ-07 | CCAAGGCGTCAATGCTGGCAGTTTTAGAGCTAGAAATAGC |
| XIA-ZWQ-08 | TGCCAGCATTGACGCCTTGGGCTAGCATTATACCTAGGAC |
| XIA-ZWQ-09 | GCGGTGTCGAAAGCATGAGCGTTTTAGAGCTAGAAATAGC |
| XIA-ZWQ-10 | GCTCATGCTTTCGACACCGCGCTAGCATTATACCTAGGAC |
| XIA-ZWQ-11 | CCAGGTGCGCACCGACGATCGTTTTAGAGCTAGAAATAGC |
| XIA-ZWQ-12 | GATCGTCGGTGCGCACCTGGGCTAGCATTATACCTAGGAC |
| XIA-ZWQ-13 | GCGCGAGATATCGTAATCCTGTTTTAGAGCTAGAAATAGC |
| XIA-ZWQ-14 | AGGATTACGATATCTCGCGCGCTAGCATTATACCTAGGAC |
| XIA-ZWQ-15 | GCGGTCTATGACACCACCATGTTTTAGAGCTAGAAATAGC |
| XIA-ZWQ-16 | ATGGTGGTGTCATAGACCGCGCTAGCATTATACCTAGGAC |
| XIA-ZWQ-17 | GGGCAGATGCCCCATATCACGTTTTAGAGCTAGAAATAGC |
| XIA-ZWQ-18 | GTGATATGGGGCATCTGCCCGCTAGCATTATACCTAGGAC |
| **Primers for sequencing** | |
| XIA-ZWQ-19 | GCGTAACAATGCCAGAATCT |
| XIA-ZWQ-20 | TGATTTGGTGATAGTTCCCA |
| XIA-ZWQ-21 | GTTCACGCCCTTGAGCTTGC |
| XIA-ZWQ-22 | TCATGATCTCCCGTCGCGAT |

Table S11. gRNA sequences used in this study.

| gRNA | Target | PAM | Stranda | spacer |
|---|---|---|---|---|
| gRNA-pcaF1 | *pcaF* | TGG | C | CCAGGTGCGCACCGACGATC |
| gRNA-pcaF2 | *pcaF* | CGG | C | GCGGTGTCGAAAGCATGAGC |
| gRNA-pcaF3 | *pcaF* | CGG | C | GCGGTCTATGACACCACCAT |
| gRNA-pcaF4 | *pcaF* | CGG | C | GCGCGAGATATCGTAATCCT |
| gRNA-soxB | *soxB* | CGG | N | GGGCAGATGCCCCATATCAC |
| gRNA-rhd | *rhd* | AGG | C | CCAAGGCGTCAATGCTGGCA |
| gRNA-dsrA | *dsrA* | TGG | N | GCCAGGGGCCGTCCTCCAGC |

a C stands for coding strand and N stands for non-coding strand.
